# 4D hybrid model interrogates agent-level rules and parameters driving hiPS cell colony dynamics

**DOI:** 10.1101/2024.08.12.607546

**Authors:** Jessica S. Yu, Blair Lyons, Susanne Rafelski, Julie A. Theriot, Neda Bagheri, Graham T. Johnson

## Abstract

Iterating between data-driven research and generative computational models is a powerful approach for emulating biological systems, testing hypotheses, and gaining a deeper understanding of these systems. We developed a hybrid agent-based model (ABM) that integrates a Cellular Potts Model (CPM) designed to investigate cell shape and colony dynamics in human induced pluripotent stem cell (hiPS cell) colonies. This model aimed to first mimic and then explore the dynamics observed in real-world hiPS cell cultures.

Initial outputs showed great potential, seeming to mimic small colony behaviors relatively well. However, longer simulations and quantitative comparisons revealed limitations, particularly with the CPM component, which lacked long-range interactions that might be necessary for accurate simulations. This challenge led us to thoroughly examine the hybrid model’s potential and limitations, providing insights and recommendations for systems where cell-wide mechanics play significant roles.

The CPM supports 2D and 3D cell shapes using a Monte Carlo algorithm to prevent cell fragmentation. Basic “out of the box” CPM Hamiltonian terms of volume and adhesion were insufficient to match live cell imaging of hiPS cell cultures. Adding substrate adhesion resulted in flatter colonies, highlighting the need to consider environmental context in modeling. High-throughput parameter sweeps identified regimes that produced consistent simulated shapes and demonstrated the impact of specific model decisions on emergent dynamics. Full-scale simulations showed that while certain agent rules could form a hiPS cell monolayer in 3D, they could not maintain it over time.

Our study underscores that “out of the box” 3D CPMs, which do not natively incorporate long-range cell mechanics like elasticity, may be insufficient for accurately simulating hiPS cell and colony dynamics. To address this limitation, future work could add mechanical constraints to the CPM Hamiltonian or integrate global agent rules. Alternatively, replacing the CPM with a methodology that directly represents cell mechanics might be necessary.

Documenting and sharing our model development process fosters open team science and supports the broader research community in developing computational models of complex biological systems.

## Introduction

Computational models are powerful tools for understanding, emulating, and controlling biological systems [1, 2]. In basic science, these models have helped to decipher the mechanisms behind diverse processes ranging from circadian rhythms [3], action potentials [4], and gene regulation [5] to morphogenesis [6], angiogenesis [7], and regeneration [8]. In translational science, they guide the development of strategies for targeted biological responses, essential in model-guided drug discovery [9], treatment design [10], and personalized medicine [11]. As these computational models become increasingly complex, emulation via machine learning techniques can be used to feasibly enable high resolution, high-throughput multi-scale and multi-class modeling of complex biological systems [12, 13].

In each scenario, the process of developing these models is as crucial as the models themselves; it directly influences the science and its practical applications. Sharing and reflecting on a model development process enables future model development by providing a deeper understanding of the strengths and limitations of existing models and revealing opportunities for integration and extension using new data and methods. *Iterative model development* involves close collaboration with subject matter experts, enhancing the quality and speed of biological insights. This approach allows for the modification of model rules or parameters based on expert feedback, facilitating a deeper exploration of how these changes align—or do not align—with observed biological phenomena [14]. *Data-driven model development* uses direct observations to guide and validate models, grounding abstract concepts like parameters and states in biologically meaningful interpretations. Observed data is used to directly inform parameter values, constrain parameter spaces, and compare computational predictions with real-world emergent biological outcomes. Our research focuses on developing computational models to understand the factors driving cell shape and colony dynamics in human induced pluripotent stem cell (hiPS cell) epithelial-like colonies grown in culture as monolayers on 2D substrates (such as Matrigel coated glass). By integrating an agent-based model (ABM) to capture cell behavior with a Cellular Potts model (CPM) to capture cell shape, and by leveraging subject matter expertise and live 3D microscopy data, we aimed to develop a hybrid model to elucidate how computational rules and parameters result in emergent behaviors that are consistent with those observed in cultured hiPS cells.

ABMs are a flexible, bottom-up modeling methodology well suited for characterizing population-scale dynamics that emerge from cell-scale and subcellular-scale interactions. In these models, agents (representing cells) follow rules (implicit representations of multiple scales of biological mechanisms) that guide their actions and interactions with other agents and with their environment [15]. ABMs have proven effective across various biological contexts, including cancer biology [16, 17, 18], immunology [19, 20], developmental biology [21, 22], and microbiology [23]. In particular, ABMs are well suited for integrating other modeling approaches to enable hybrid modeling.

CPMs are a grid-based modeling methodology well suited for characterizing cell-scale dynamics that emerge from subcellular-scale interactions. In these models, cells are mapped onto a shared grid of voxels and evolve via the minimization of a voxel-level Hamiltonian energy function [24]. Originally developed to study cell sorting [25], CPMs have been extended to study a variety of phenomena including tumor growth [26, 27, 28], cell movement [29], tissue morphogenesis [30], and wound healing [31]. Although CPMs are most commonly used in 2D, often representing real-world 3D systems as a 2D slice, studies have shown notable differences between biological systems studied in 2D versus 3D [32, 33, 34]. There are efforts toward making large-scale 3D CPMs computationally effective [35].

Here specifically, we apply an iterative and data-driven approach to develop a hybrid agent-based model (ABM) that integrates a Cellular Potts Model (CPM). This hybrid model aims to investigate theoretical rules that might drive cell shape and colony dynamics in hiPS cell colonies. We use this model to simulate the formation and growth of hiPS cell colonies in 3D, exploring how underlying agent rules influence 3D cell shape and colony dynamics in simulated hiPS cells, both with and without a simulated 2D culture substrate intended to emulate the Matrigel used in the lab.

## Results

### Hybrid agent-based and Cellular Potts model integrates 3D hiPS cell image data, behavior, and shape dynamics

The hybrid model combines an ABM driving cell behavior with a CPM representing cell shape within the ARCADE modeling framework [36, 18, 14] (Figure 1A).

**Figure 1.**
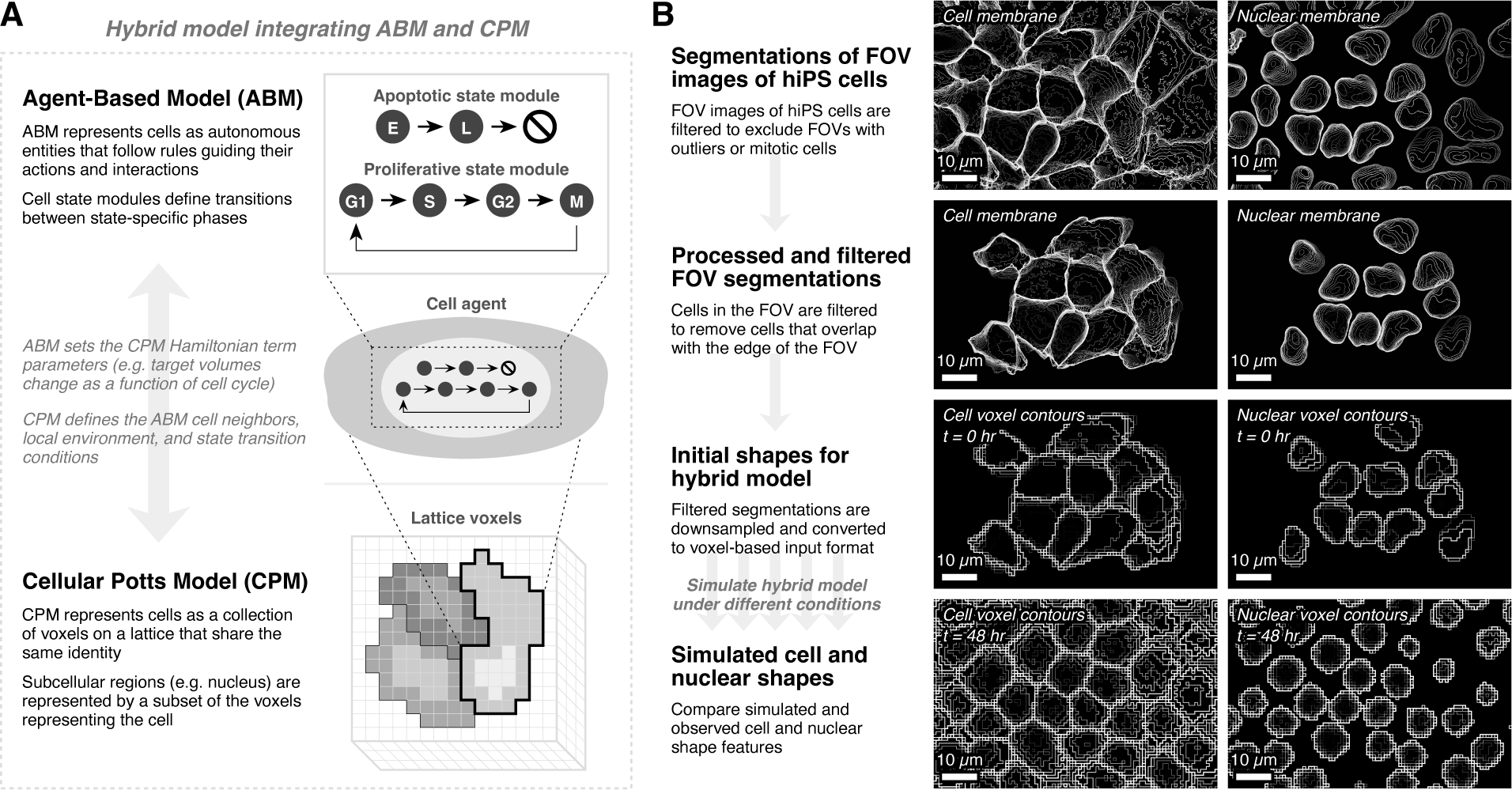
Integration of an ABM (for cell behavior) with a CPM (for cell shape) enables simulation of hiPS cell colonies. (**A**) Overview of hybrid model that integrates between the agent-based model (ABM) and Cellular Potts Model (CPM). Agents represent cells, which can be proliferative or apoptotic. Proliferative cells progress through cell cycle phases: G1 to S to G2 to M. Apoptotic cells progress through early (E) and late (L) apoptosis, before being removed from the simulation. (**B**) Overview of simulation and analysis workflow. Segmented field of view (FOV) images are selected from the WTC-11 hiPSC Single-Cell Image Dataset v1 [37]. These images are filtered to exclude FOVs containing cells marked as outliers and mitotic cells. Cells in an FOV are filtered to remove cells that cross the edge of the FOV. These filtered segmentations are downsampled to the spatial resolution of the model (from 0.108 *µm*/voxel to 1 or 2 *µm*/voxel) and converted to the voxel-based input format used by the hybrid model. Models are then simulated under different agent rules and parameter conditions to produce simulated cell and nuclear shapes that are further analyzed by comparing to the observed data.

The ABM defines cell agent states and checks for transitions between states and state-specific phases. While the ABM can be used to define complex behavioral and state transition rules, we deliberately utilize a very simple set of agent rules in which the majority of cells maintain a proliferative state, progressing through the cell cycle and occasionally becoming apoptotic.

The CPM defines simply connected cell shapes and applies changes using the Hamiltonian energy function. The basic Hamiltonian consists of two terms: a volume constraint term and an adhesion contact energy term. The volume constraint term keeps the cell volume near a specified target volume, while the adhesion contact energy term quantifies contact energies between different types of voxels. The Hamiltonian is flexible for adding in additional terms, such as a differential substrate adhesion contact energy term that specifies the contact energy between cell voxels and substrate voxels. The CPM can be adapted to represent subcellular regions; we specifically define nuclear voxels as a subset of the voxels representing the cell and focus on 3D simulations throughout this study.

The hybrid model is simulated by updating both models at each tick of the simulation and synchronizing the relevant parameters. Each cell agent in the ABM updates relevant Hamiltonian parameters in the CPM, such as target volume which changes as a function of the cell cycle phase. In the other direction, the CPM defines current cell volumes, shapes, and neighbors which impact state transition conditions in the ABM.

By coupling the ABM and the CPM into a hybrid model, we take advantage of the strengths of the two modeling methodologies while also providing flexibility for swapping in different implementations or methodologies. For example, models of metabolism or signaling networks can be used in place of the simple state transition rules in the ABM.

To initialize and assess simulations of this hybrid model, we utilize observed data from the WTC-11 hiPSC Single-Cell Image Dataset v1 [37] (Figure 1B). The dataset contains fields of view (FOV) images taken from different locations within hiPS cell colonies. These cell and nuclear images are segmented and used as initial cell and nuclear shapes for simulations, which allows us to start from a realistic initial condition. These simulations are run under a variety of conditions to produce simulated cell and nuclear shapes which are quantitatively compared to the observed cell and nuclear shapes.

### Initial model with volume and adhesion constraints produces a sphere-shaped colony

We start with a model using the basic Hamiltonian consisting of a volume constraint term and an adhesion contact energy term (Figure 2A, Supplemental Table S1A).

**Figure 2.**
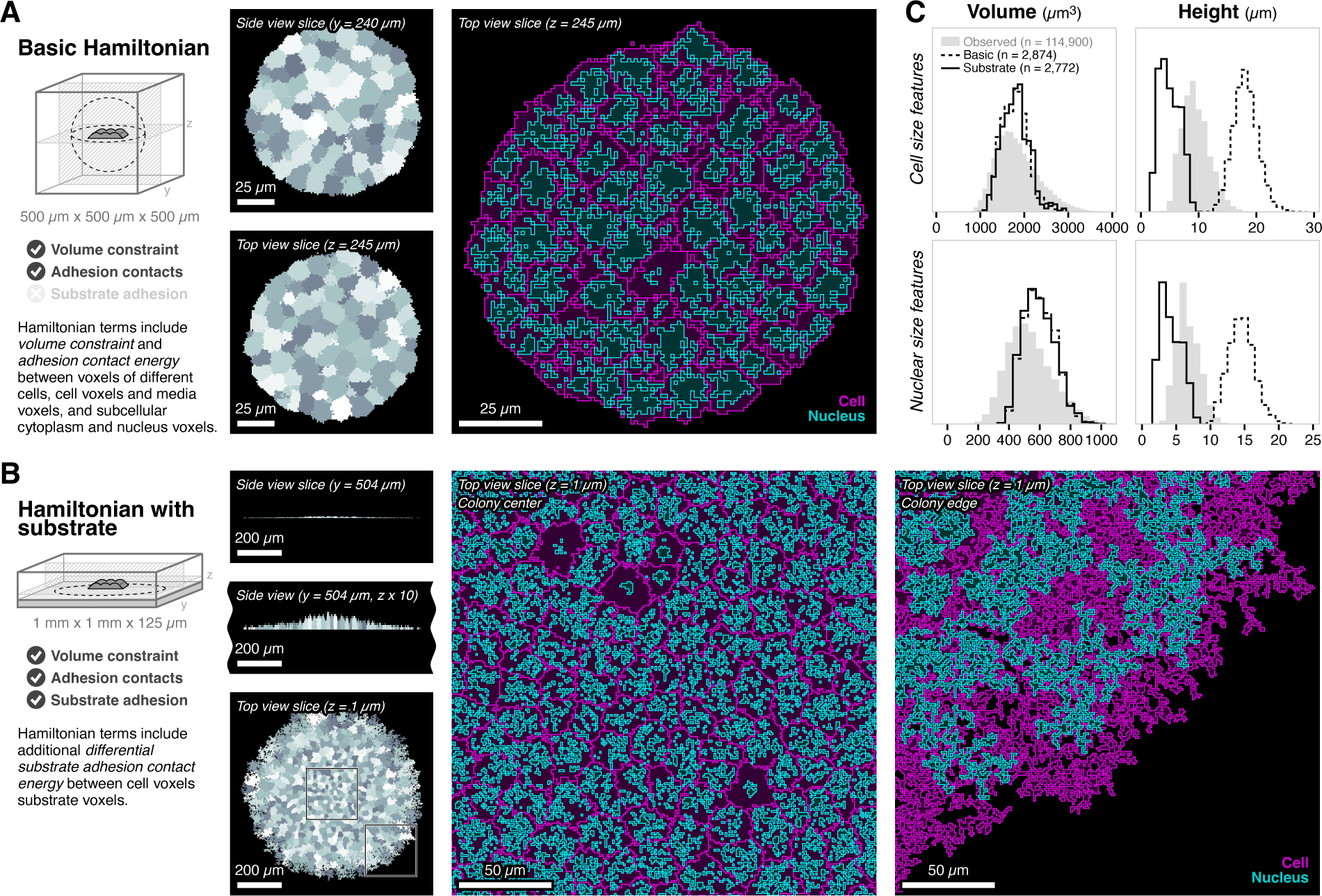
Iterative simulations of hybrid model with different Hamiltonian terms change emergent colony shape from spherical to flat. (**A**) Summary of basic Hamiltonian simulations. Diagram shows gray cells initialized at *t* = 0 days in the center of a 500 *µm* x 500 *µm* x 500 *µm* simulation environment and the dotted black outline shows approximate colony shape at *t* = 4 days. Gray hatched slices indicate the location of the top view (*z* = 245 *µm*) and the side view (*y* = 240 *µm*) shown in the simulation snapshots. Top and side view slices chosen to show the middle of the colony. Gray shading is used to differentiate between cells. Zoomed in simulation snapshot shows the cell and nuclear contours of the top view. (**B**) Summary of Hamiltonian with substrate simulations. Diagram shows gray cells initialized at *t* = 0 days in the bottom of a 1 mm x 1 mm x 125 *µm* simulation environment and the dotted black outline shows approximate colony shape at *t* = 4 days. Gray hatched slices indicate the location of the top view (*z* = 1 *µm*) and the side view (*y* = 504 *µm*) shown in the simulation snapshots. Top view slice chosen to show voxels immediately above the substrate at *z* = 0; side view slice chosen to show the middle of the colony. Gray shading is used to differentiate between cells. Zoomed in simulation snapshots show the cell and nuclear contours of the top view at the center and edge of the colony. (**C**) Histograms of cell and nuclear height and volume for basic Hamiltonian (dotted line) and Hamiltonian with substrate (solid line). Only simulated cells in S phase at *t* = 4 days included. The gray shaded histogram shows observed values of volumes and heights from observed data. See also Figure S1.

Simulations are initialized from select FOVs by placing the filtered and sampled cell and nuclear shapes in the center of a 500 *µm* x 500 *µm* x 500 *µm* environment using a spatial resolution of 1 *µm*/voxel. Simulations are run for 5,760 ticks at a temporal resolution of 1 minute/tick representing 4 days of growth. We run 10 replicates with a simulation wall clock time of 214.2 *±* 10.6 hours (Supplemental Table S2A). All adhesion parameters (cell-cell, cell-media, and subcellular) are set to baseline value of 50.

Simulated cell cycle phase durations are consistent with literature values (Supplemental Figure S1A). At the end of *t* = 4 days, cells form a sphere-shaped colony for all replicates (Figure 2A, Supplemental Figure S1B), consistent with previous 3D CPM models of tumor growth [38, 39]. Simulated cell and nuclear volume distributions are consistent with observed cell and nuclear volume distributions (Figure 2C). However, simulated cell and nuclear height distributions are much taller, averaging 18.1 *±* 2.2 *µm* and 14.5 *±* 1.9 *µm*, compared to observed average heights of 9.5 *±* 2.4 *µm* and 6.7 *±* 1.7 *µm*, respectively. The simulated height distributions are consistent with approximately spherical cells; spheres with volumes 1000 - 3000 *µm*^3^ have diameters of 12 - 18 *µm*.

Because the basic Hamiltonian terms do not have any anisotropy, all dimensions are treated equally to produce sphere-shaped colonies. We note that in 2D, this basic Hamiltonian produces a monolayer by definition. In 3D, we can explore how these basic terms can be adjusted to enable the formation of a simulated hiPS cell monolayer consistent with the observed growth of real cultured hiPS cells.

### Adding environmental constraints and substrate adhesion produces a flattened dome-shaped colony

We hypothesize that adding in a substrate will enable simulated hiPS cell colonies to form and maintain a monolayer. We introduce a layer of substrate voxels at *z* = 0 in the lattice environment and add a term to the Hamiltonian that captures adhesion between cell and substrate voxels (Figure 2B, Supplemental Table S1A).

Simulations are initialized from select FOVs by placing the filtered and sampled cell and nuclear shapes at the bottom of a 1 mm x 1 mm x 125 *µm* environment immediately above the substrate using a spatial resolution of 1 *µm*/voxel. Simulations are run for 5,760 ticks at a temporal resolution of 1 minute/tick representing 4 days of growth. We run 10 replicates with a simulation wall clock time of 214.5 *±* 14.6 hours (Supplemental Table S2A). All adhesion parameters (cell-cell, cell-media, subcellular, and cell-substrate) are set to baseline value of 50.

Simulated cell cycle phase durations remain consistent with literature values (Supplemental Figure S1A). At the end of *t* = 4 days, cells form extremely flat colonies, with cells at the edge of the colony adopting a fingering morphology (Figure 2B, Supplemental Figure S1C). Nuclei at the center of the colony exhibit fingering morphologies within more compact cell shapes while nuclei at the edge of the colony conform to the fingering morphology of the cell. Simulated cell and nuclear volume distributions remain consistent with both the observed volume distributions and the volume distributions from simulations using the basic Hamiltonian (Figure 2C). In contrast, the simulated cell and nuclear height distributions are now much lower than both observed and basic Hamiltonian height distributions, averaging 5.0 *±* 1.8 *µm* and 4.2 *±* 1.6 *µm*, respectively.

We note that this model does not include an explicit height constraint term analogous to the volume constraint term. Instead, the decrease in cell and nuclear heights emerges due to increased cell area as the cells adhere to the substrate. By including a substrate term in the model, we shift simulated hiPS cell colony behavior from sphere-like to monolayer-like. However, simulated cell and nuclear shapes remain inconsistent with observed cell and nuclear shapes and suggest that additional parameter tuning is necessary.

### High-throughput parameter sweep identifies bounds for relative adhesion values

We hypothesize that relative balance between cell-cell adhesion, cell-media adhesion, subcellular adhesion, and cell-substrate adhesion is critical for producing more realistic cell and nuclear shapes. However, full simulations are computationally expensive, averaging 8.9 *±* 0.5 days (Supplemental Table S2A), which make it infeasible to run high-throughput parameter sweeps across these adhesion parameters. Instead, we note that the inconsistent cell shapes appear very early in the simulation and use a data-driven approach to narrow down the parameter space by comparing shape features from short-duration simulations to observed shape features.

We run simulations for all combinations of five different levels of adhesion parameters for a total of 625 conditions (Figure 3A, Supplemental Table S1B). Simulations are run with 10 replicates at a spatial resolution of 1 *µm*/voxel in a 200 *µm* x 200 *µm* x 20 *µm* environment and a temporal resolution of 1 minute/tick for 60 ticks (representing one hour of time). Framework parameters are set to baseline values of 1 Monte Carlo step and a Boltzmann temperature of 10. Simulations have a wall clock time of 2.2 *±* 1.0 minutes, with no noticeable differences between conditions (Supplemental Table S2B).

**Figure 3.**
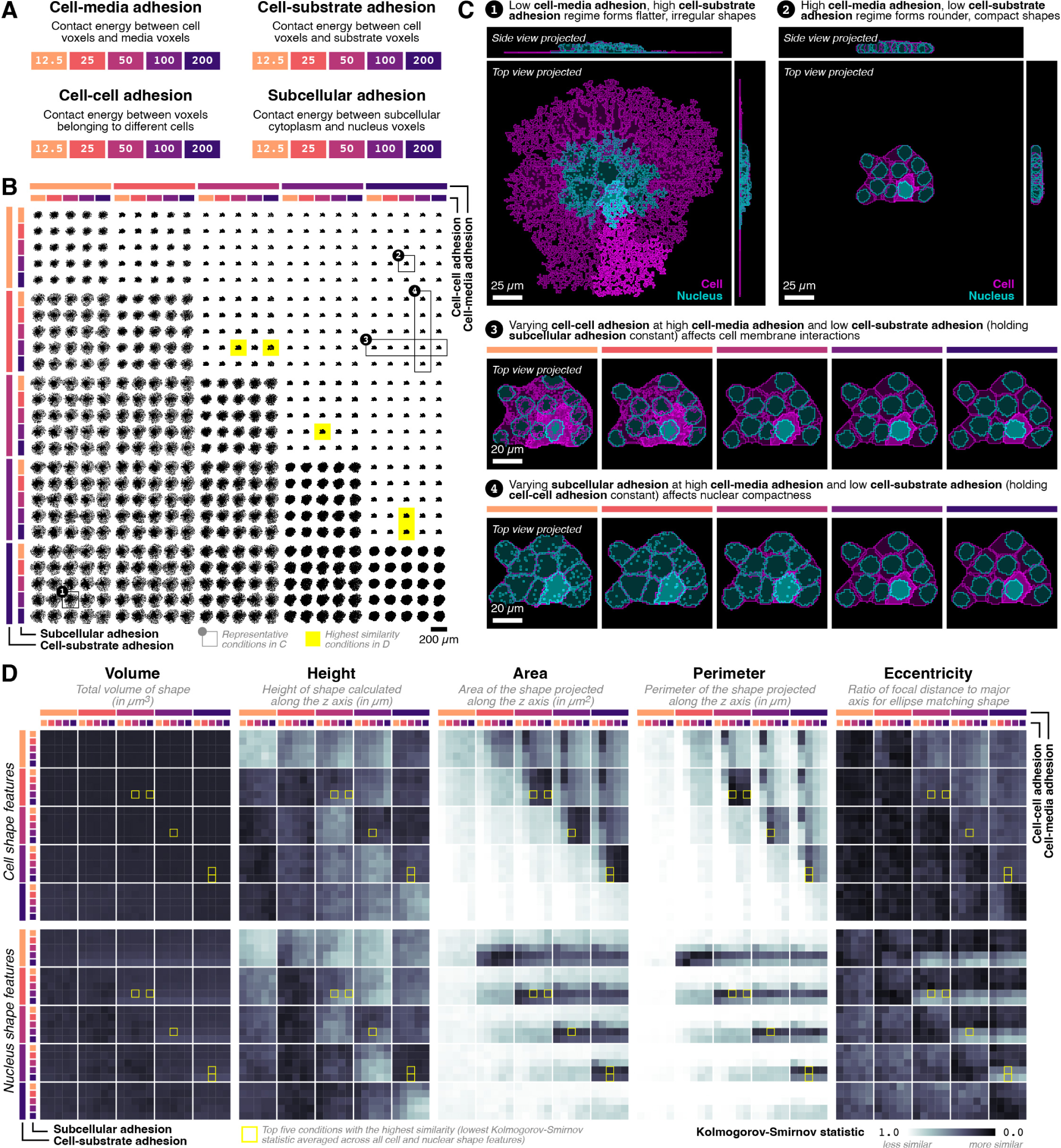
Relative adhesion parameter simulations. (**A**) Summary of the adhesion parameters and levels. (**B**) Contact sheet showing the colony contour projected along the z axis for all conditions for a single replicate at *t* = 1 hour. Conditions annotated with numbered circles and solid outlines indicate key observations corresponding to cell and nuclear contours shown in Panel C. Conditions highlighted in yellow indicate conditions resulting in simulated cell and nuclear shape features with the highest average similarity to observed cell and nuclear shape features, based on the Kolmogorov-Smirnov statistic shown in Panel D. (**C**) Cell and nuclear shape contours for select conditions. Contours are calculated on projected shapes along the z axis for top view or y axis for side view. Representative cell and nuclear shape are highlighted with brighter color. (**D**) Heatmaps showing Kolmogorov-Smirnov statistic calculated between the simulated and observed distributions of different features under different simulation conditions. Darker colors indicate lower Kolmogorov-Smirnov statistic values, which indicates the simulated distribution is more similar to the observed distribution. Conditions producing the highest average similarity are marked by yellow borders (corresponding to the yellow highlights in Panel B). See also Figure S2.

Overall, the relative balance between the different adhesion terms produces diverse cell and colony shapes (Figure 3B, Figure 3C). The balance between cell-media adhesion and cell-substrate adhesion seem to be the strongest drivers of emergent shapes, while cell-cell adhesion and subcellular adhesion produce more subtle effects. In the low cell-media, high cell-substrate adhesion regime, cells and nuclei exhibit very flat, irregular shapes (Figure 3C, 1). In this regime, cell-cell and subcellular adhesion do not have a strong influence. In contrast, in the high cell-media, low cell-substrate adhesion region, cells and nuclei are much rounder and more compact (Figure 3C, 2). In this regime, cell-cell and subcellular adhesion have a stronger influence on emergent cell shapes. Increasing cell-cell adhesion affects how much the cell membrane “mixes” with neighboring cell membranes (Figure 3C, 3). Increasing subcellular adhesion affects the compactness of the nuclei (Figure 3C, 4).

To compare shapes between conditions, we calculate various shape features on the 3D shape (volume and height) and the 2D projection along the z axis (area, perimeter, eccentricity, major axis length, minor axis length, orientation, extent, and solidity) of both simulated and observed cells and nuclei. Quantifying similarity using the Kolmogorov-Smirnov statistic, we see that different features show different patterns of similarity as a function of adhesion conditions (Figure 3D, Supplemental Figure S2K). Cell and nuclear volume and orientation are similar across all conditions as expected; volume is not impacted by the adhesion terms and we do not expect adhesion to favor any polarity in shapes. Height shows subtle differences as a function of cell-media adhesion and cell-substrate adhesion, but generally have high similarity between simulated and observed data with no clear patterns. Area, perimeter, axis lengths, extent, and solidity show a similar pattern of highest similarity in the high cell-media, low cell-substrate adhesion regime and lowest similarity in the low cell-media, high cell-substrate adhesion regime, consistent with our previous qualitative analysis. Within the high cell-media, low cell-substrate regime, there is higher similarity in cell features for higher cell-cell adhesion and higher similarity for nuclear features for higher subcellular adhesion. Eccentricity shows the opposite trend with higher similarity in the low cell-media, high cell-substrate regime and lower similarity in the high cell-media, low cell-substrate regime, although absolute similarity remains high across all conditions. Averaging similarity across metrics, we select the top five relative adhesion conditions that produce cell and nuclear shapes most similar to observed data for further study.

We also evaluate the impact of adhesion parameters on shape features using ANOVA (Supplemental Figure S2A-J). We perform two-way ANOVA for each combination of cell-media and cell-substrate adhesion, using cell-cell and subcellular adhesion as the factors, for each feature. Overall, the low cell-media, high cell-substrate adhesion regime produces cells with higher area, higher perimeter, higher axis lengths, lower extent, and lower solidity compared to cells in the high cell-media, low cell-substrate adhesion regime. Again, volume and orientation do not show significant differences between the different conditions. Generally, there are significant interaction effects between cell-cell adhesion and subcellular adhesion in the high cell-media, low cell-substrate adhesion regime, but no interaction between cell-cell adhesion and subcellular adhesion in the low cell-media, high cell-substrate adhesion regime. This contrast indicates that in the low cell-media, high cell-substrate adhesion regime, the impact of cell-cell adhesion and subcellular adhesion on shapes are independent (and, for certain features, negligible). In the high cell-media, low cell-substrate adhesion regime, where there exist significant interaction effects, cell-cell adhesion shows significant simple main effects on cell features while subcellular adhesion shows significant simple main effects on nuclear features.

Relative cell-cell, cell-media, cell-substrate, and subcellular adhesion produces diverse cell and nuclear shapes. Simulating many different relative adhesion conditions for a short duration and quantitatively comparing simulated shape features to observed data is a feasible approach that enables us to identify relative adhesion conditions that produce simulated shapes most consistent with observed shapes and understand how interactions between adhesion terms impact emergent shapes.

### Parameter sweep on technical terms emphasizes importance of framework specifications

It is critical to understand how the choices of modeling framework parameters impact results, to ensure that insights gained from the model are not artifacts of the framework [14]. The CPM framework utilizes two technical parameters: the number of Monte Carlo steps and the Boltzmann temperature. We aim to understand how these parameters impact results, so we can make informed choices on which parameters to use for simulations that balance biological insight and computational expense.

We run simulations for all combinations of cell-media adhesion and cell-substrate adhesion along with Monte Carlo steps and the Boltzmann temperature for a total of 400 conditions (Figure 4A, Supplemental Table S1C). Simulations are run with 10 replicates at a spatial resolution of 1 *µm*/voxel in a 200 *µm* x 200 *µm* x 20 *µm* environment and a temporal resolution of 1 minute/tick for 60 ticks (representing one hour of time). Cell-cell adhesion and subcellular adhesion are set to baseline values of 50. Simulation wall clock times vary as a function of technical parameters, ranging from 1.9 *±* 1.3 minutes for Monte Carlo steps = 1 to 7.4 *±* 4.8 for Monte Carlo steps = 4 and 1.3 *±* 0.9 minutes for Boltzmann temperature = 1 to 7.6 *±* 4.3 minutes for Boltzmann temperature = 4 (Supplemental Table S2C).

**Figure 4.**
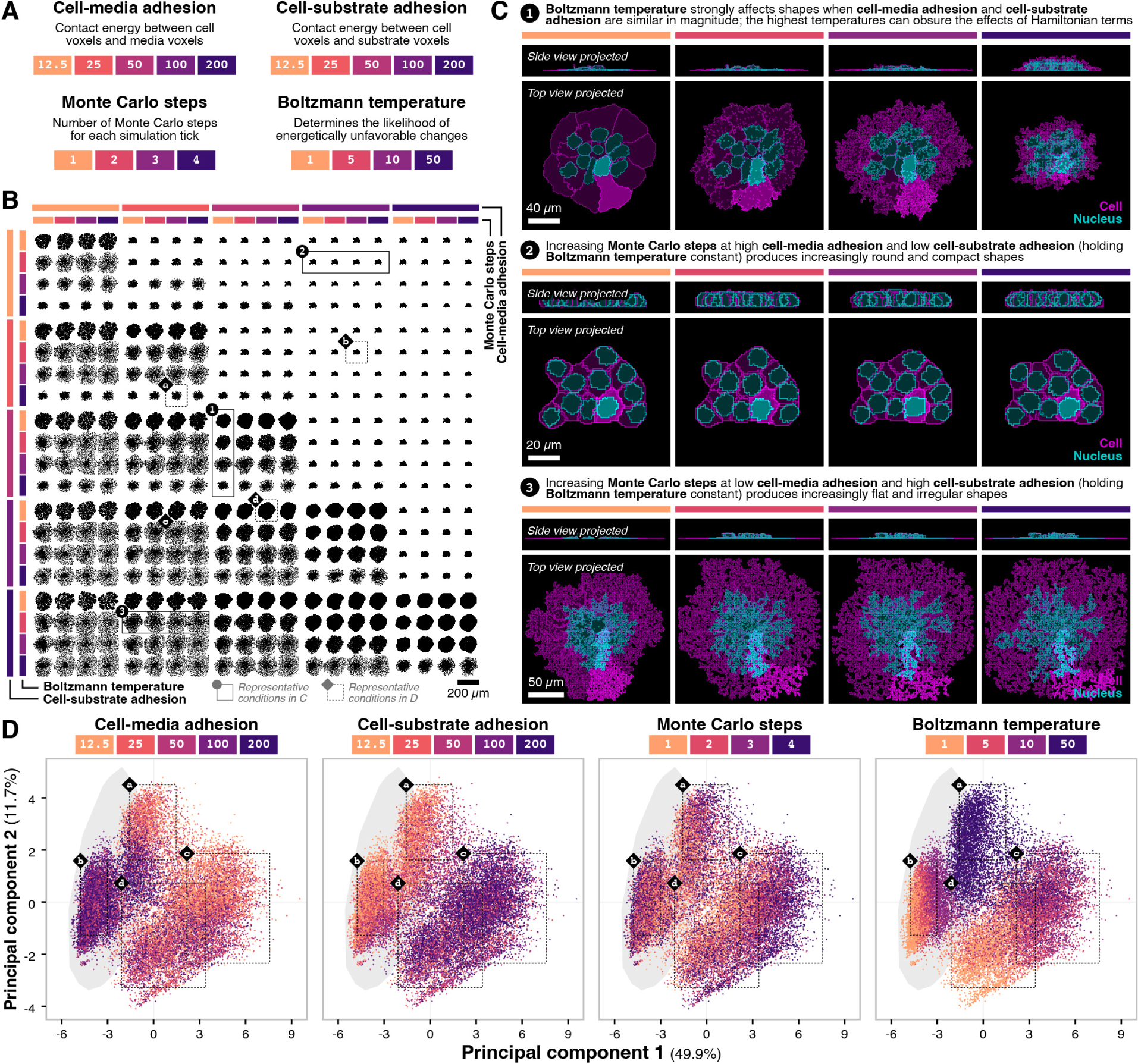
Framework-specific parameter simulations. (**A**) Summary of the adhesion and framework parameters and levels. (**B**) Contact sheet showing the colony contour projected along the z axis for all conditions for a single replicate at *t* = 1 hour. Conditions annotated with numbered circles and solid outlines indicate key observations corresponding to cell and nuclear contours shown in Panel C. Conditions annotated with lettered diamonds and dotted outlines mark select conditions falling within the clusters observed in Panel D. (**C**) Cell and nuclear shape contours for select conditions. Contours are calculated on projected shapes along the z axis for top view or y axis for side view. Representative cell and nuclear shape are highlighted with brighter color. (**D**) Scatter plots of cell and nuclear shape features for different parameter conditions projected into PC space, colored by the parameter level. Each point represents the shape features of a single cell from a simulation with the indicated parameter conditions. Dotted outlines mark the bounding box of shape features from all cells for select conditions (corresponding to the lettered diamond annotations in Panel B). Gray shaded contour area indicates observed data projected into the same PC space. See also Figure S3.

Overall, we see a clear impact of Boltzmann temperature on simulations in the low cell-media, high cell-substrate regime while the impact of the number of Monte Carlo steps is more subtle (Figure 4B, Figure 4C). In the low cell-media, high cell-substrate regime, the lowest Boltzmann temperature produces noticeably less fingering behavior compared to other temperatures (Figure 4C, 1). At the highest Boltzmann temperature, fluctuations due to energetically unfavorable changes produce taller colonies, obscuring the effects of cell-media and cell-substrate adhesion (Figure 4C, 1). The number of Monte Carlo steps seems to push the simulated colony toward more “extreme” shapes: increasing Monte Carlo steps produces more round and compact shapes in the high cell-media, low cell-substrate adhesion regime (Figure 4C, 2) while increasing Monte Carlo steps produces more flat and irregular shapes in the low cell-media, high cell-substrate adhesion regime (Figure 4C, 3).

We apply Principal Component Analysis (PCA) to all the simulated cell and nuclear shape features where two components capture 61.6% of the variance (Figure 4D). PC1 captures variance between cells with flat, irregular morphologies (high area, high perimeter, high axis lengths, low extent, low solidity) and cells with compact, rounded morphologies (low area, low perimeter, low axis lengths, high extent, high solidity). PC2 captures variance in cell and nuclear heights. We see four main regions consistent with our previous qualitative analysis. Region a consists of simulations using the highest temperature in which energetically unfavorable changes have a larger impact on emergent shapes. Region b represents the high cell-media, low cell-substrate adhesion regime at medium Boltzmann temperatures with the highest overlap with observed data. In this region, the number of Monte Carlo steps has a smaller effect as cells are already in an energetically favorable shape. Region c represents the low cell-media, high cell-substrate adhesion regime at medium Boltzmann temperatures where changes in the number of Monte Carlo steps have a larger effect as the Hamiltonian balances between the compact and rounded morphology favored by the cell-media adhesion term and the flat, irregular morphology favored by the cell-substrate adhesion term. Region d represents cells with higher cell-substrate adhesion at the lowest Boltzmann temperature, which form compact but flat shapes.

We evaluate the impact of framework parameters on shape features using ANOVA (Supplemental Figure S3A-J). As previously, volume and orientation are not impacted by the different conditions. Perimeter, axis lengths, extent and solidity show significant interactions between the number of Monte Carlo steps and the Boltzmann temperature across all cell-media and cell-substrate adhesion conditions, indicating that the technical parameters do not have independent effects. Height and area generally show significant interactions in the high cell-media, low cell-substrate adhesion regime and significant main effects in the low cell-media, high cell-substrate adhesion regime.

Framework-specific parameter conditions can dramatically affect simulation outcomes, and impacts may be non-linear with respect to other simulation conditions. Understanding how these framework-specific parameters impact results is important for selecting appropriate conditions and avoiding framework-dependent artifacts.

### Spatial and temporal resolution terms have variable effect based on emergent dynamics

Simulations can be run at higher or lower spatial and temporal resolution, with higher resolutions providing increased accuracy at higher computational costs. Depending on the purpose, a lower spatial or temporal resolution may be sufficient. We aim to assess how higher or lower resolution can influence model outcomes.

We run simulations for all combinations of cell-media adhesion and cell-substrate adhesion along with spatial and temporal resolution for a total of 200 conditions with 10 replicates each (Figure 5A, Supplemental Table S1D). Low spatial resolution simulations use a spatial resolution of 2 *µm*/voxel in a 100 *µm* x 100 *µm* x 10 *µm* environment while high spatial resolution simulations use a spatial resolution of 1 *µm*/voxel in a 200 *µm* x 200 *µm* x 20 *µm* environment. Low temporal resolution simulations use temporal resolution of 5 minutes/tick for 12 ticks while high temporal resolution simulations use temporal resolution of 1 minute/tick for 60 ticks (all representing one hour of time). Cell-cell adhesion and subcellular adhesion are set to baseline values of 50; framework parameters are set to baseline values of 1 Monte Carlo step and a Boltzmann temperature of 10. Simulation wall clock time varies as a function of resolution: low spatial resolution uses 2.7 *±* 1.8 sec while high spatial resolution uses 79.9 *±* 68.7 sec, and low temporal resolution uses 14.2 *±* 15.8 sec while high temporal resolution uses 68.3 *±* 77.4 sec (Supplemental Table S2D).

**Figure 5.**
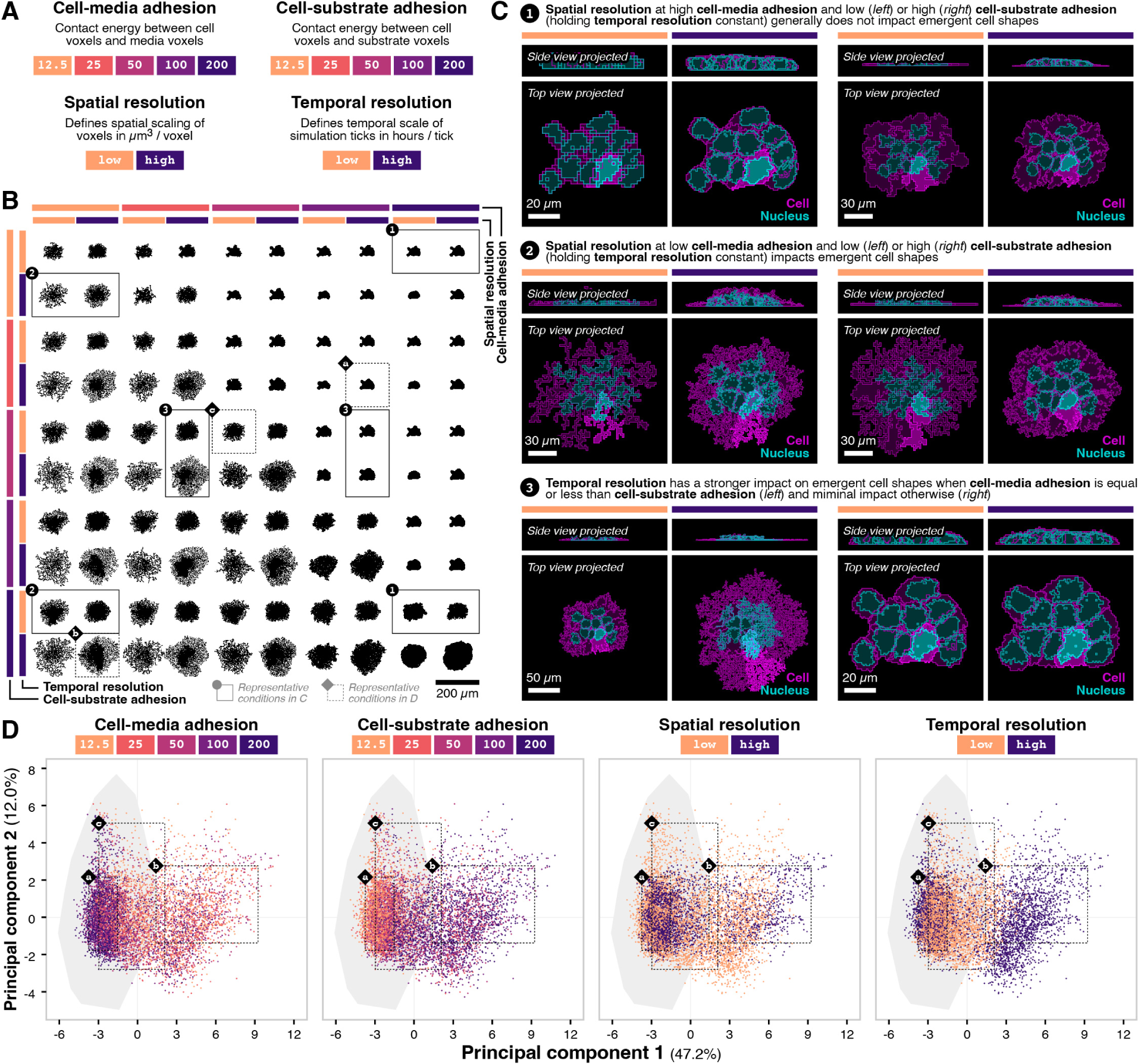
Spatial and temporal resolution parameter simulations. (**A**) Summary of the adhesion and resolution parameters and levels. (**B**) Contact sheet showing the colony contour projected along the z axis for all conditions for a single replicate at *t* = 1 hour. Conditions annotated with numbered circles and solid outlines indicate key observations corresponding to cell and nuclear contours shown in Panel C. Conditions annotated with lettered diamonds and dotted outlines mark select conditions falling within the clusters observed in Panel D. (**C**) Cell and nuclear shape contours for select conditions. Contours are calculated on projected shapes along the z axis for top view or y axis for side view. Representative cell and nuclear shape are highlighted with brighter color. (**D**) Scatter plots of cell and nuclear shape features for different parameter conditions projected into PC space, colored by the parameter level. Each point represents the shape features of a single cell from a simulation with the indicated parameter conditions. Dotted outlines mark the bounding box of shape features from all cells for select conditions (corresponding to the lettered diamond annotations in Panel B). Gray shaded contour area indicates observed data projected into the same PC space. See also Figure S4.

Overall, we see a larger impact of resolution in the low cell-media, high cell-substrate adhesion regime compared to the high cell-media, low cell-substrate adhesion regime (Figure 5B, Figure 5C). When cell-media adhesion is high, spatial resolution generally does not have an impact on the shapes (other than the actual spatial resolution) for both low and high values of cell-substrate adhesion (Figure 5C, 1). However, when cell-media adhesion is low, spatial resolution affects emergent cell and nuclear shapes beyond expected differences due to spatial resolution for both low and high values of cell-substrate adhesion (Figure 5C, 2). Temporal resolution plays a strong role in emergent shapes when cell-media adhesion is equal or less than the cell-substrate adhesion, and a negligible effect otherwise (Figure 5C, 3).

We again apply PCA to all the simulated cell and nuclear shape features where two components capture 59.6% of the variance (Figure 5D). PC1 again captures variance between cells with flat, irregular morphologies (high area, high perimeter, high axis lengths, low extent, low solidity) and cells with compact, rounded morphologies (low area, low perimeter, low axis lengths, high extent, high solidity). PC2 now captures variance between tall and large cells (high volume, high height) and short and small cells (low volume, low height). We see three main regions. Region a represents the high cell-media, low cell-substrate adhesion regime where spatial and temporal resolution do not noticeably impact cell and nuclear shapes, and remain consistent with observed shapes. Regions b and c represent the low cell-media, high cell-substrate adhesion regime where region b contains high temporal resolution simulations and region c contains low temporal resolution simulations. The cell-substrate adhesion term produces dramatic changes in cell and nuclear shape and temporal resolution impacts how often those shapes are changed.

Finally, we quantify the impact of resolution on shape features using ANOVA (Supplemental Figure S4A-J). Across features, there are generally not statistically significant interaction effects in the high cell-media, low cell-substrate adhesion regime. When the interaction is significant, the feature values are often not qualitatively different in magnitude. In the low cell-media, high cell-substrate adhesion regime, there are significant interaction effects for area, perimeter, extent, and solidity indicating that the choice of spatial and temporal resolution is not independent. In contrast, height and eccentricity only show significant main effects (no interaction). These computational models can be run at different resolutions that may be appropriate for different questions. One can reduce temporal or spatial resolution in favor of computational efficiency in cases where the additional detail is not necessary.

### Full scale simulations of selected adhesion parameter conditions produce diverse colony and cell shapes

We now run full simulations for the top five conditions producing cell and nuclear shapes most similar to those in our observed data, combined with our updated understanding of how technical parameters can impact emergent outcomes (Supplemental Table S1E). We used Monte Carlo steps of 3 and a Boltzmann temperature of 5, both of which are middle values that we do not expect to produce any framework-specific artifacts. The selected conditions fall closer to the high cell-media, low cell-substrate adhesion regime where low spatial and temporal resolution are likely sufficient. However, we are interested in cell shapes and colony dynamics and choose to use high spatial and temporal resolution.

Simulations use a 1 mm x 1 mm x 125 *µm* environment at a spatial resolution of 1 *µm*/voxel and run for 5,760 ticks at a temporal resolution of 1 minute/tick representing 4 days of growth. We run a single replicate for each condition with a simulation wall clock time of 405.6 *±* 7.7 hours (Supplemental Table S2E).

Conditions A, B, and C represent combinations of high, medium, and low relative cell-media and cell-substrate adhesion, respectively, while holding cell-cell adhesion constant at 50 and subcellular adhesion constant at 100. Condition D is equivalent to condition A, with increased subcellular adhesion of 200. Condition E is equivalent to condition C, with increased cell-cell adhesion of 200.

Overall, each condition produces distinct colony shapes after 4 days of growth, despite similar behavior in the shorter, parameter sweep simulations with only 1 hour of growth. Condition A forms a square-shaped colony due to high cell-media adhesion, which makes it energetically unfavorable for cell voxels to be adjacent to media voxels (Figure 6A). Condition B forms a square-shaped colony with rounded edges compared to condition B due to the lower cell-media adhesion (Figure 6B). Condition C forms a more circular colony as cell-media adhesion is further decreased (Figure 6C). As cell-media adhesion decreases, cell-substrate adhesion must also decrease, which results in taller cells and colonies. Keeping cell-substrate adhesion high while decreasing cell-media adhesion would bring us into the low cell-media adhesion, high cell-substrate adhesion regime characterized by flat, irregular shapes. For condition D, with increased subcellular adhesion relative to condition A, the overall colony is slightly taller due to more compact nuclei. For condition E, with increased cell-cell adhesion relative to condition A, we observe an extreme behavior where the increased cell-cell adhesion contact energy causes cells to avoid touching other cells.

**Figure 6.**
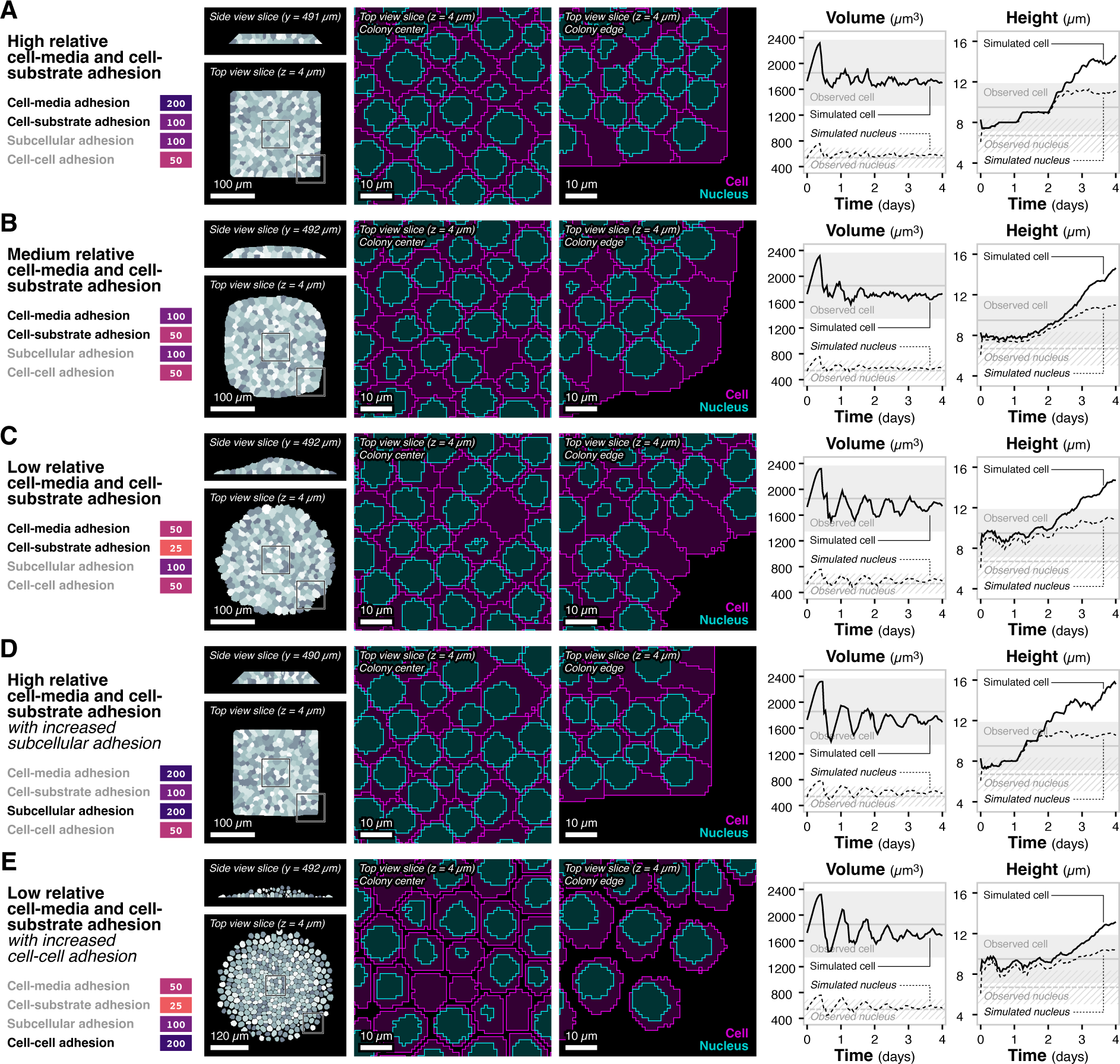
Full simulations for selected adhesion parameter conditions. (**A**) Simulation snapshots and metric time courses for the high relative cell-media and cell-substrate adhesion condition. Colony views show top and side view of cell shapes. Gray shading is used to differentiate between cells. Zoomed in views show top view of cell and nuclear contours at the center and edge of the colony. Time courses show average cell (solid) and nuclear (dotted) volume and height. Solid gray horizontal line and shaded area show mean and standard deviation, respectively, of cell volumes and heights in the observed data. Dotted gray horizontal line and hatched area show mean and standard deviation, respectively, of nuclear volumes and heights in the observed data. (**B**) Simulation snapshots and metric time courses for the medium relative cell-media and cell-substrate adhesion condition. *Analogous to Panel A*. (**C**) Simulation snapshots and metric time courses for the low relative cell-media and cell-substrate adhesion condition. *Analogous to Panel A*. (**D**) Simulation snapshots and metric time courses for the high relative cell-media and cell-substrate adhesion with increased subcellular adhesion condition. *Analogous to Panel A*. (**E**) Simulation snapshots and metric time courses for the low relative cell-media and cell-substrate adhesion with increased cell-cell adhesion condition. *Analogous to Panel A*. See also Figure S5.

All five conditions are able to form a simulated hiPS cell monolayer until day 2, at which point they are no longer able to maintain a monolayer and instead begin to grow on top of each other (Supplemental Figure S5). While average simulated cell and nuclear volume trajectories over time are consistent over time and with observed data, average cell and nuclear height trajectories reflect this transition from a monolayer to a multilayer colony (Figure 6). As relative cell-media and cell-substrate adhesion decreases from condition A to B to C, changes in cell and nuclear height trajectories are non-monotonic. For condition A, we observe an almost stepwise increase in height over time, passing the observed heights around day 2.5. For condition B, increase in height is very smooth, passing the observed heights around day 3. For condition C, height shows a periodic behavior for the first 2 days before passing the observed heights around day 3. Condition D matches condition A until day 1, after which cell and nuclear heights increase much faster and pass observed heights at day 2. With the higher subcellular adhesion in condition D, nuclei are less compressible and cause cell heights to match nuclear heights. Condition E matches condition C until day 1 and then maintains the periodic behavior until day 2.5 and low heights until day 4. The higher cell-cell adhesion in condition E provides a driving force that increases movement of cells at the edge of the colony, creating additional space for growth of cells at the center of the colony. While this condition maintains a monolayer for the longest of the five conditions, the individual simulated cell behavior is unrealistic.

Relative adhesion parameter, framework parameter, and resolution parameter sweeps at short durations identified simulation conditions that produce colonies that are much more consistent with observed colonies in longer duration simulation. These simulated cells grow as a monolayer colonies for the first 2-3 days, but subsequently lose their monolayer behavior and begin to grow on top of each other, which indicates that these simple mechanisms (volume constraint and adhesion contact energies) are not sufficient to explain how real hiPS cell monolayers are formed and maintained in cultured biological systems.

## Discussion

Initially, we aimed to uncover rules that lead to the monolayer formation and maintenance observed in our cultured hiPS cell colonies. To achieve this, we integrated an ABM with a CPM to build a hybrid model that characterizes and interrogates the agent rules driving 3D cell shape and colony dynamics in simulated hiPS cells. Our goal was to find and test rules that could mimic, and then more broadly explore, the dynamics observed in real-world hiPS cell cultures. The initial outputs showed great potential, appearing to mimic small colony behaviors at first glance. However, after conducting longer runs and performing deep quantitative comparisons to the observed data, the results came up short. This model development approach provides interesting biological and technical insights, and the flexible 3D hybrid model could be further developed for diverse cell biology applications. Consequently, we focused on deeply examining the potential and limitations of our hybrid model to inform the community and recommend further development ideas for systems where critical cell-wide mechanics play significant roles, as observed in our hiPS cell research.

The CPM was implemented to support 2D and 3D cell shapes using a Monte Carlo algorithm that prevents cell fragmentation. Through iterative model development, we demonstrated that the basic CPM Hamiltonian terms of volume and adhesion are not sufficient to produce cell shapes and colony dynamics consistent with live cell imaging of hiPS cell cultures. Adding substrate adhesion produces flatter colonies, highlighting the importance of accounting for environmental context when characterizing and iteratively modeling cell dynamics. Through data-driven model development, we conducted high-throughput parameter sweeps of biologically-relevant adhesion parameters and technically-relevant model framework and simulation resolution parameters. This analysis identified regions of the parameter space that produce cell shapes consistent with those observed in cultured hiPS cells. It also highlighted how specific hybrid model decisions can impact emergent simulated shape dynamics. Finally, we ran full-scale simulations of our model and demonstrated that certain combinations of agent rules are sufficient to form, but not maintain, a simulated hiPS cell monolayer in 3D.

Iterative and data-driven model development enables the systematic evaluation of how model rules and parameters drive emergent 3D cell shape and colony dynamics. As with all methodologies, it is important to note the limitations. First, the full-scale simulations are computationally expensive, requiring wall clock times over four times the simulated growth time. Second, the observed data used to train and benchmark simulations, from the WTC-11 hiPSC Single-Cell Image Dataset v1 [37], is not temporal. We therefore avoid making temporal comparisons and emphasize the challenges of comparing dynamic simulations to static observations. High resolution, single-cell temporal imaging data of large colony regions will be critical for further model development. Finally, our simulations do not fully emulate the experimental conditions under which the observed data was collected. While we match the total growth duration of the experiments, our simulations start with fewer cells and the simulated colony sizes are smaller than those in the observed data. In this study, we focus on exploring how different model rules and parameters impact simulated cell and colony shapes, rather than exactly reproducing cell and colony features. In order to achieve the latter, these limitations would need to be addressed.

A major challenge in cell biology is that simulating certain cell and tissue behaviors in 3D (such as the emergence of monolayers) is not always as simple as adding a third dimension to an existing 2D model. While our goal was to build the simplest model that could reproduce observed behaviors, our use of 3D CPMs to simulate monolayers proved challenging with the basic “out of the box” CPM rules. This study highlights how additional, alternative, or more complex agent state and shape rules may be necessary to more accurately characterize cell and colony simulations. Notably, standard 3D CPMs do not natively incorporate long-range, cell-scale mechanics. Emergent dynamics arise from integrating global measurements, such as volume, with local, voxel-scale interactions, which only involve nearest neighbors. As we now observe, these default interactions cannot mimic the organized or directional pushing and pulling forces of cytoskeletal structures, which often need to span across all the voxels that make up the diameter of any given cell. Our results indicate that these mechanics are likely necessary to capture the formation and maintenance of hiPS cell monolayer and this 3D CPM limitation likely extends to contexts in which cell-scale forces, such as elasticity, dominate the overall 3D colony-scale behavior of the system. To address this shortcoming, one could consider systematically adding mechanical constraints to the CPM Hamiltonian at the cell agent level to mimic the effects of forces (increasing model complexity), or integrate local agent rules with global rules derived from separate models operating at a higher spatiotemporal scale (increasing model abstraction). An alternative strategy could be to replace the CPM with a modeling methodology that directly represents cell mechanics [40, 41, 42]. While there may be opportunities to further develop and apply our modeling approach; as with all models and their applications, the tolerance for model complexity, assumptions, and accuracy are subjective and should be deeply considered when selecting a model structure and framework.

Advances in multi-scale models capable of integrating multi-modal subcellular-, cell-, and colony-scale mechanics present an active area of research that shows great promise. We believe that sharing both the model and its development process fosters a culture of open team science and further engages diverse researchers to participate in collaborative systems biology. As we continue our efforts toward understanding hiPS cells, we hope that documenting and disseminating the model development process (in addition to the model) will support the interpretation and application of computational models of complex biological systems.

## Methods

### Model framework

We extend our previously published agent-based modeling framework ARCADE [36, 18, 14] to integrate a Cellular Potts Model (CPM) of cell and subcellular shapes.

#### Packages, interfaces, and classes

We separate the existing `patch` implementation for tumor cell growth and vasculature modeling into a dedicated package and implement the CPM in the `potts` package. Interfaces and abstract classes shared between the two implementations are placed in the `core` package. Within each package are three main packages that manage the overall simulation (`sim`), agents within the simulation (`agent`), and the simulation environment (`env`).

The `sim` package contains classes for running simulations, loading inputs, and saving outputs.

The `agent` package contains interfaces and classes related to model agents:

- `cell` for representing individual cell agents (e.g. tumor cell, healthy tissue cell, stem cell)
- `module` for representing exclusive subcellular behaviors or mechanisms of a `cell` (e.g. cell cycle)
- `process` for representing non-exclusive subcellular behaviors or mechanisms of a `cell` (e.g. metabolism, signaling networks)
- `action` for representing non-cell entities that interact with a `cell` (e.g. external perturbations to simulate a wound or treatment)

The `env` package contains interfaces and classes related to the model environment:

- `location` for representing individual `grid` locations (e.g. hexagonal patch, collection of voxels)
- `grid` for representing the grid on which `cell` agents exist
- `lattice` for representing environment layers (e.g. nutrients, signaling molecules)
- `component` for representing non-agent entities in the environment (e.g. vasculature)
- `operation` for representing environmental behaviors or mechanisms within a `lattice` (e.g. diffusion) Additional packages provide support for visualization (`vis`) and various utility classes (`util`).

Key `potts` implementation interfaces and classes are summarized in Table 1 and covered in detail in the following sections.

**Table 1.** Key potts implementation classes and interfaces.

| Package | Class / Interface | Description |
| --- | --- | --- |
| `potts.sim` | `Potts` | Implementation of CPM algorithm |
| `potts.sim.hamiltonian` | `Hamiltonian` | Interface for Hamiltonian terms |
| `potts.agent.cell` | `PottsCell` | Simple stem cell agent compatible with CPM |
| `potts.agent.module` | `PottsModuleProliferation` | Cell cycle for cells in proliferative state |
| `potts.agent.module` | `PottsModuleApoptosis` | Cell death for cells in apoptotic state |
| `potts.env.location` | `PottsLocation2D` | Collection of voxels for 2D CPM |
| `potts.env.location` | `PottsLocation3D` | Collection of voxels for 3D CPM |

#### Cellular Potts Model

The CPM is a modeling framework in which cell shapes are represented by the collection of voxels on a lattice that all share the same identity [30, 24]. Dynamics within a CPM are driven by the minimization of energy described by a Hamiltonian, evolved through a Monte Carlo algorithm.

##### Hamiltonian

The original formulation of the Hamiltonian is given by:

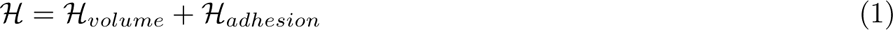

where *ℋ_volume_* represents the volume constraint keeping cell agents near a target volume and *ℋ_adhesion_* represents adhesion contact energies between voxels in the lattice.

The volume constraint is calculated as:

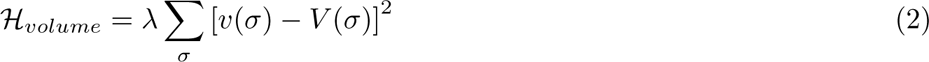

where *λ* represents the strength of the volume constraint, *σ* indexes across cells, *v* is the current volume, and *V* is the target volume. For simulations in 2D, this term uses area in place of volume.

The adhesion term can be separated into the cell-cell and cell-media terms, calculated as:

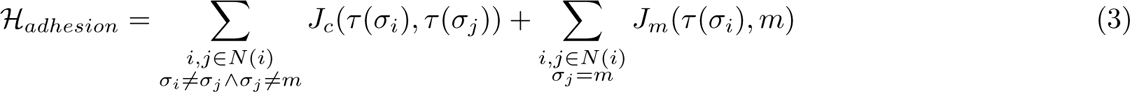

where *i* and *j* index across lattice sites, *N* (*i*) are the lattice sites neighboring site *i*, *σ* is the identity of the cell, *m* represents media, *τ* is the cell type, *J_c_* is the strength of the cell-cell adhesion contact energy, and *J_m_* is the strength of the cell-media adhesion contact energy. We expand the adhesion term to account for subcellular adhesion contact energy between nuclear and cytoplasmic voxels, calculated as:

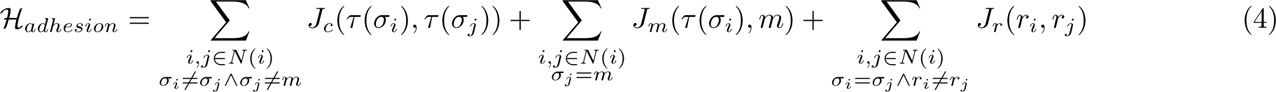

where *r* is the region type and *J_r_* is the strength of the subcellular adhesion contact energy. For substrate adhesion, we add an additional term to the Hamiltonian:

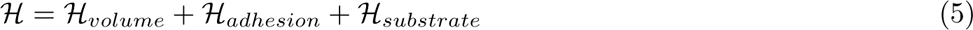

where *ℋ_substrate_*represents the contact energies between cell voxels and a substrate located at *z* = 0.

The substrate term is analogous to the adhesion terms, calculated as:

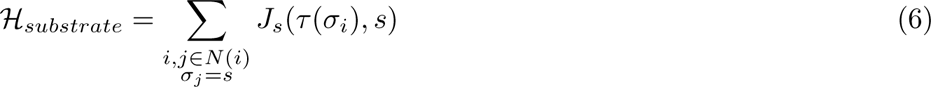

where *s* represents substrate and *J_s_*is the strength of the substrate adhesion contact energy.

##### Monte Carlo algorithm

We use the Monte Carlo algorithm described in [43], which prevents cell fragmentation and improves computational efficiency, and extend it to support 3D simulations and enable subcellular regions. The algorithm for a copy attempt is summarized below. Note that cells with and without subcellular regions can be simulated together; items in *italics* apply only for cells with subcellular regions.

1. Randomly select a candidate lattice site *i* corresponding to identity *σ_source_ and region r_source_*
2. Randomly select a target identity *σ_target_* from identities in the neighborhood of site *i*. *Randomly select a target region r_target_ from identities in the neighborhood of site i. If both a valid target identity and a valid target region exist, randomly select which to update.* If no valid option exists, skip to final step.
3. If updating identity and *σ_source_* is not media, check the local connectivity of cell *σ_source_*. If connected, proceed to next step. Otherwise, proceed to final step.
4. *If r_source_ is not cytoplasm, check the local connectivity of region r_source_. If connected, proceed to next step. Otherwise, proceed to final step*.
5. If updating identity and *σ_target_* is not media, check the local connectivity of cell *σ_target_*. If connected, proceed to next step. Otherwise, proceed to final step.
6. *If r_target_ is not cytoplasm, check the local connectivity of region r_target_. If connected, proceed to next step. Otherwise, proceed to final step*.
7. Calculate change in Hamiltonian energy Δ*H* resulting from changing identity of lattice site *i* from *σ_source_* to *σ_target_*. *If updating region, instead calculate change in Hamiltonian energy* Δ*H resulting from changing region of lattice site i from r_source_ to r_target_*.
8. Accept the change with probability

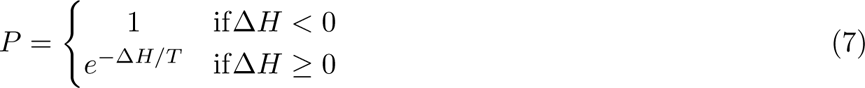

where *T* is the simulation temperature.

9. Increment the number of copy attempts

A single Monte Carlo step consists of *N* = *L × W × H* copy attempts, where *L*, *W*, and *H* are the length, width, and height, respectively, of the lattice. Multiple Monte Carlo steps can be performed during each tick of the simulation.

#### Cell agents

Cell agents represent simple stem cells, with a unique identity corresponding to a collection of voxels on the CPM lattice. Cells exist in one of two states: proliferative or apoptotic. Cell state defines the set of rules that the agent follows at each tick of the simulation.

##### Proliferative cells

Proliferative cells represent actively growing and dividing cells. Proliferative cells progress from G1 to S to G2 to M phase, where the duration of each phase is defined by a Poisson distribution. A cell will transition to the next phase after the appropriate number of steps, drawn from the phase-specific Poisson distribution, have passed. Poisson distributions are parameterized based on the Erlang distributions fit to single-cell measurements of phase duration in H9 cells reported in [44].

In each phase, cells update their target cell volume as a function of cell growth rate and baseline cell volume. For cells with nuclei, cells also update their target nuclear volume as a function of nuclear growth rate and baseline nuclear volume. Baseline volumes are estimated from the WTC-11 hiPSC Single-Cell Image Dataset v1 [37]. Growth rates are set to enable doubling of average cell volume over the course of an average cell cycle. These target volumes are used to define *V* for calculations of the Hamiltonian during the CPM step.

Cells in G1 phase and G2 phase have a low change of switching to an apoptotic state [45]. Cells in G2 phase additionally have a size checkpoint, in which they will not proceed to M phase until they have reached a specified volume threshold to prevent cells from dividing prematurely.

Once cells complete M phase, they will divide into two daughter cells. All the voxels in the CPM that belong to the mother cell are separated along the shortest axis and divided between the two daughter cells, ensuring that they remain simply connected.

##### Apoptotic cells

Apoptotic cell represent cells undergoing programmed cell death. Apoptotic cells progress from early to late apoptosis, analogous to proliferative cells. Duration and growth rate parameters are approximated based on average apoptosis durations [45]. When a cell falls below a threshold volume, the cell agent is removed from the simulation and all corresponding voxels in the CPM are set to media.

#### Model environment

Each cell agent is associated with location object that contains a list of voxels corresponding to sites in the CPM lattice with the same identity. For 2D simulations, each voxel contains *x* and *y* coordinates. For 3D simulations, each voxel contains *x*, *y*, and *z* coordinates. For simulations with subcellular regions, each location object contains additional lists of voxels corresponding to sites in the CPM lattice with the same identity separated by region.

The CPM lattice is kept in sync with the voxel lists. When a candidate lattice site is changed, the corresponding voxel is moved from the source cell to the target cell. Changes in candidate lattice sites from a media voxel to a cell voxel will add voxels to the voxel list. Changes in candidate lattice sites from a cell voxel to a media voxel will remove voxels to the voxel list.

Voxels lists are only updating during cell division and cell death, in which the lists of voxels are split or removed, respectively. The corresponding sites on the CPM lattice are updated based on the resulting lists.

To avoid additional boundary checks, the CPM lattice contains a one voxel boundary in all directions.

### Model simulations

All simulations are initialized using cell and nuclear shapes from the WTC-11 hiPSC Single-Cell Image Dataset v1 [37]. FOVs are selected based on the following criteria:

- FOV does not contain any outliers
- All cells in the FOV have the cell stage "M0" denoting cells in interphase
- FOV contains 11 cells (the average number of cells per FOV in the dataset)

We further filter the FOVs to select 10 FOVs (corresponding to 10 replicates) that provide a volume distribution representative of the overall volume distribution of the dataset. All cells in the dataset are binned by volume and counts in each bin are rescaled for a total of 10 cells. For bins (volume in pixels) of [0, 396916, 793832, 1190748, 1587664, 1984580, 2381496, 2778412, 3175328, 3572245, 3969161], we have rescaled counts of [0, 0, 3, 4, 2, 1, 0, 0, 0, 0]. The following FOV segmentations were selected:

- Replicate 0: `fov_seg_path/000c9e5e08b0616b9bdf8f01c93f9136ec4a22fc7b29c52dc6c4e4fc7e52fa5f_ 3500003034_100X_20190520_2-Scene-017-P17-D07_CellNucSegCombined.ome.tiff`
- Replicate 1: `fov_seg_path/002b582bd22765d1ff386c72edfb8c6f98807cd892f7e2a3898c3f649bff17cc_3500001881_100X_20180330_1e-Scene-08-P22-E05_CellNucSegCombined.ome.tiff`
- Replicate 2: `fov_seg_path/0073601263e599d4e7baa748667238ae9124bd562fca148507d0442cd4f83082_ 3500001890_100X_20180402_4-Scene-19-P57-F06_CellNucSegCombined.ome.tiff`
- Replicate 3: `fov_seg_path/01120fe20a69b4ca884253bbc470a1cbe4c77e94e2c2faa043a392435d273237_3500001541_100X_20171114_2-Scene-15-P34-F08_CellNucSegCombined.ome.tiff`
- Replicate 4: `fov_seg_path/0128eecf759f4c2b6ad5f6886edfd1b0dae58cd2eec4c858c564fffe92a8f771_ 3500002017_100X_20180508_1c-Scene-13-P42-F07_CellNucSegCombined.ome.tiff`
- Replicate 5: `fov_seg_path/02237b60a56fc6fec90f4684e71dbd3fa01978a56e5fbe10b76e2c5d43092625_3500002059_100X_20180522_1r-Scene-11-P35-E07_CellNucSegCombined.ome.tiff`
- Replicate 6: `fov_seg_path/03c56fd547a7fdf0b7016d6c7e340d05e2fb56ce709cf6980860d2edfe9b3d28_ 3500003044_100X_20190531-Scene-072-P72-F09_CellNucSegCombined.ome.tiff`
- Replicate 7: `fov_seg_path/060082578e5a0d1ddd976597440f19ce2ceb02ba537c095c6308d2a8e2e22cf6_3500002810_100X_20190318-Scene-057-P57-F10_CellNucSegCombined.ome.tiff`
- Replicate 8: `fov_seg_path/078d2fad1399b237517196282bb0a2e3b782614d92a418643d328e69e17ff900_ 3500001123_100X_20170725_1-Scene-2-P2-E05_CellNucSegCombined.ome.tiff`
- Replicate 9: `fov_seg_path/39fdd9fa3fb2967a3033291564004069b1d76a157422a5fe3534ed335fa5b780_3500000847_100X_20170419_5-Scene-03-P33-F05_CellNucSegCombined.ome.tiff`

Each FOV was then sampled to a resolution of 1 *µ*m / voxel or 2 *µ*m / voxel and converted to the ARCADE `LOCATIONS` file format. Individual cell data, including cell and nuclear volumes and heights, are converted to the ARCADE `CELLS` file format. The `CELLS` file also includes initial cell state and phase estimated based on cell volume assuming a linear increase in volume over time.

Simulation input files are provided in Table SS1. Simulation wall clock times are provided in Table SS2.

### Simulation analysis

#### Cell and nuclear shape features

Simulated and observed cell and nuclear shapes are quantified using the shape features summarized in Table 2. Simulated features are calculated by converting model outputs to image arrays. Observed features are calculated from the segmentations of cells in all selected FOVs from the WTC-11 hiPSC Single-Cell Image Dataset v1 (*n* = 114, 900) [37].

**Table 2.** Cell and nuclear shape features. Features marked with a * are calculated on shapes projected along the z axis using the Python package *scikit-image* module `measure` method `regionprops`. Voxels refer to the 3D shape; pixels refer to the projected 2D shape. Features with units are scaled using the appropriate voxel size of the observed or simulated data; unitless features are not adjusted.

| Feature | Description | Units |
| --- | --- | --- |
| Volume | Total number of voxels | $\mu\text{m}^3$ |
| Height | Range of voxels in the z axis | $\mu\text{m}$ |
| Area | Number of pixels in the region | $\mu\text{m}^2$ |
| Perimeter | Length of the line through the center of border pixels | $\mu\text{m}$ |
| Eccentricity | Ratio of focal distance over major axis length of ellipse with same second moments | - |
| Major axis length | Major axis length of the ellipse with same normalized second moment | $\mu\text{m}$ |
| Minor axis length | Minor axis length of the ellipse with same normalized second moment | $\mu\text{m}$ |
| Orientation | Angle between the 0th axis and major axis of ellipse with same second moments | - |
| Extent | Ratio of pixels in the region to pixels in the total bounding box | - |
| Solidity | Ratio of pixels in the region to pixels of the convex hull image | - |

#### Kolmogorov-Smirnov similarity testing

In Figure 3D, similarities between observed and simulated cell and nuclear shape features are quantified using the Kolmogorov-Smirnov (KS) statistic. The KS statistic is calculated using the Python package *scipy* module `stats` method `ks_2samp`.

#### Principal component analysis

In Figure 4D and Figure 5D, simulated cell and nuclear shape features are transformed using Principal Component Analysis (PCA). PCA is calculated separately for Figure 4D and Figure 5D. For each PCA, data is combined across all conditions and *n* = 10 replicates, filtered to include only cells in S phase at *t* = 1 hour, and z-scored by feature. The PCA is performed using the Python package *scikit-learn* module `decomposition` class `PCA`.

#### ANOVA testing

Differences in cell and nuclear shape features are analyzed using two-way analysis of variance (ANOVA) for each combination of cell-media and cell-substrate adhesion. Different simulation sets use different parameters as the two factors **X1** and **X2**:

- Relative adhesion parameter simulations

**– X1** = Cell-cell adhesion
**– X2** = Subcellular adhesion
- Framework-specific parameter simulations

**– X1** = Monte Carlo steps
**– X2** = Temperature
- Spatial and temporal resolution parameter simulations

**– X1** = Spatial resolution
**– X2** = Temporal resolution

For each feature and combination of cell-media and cell-substrate adhesion, we perform two-way ANOVA with interaction with *α* = 0.05.

For cases where the interaction term is significant, we run Simple Main Effects testing using *α* = 0.05 with a Bonferroni correction for factor **X1** at each level of factor **X2**, and vice versa. If there are significant effects for factors with more than two levels, we then perform pairwise Tukey tests with *α* = 0.01 to identify which pairs of factor **X1** levels have significant difference at constant levels of factor **X2**, and vice versa.

For cases where the interaction term is not significant, we repeat the two-way ANOVA without the interaction term with *α* = 0.05. If there are significant effects for factors with more than two levels, we then perform pairwise Tukey tests with *α* = 0.01 to identify which pairs of factor **X1** levels have significant difference across all levels of factor **X2**, and vice versa.

ANOVA testing results are summarized in Supplementary Figures S2, S3, and S4. Cell and effect means as well as detailed ANOVA and Tukey test results are provided with the simulation data.

#### Cell and nuclear shape contours

Simulated cell and nuclear shapes contours are calculated from select slices (Figure 2 and 6, Supplementary Figure S1 and S5) or from projections (Figure 3, 4, and 5). For slice contours, the cell or nuclear voxels are converted to pixels by including only voxels for the selected slice. For projection contours, the cell or nuclear voxels are converted to pixels by merging along the projection axis (z axis for top view, x or y axis for side views). Contours are then taken as the border around each continuous region of pixels.

All contours are drawn in the following order: cell contour fill, nuclear contour fill, cell contour outline, nuclear contour outline.

## Acknowledgements

We wish to thank Allen Institute founders, Jody Allen & Paul G. Allen, for their vision, encouragement, and support. We thank the members of the Scientific Simulations Working Group, as well as all our colleagues at the Allen Institute for Cell Science, with whom we have had numerous, stimulating, and informative scientific discussions about this topic. N.B. is supported by the Washington Research Foundation and a National Science Foundation CAREER award CBET-1653315. J.A.T. is supported by the Howard Hughes Medical Institute (HHMI) award number CC34670. This article is subject to HHMI’s Open Access to Publications policy. HHMI laboratory heads have previously granted a nonexclusive CC BY 4.0 license to the public and a sublicensable license to HHMI in their research articles. Pursuant to those licenses, the author-accepted manuscript of this article can be made freely available under a CC BY 4.0 license immediately upon publication.

## Author contributions

J.S.Y. Conceptualization, Software, Formal analysis, Writing - Original Draft, Writing - Review & Editing, Visualization; B.L. Writing - Review & Editing; S.R. Conceptualization, Writing - Review & Editing; J.A.T. Conceptualization, Writing - Review & Editing; N.B. Conceptualization, Writing - Review & Editing, Visualization, Supervision; G.T.J. Conceptualization, Writing - Review & Editing, Visualization, Supervision.

## Data availability statement

Simulation data generated in this study are available as Quilt packages. The Hamiltonian terms dataset is available at https://open.quiltdata.com/b/allencell/packages/aics/hybrid_model_simulations_hamiltonian_terms. The relative adhesion parameter dataset is available at https://open.quiltdata.com/b/allencell/packages/aics/hybrid_model_simulations_relative_adhesion. The Cellular Potts Model framework-specific parameter dataset is available at https://open.quiltdata.com/b/allencell/packages/aics/hybrid_model_simulations_potts_parameters. The spatial and temporal resolution parameter dataset is available at https://open.quiltdata.com/b/allencell/packages/aics/hybrid_model_simulations_spatial_temporal_resolution. The full monolayer simulations data is available at https://open.quiltdata.com/b/allencell/packages/aics/hybrid_model_simulations_full_monolayer.

## Code availability statement

All source code for the model (ARCADE v3.1.4) is publicly available on GitHub at https://github.com/bagherilab/ARCADE. Pipeline code for running and analyzing simulations is publicly available on GitHub at https://github.com/allen-cell-animated/cell-abm-pipeline/.

## Declaration of generative AI and AI-assisted technologies in the writing process

During the preparation of this work the authors used ChatGPT-4 to reduce word count and streamline verbose take home messages. After using this tool, the authors reviewed and edited the content as needed and take full responsibility for the content of the publication.

## Supplemental figures and tables

**Figure S1.**
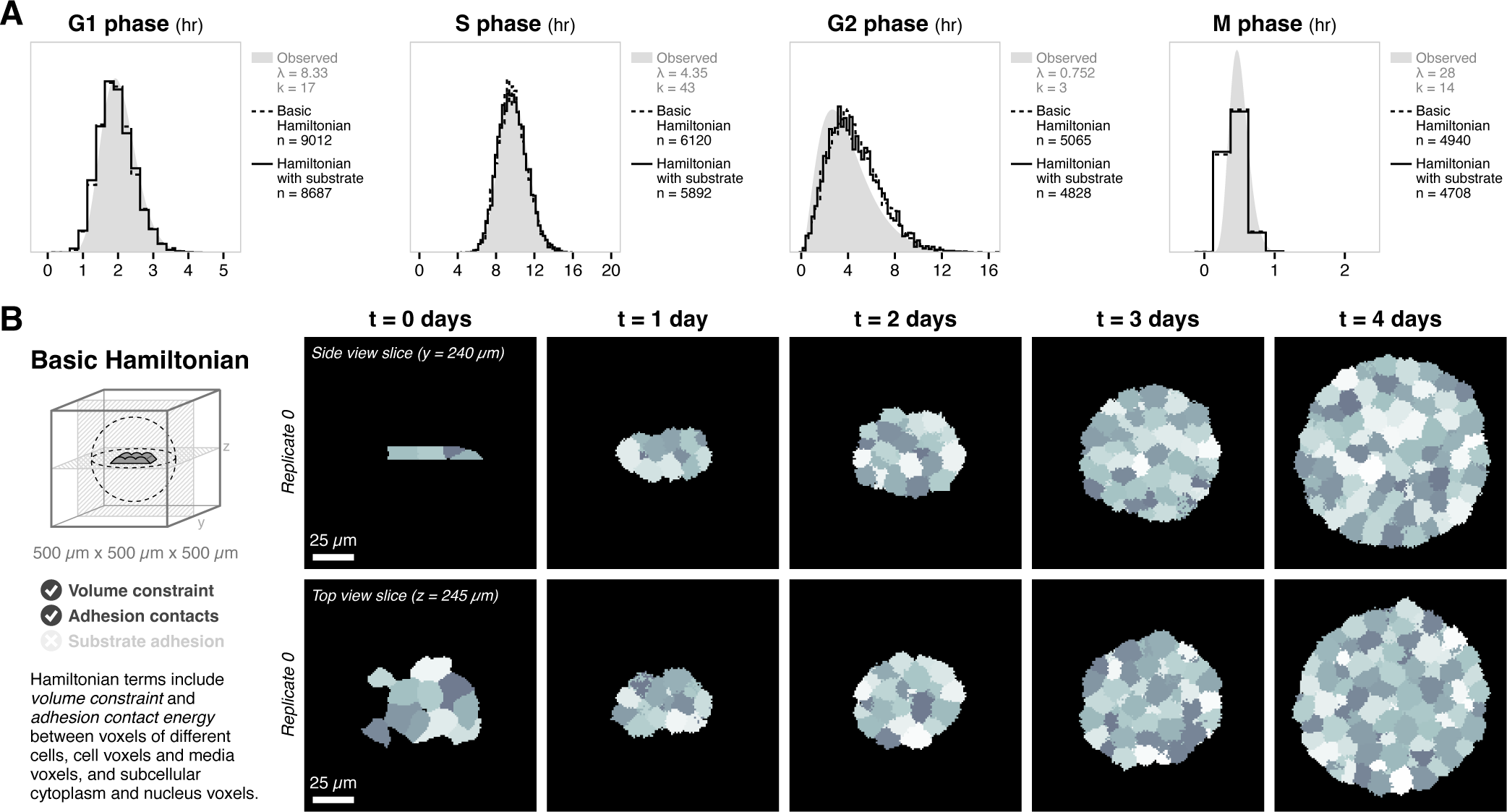
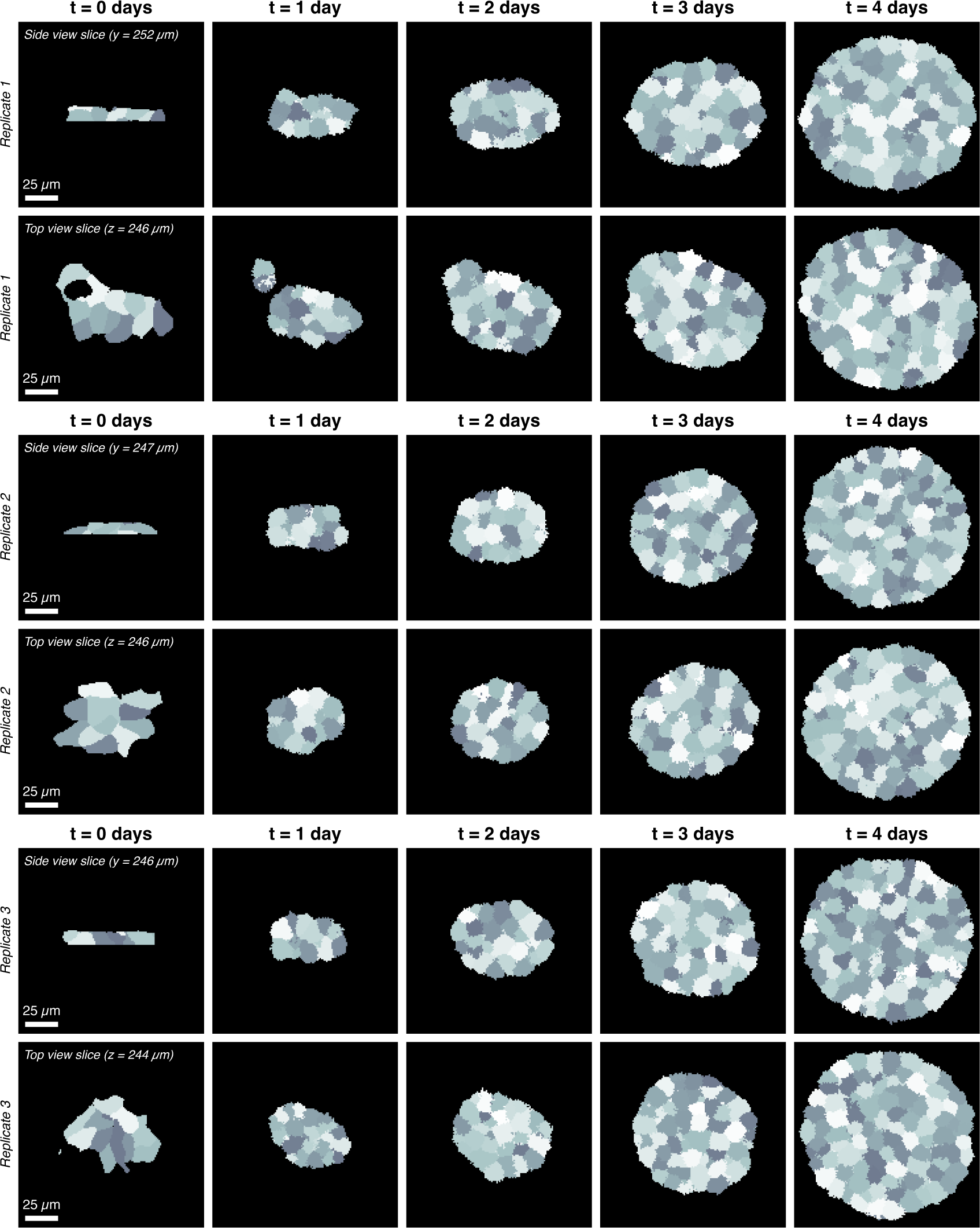
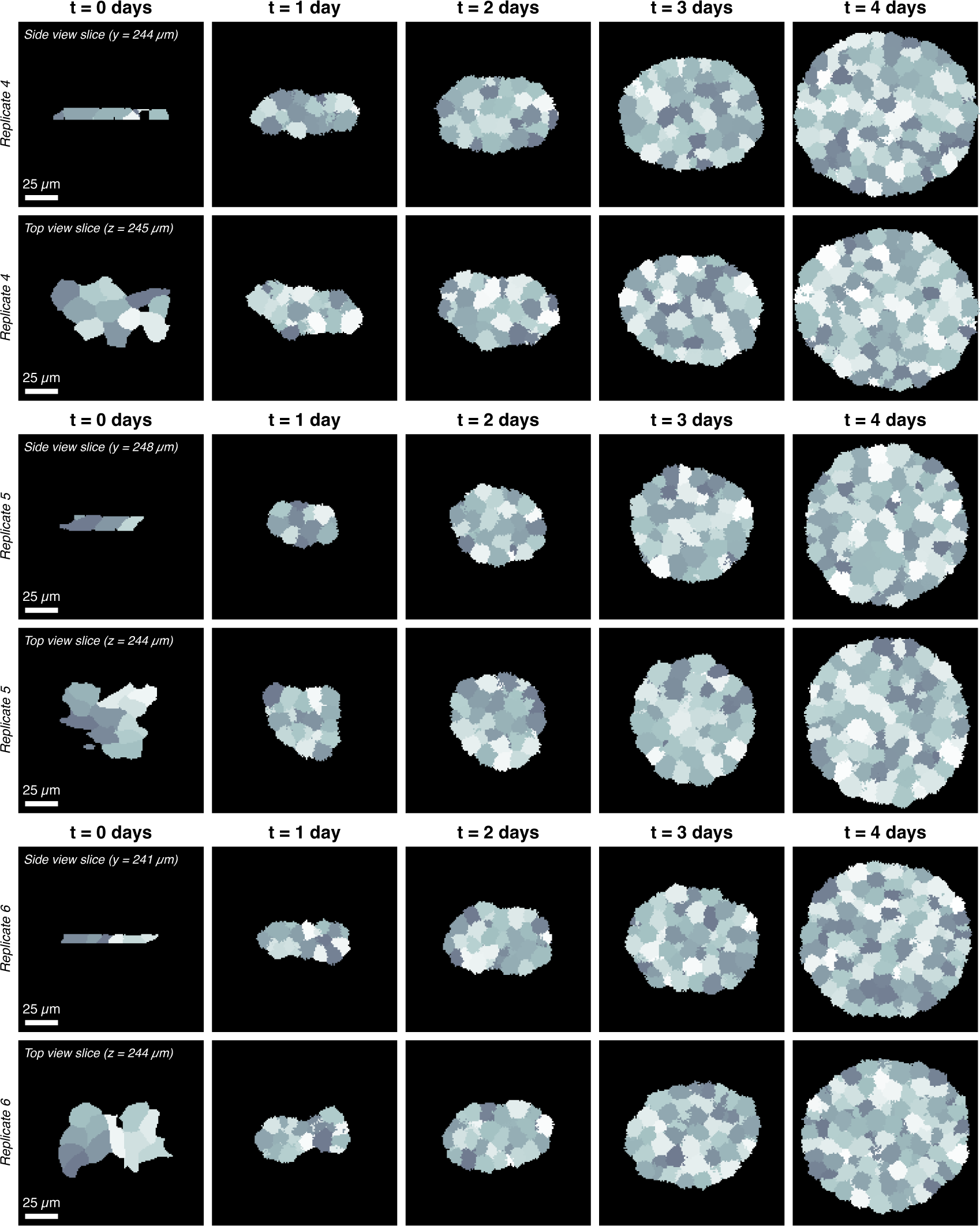
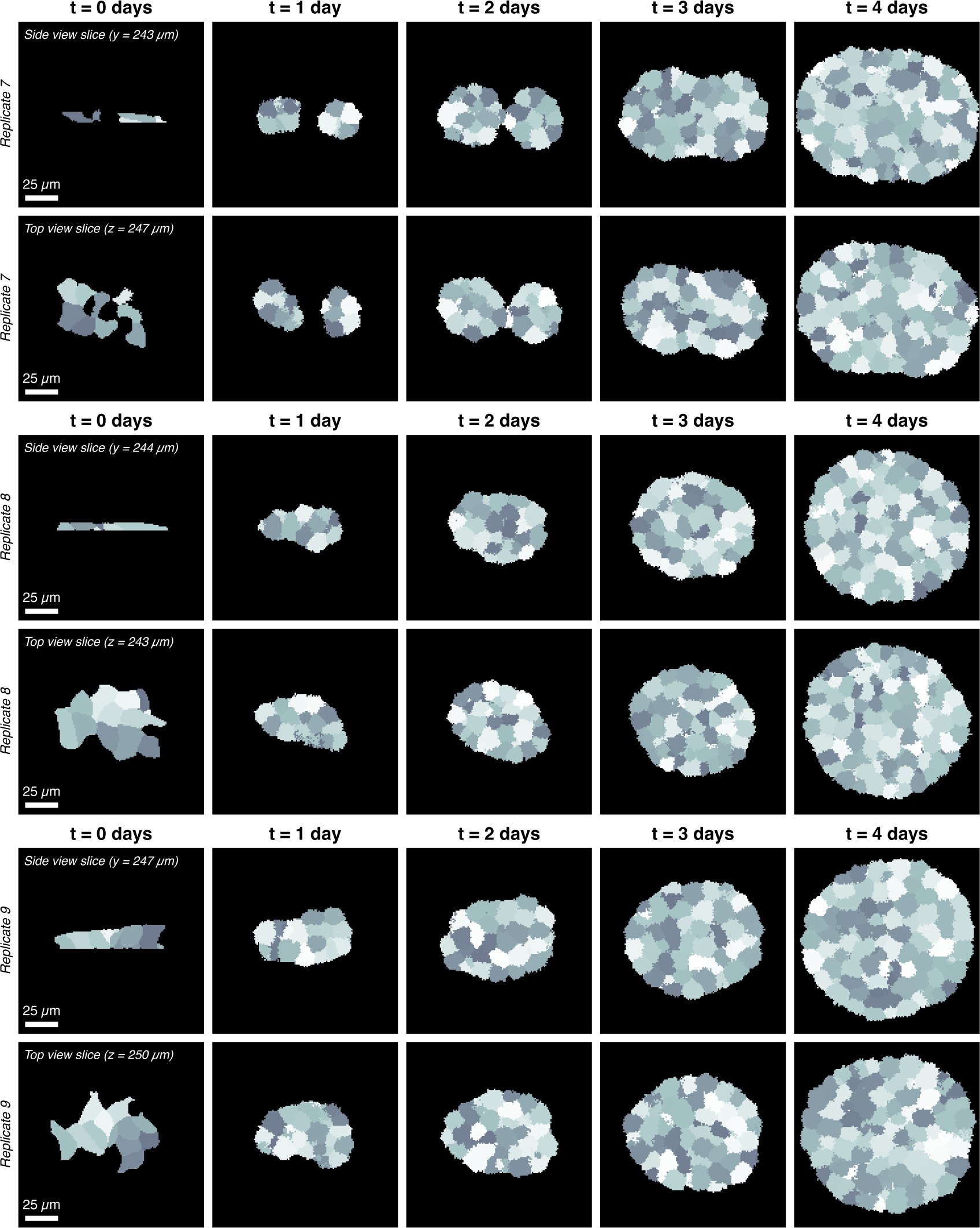
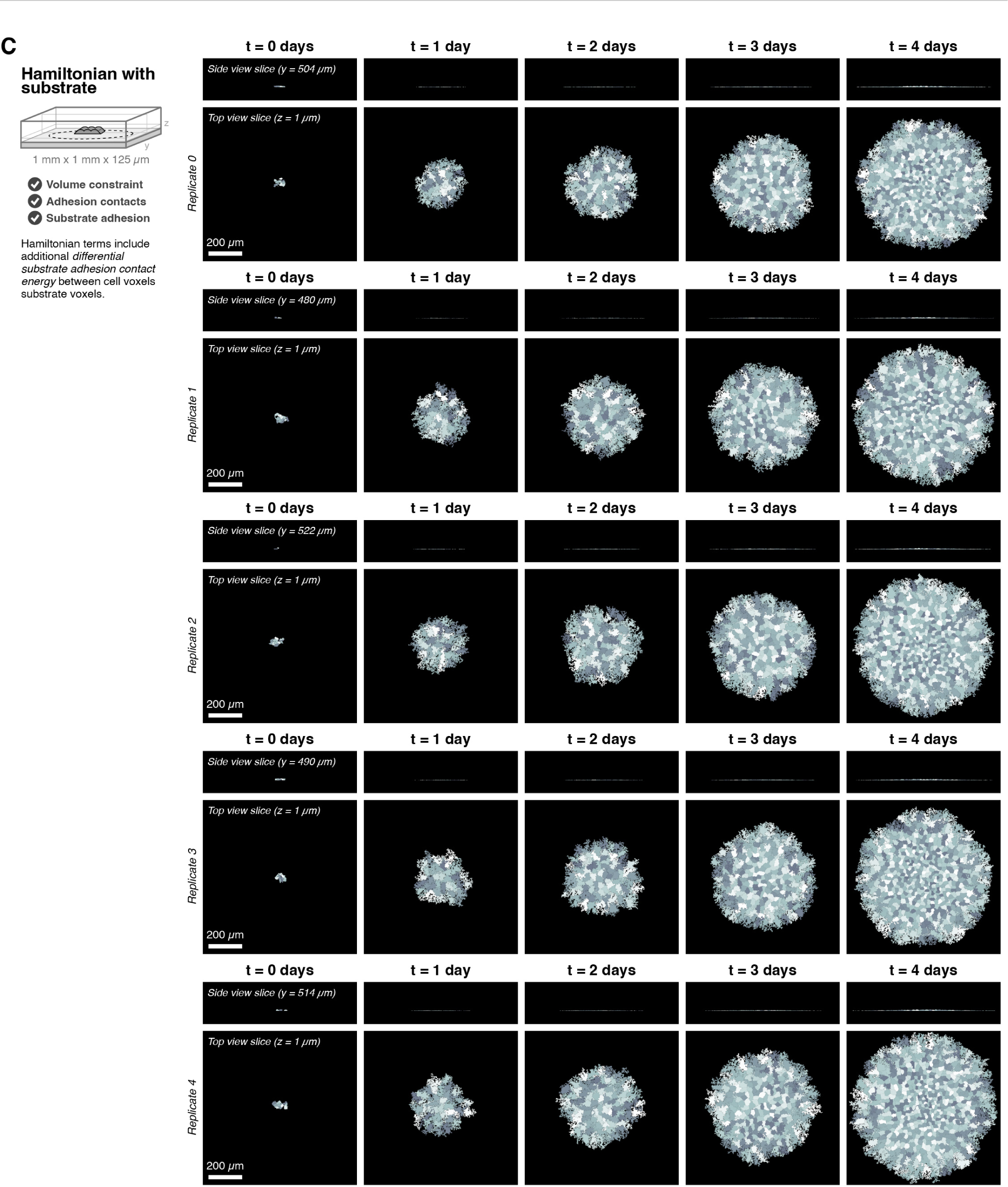
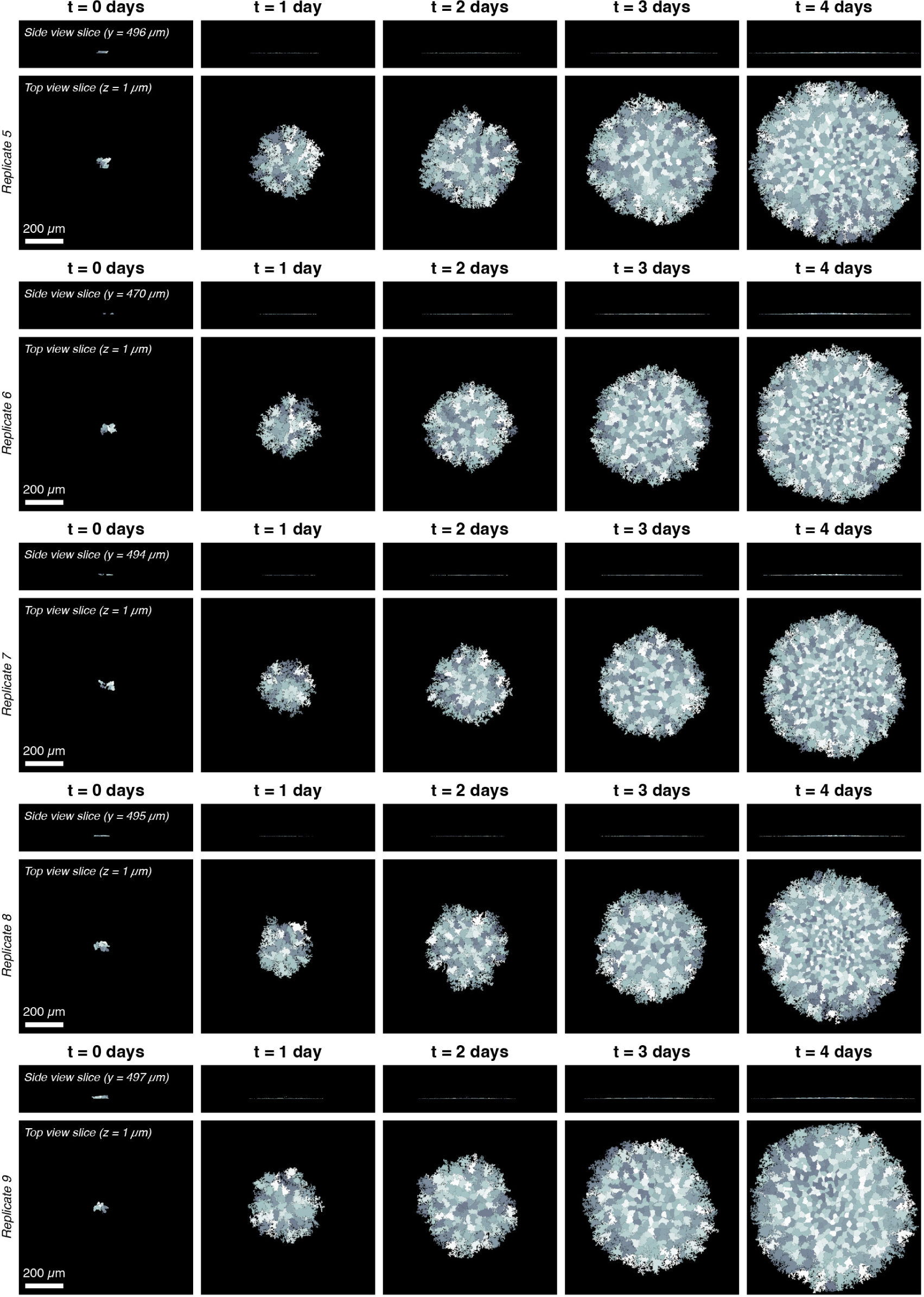
Iterative simulations of hybrid model with different Hamiltonian terms changes emergent colony shape from spherical to flat. *Related to Figure 2*. (**A**) Distributions of cell cycle phase duration across all *n* = 10 replicates for both the basic Hamiltonian and Hamiltonian with substrate conditions. Reference distribution in gray is an Erlang distribution with the specified *λ* and *k* parameters. (**B**) Cell and colony shapes for select time points across replicates for the basic Hamiltonian condition. Hamiltonian includes volume constraint and adhesion contact energies (cell-cell, cell-media, and subcellular). Diagram shows gray cells initialized at *t* = 0 days in the center of a 500 *µm* x 500 *µm* x 500 *µm* simulation environment and the dotted black outline shows approximate colony shape at *t* = 4 days. Gray hatched slices indicate the location of the top view (*z* = 245 *µm*) and the side view (*y* = 240 *µm*) shown in the simulation snapshots. Gray shading is used to differentiate between cells. (**C**) Cell and colony shapes for select time points across replicates for the Hamiltonian with substrate condition. Hamiltonian includes volume constraint, adhesion contact energies (cell-cell, cell-media, and subcellular), and differential cell-substrate adhesion. Diagram shows gray cells initialized at *t* = 0 days in the bottom of a 1 mm x 1 mm x 125 *µm* simulation environment and the dotted black outline shows approximate colony shape at *t* = 4 days. Gray hatched slices indicate the location of the top view (*z* = 1 *µm*) and the side view (*y* = 504 *µm*) shown in the simulation snapshots. Gray shading is used to differentiate between cells.

**Figure S2.**
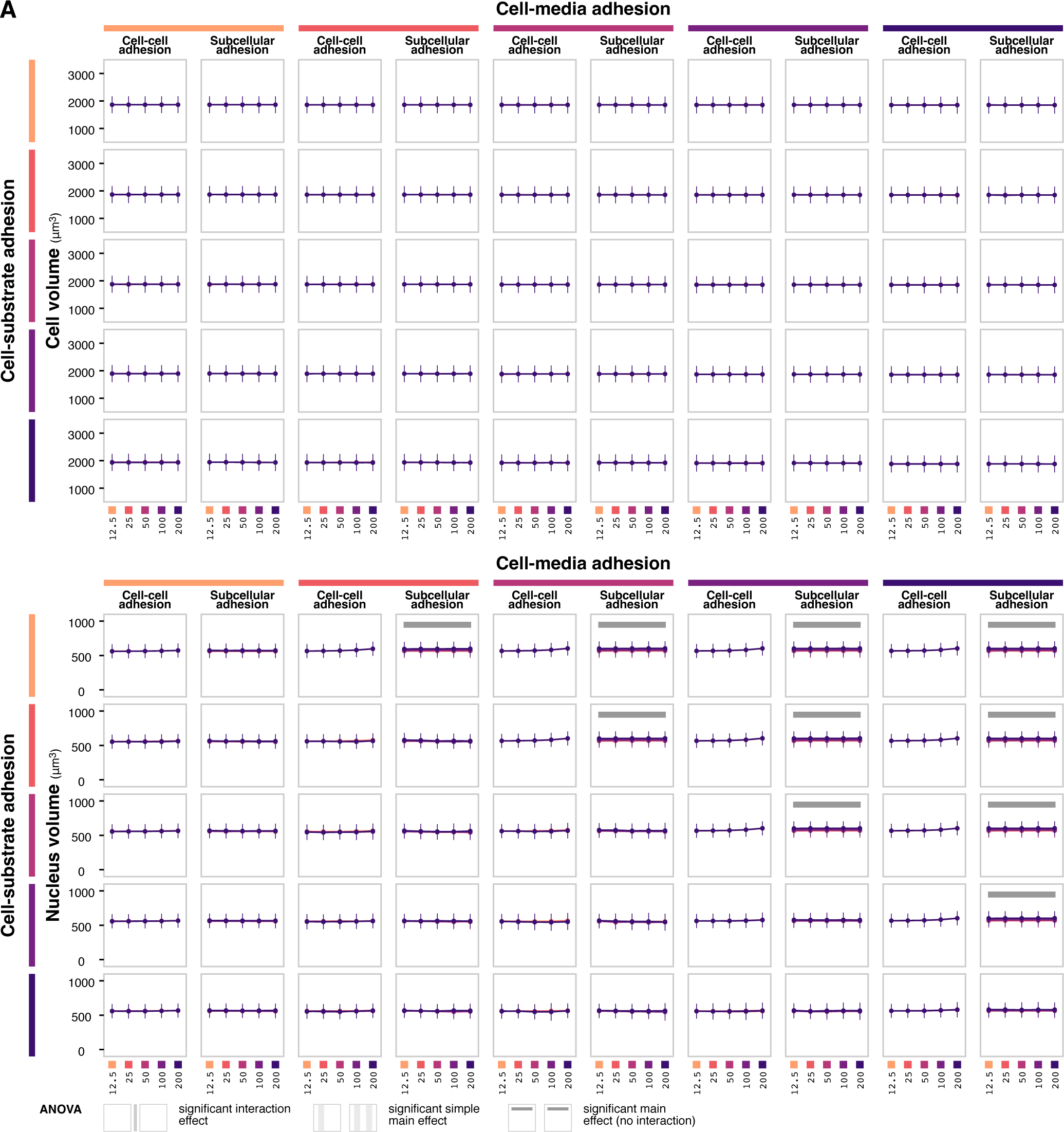
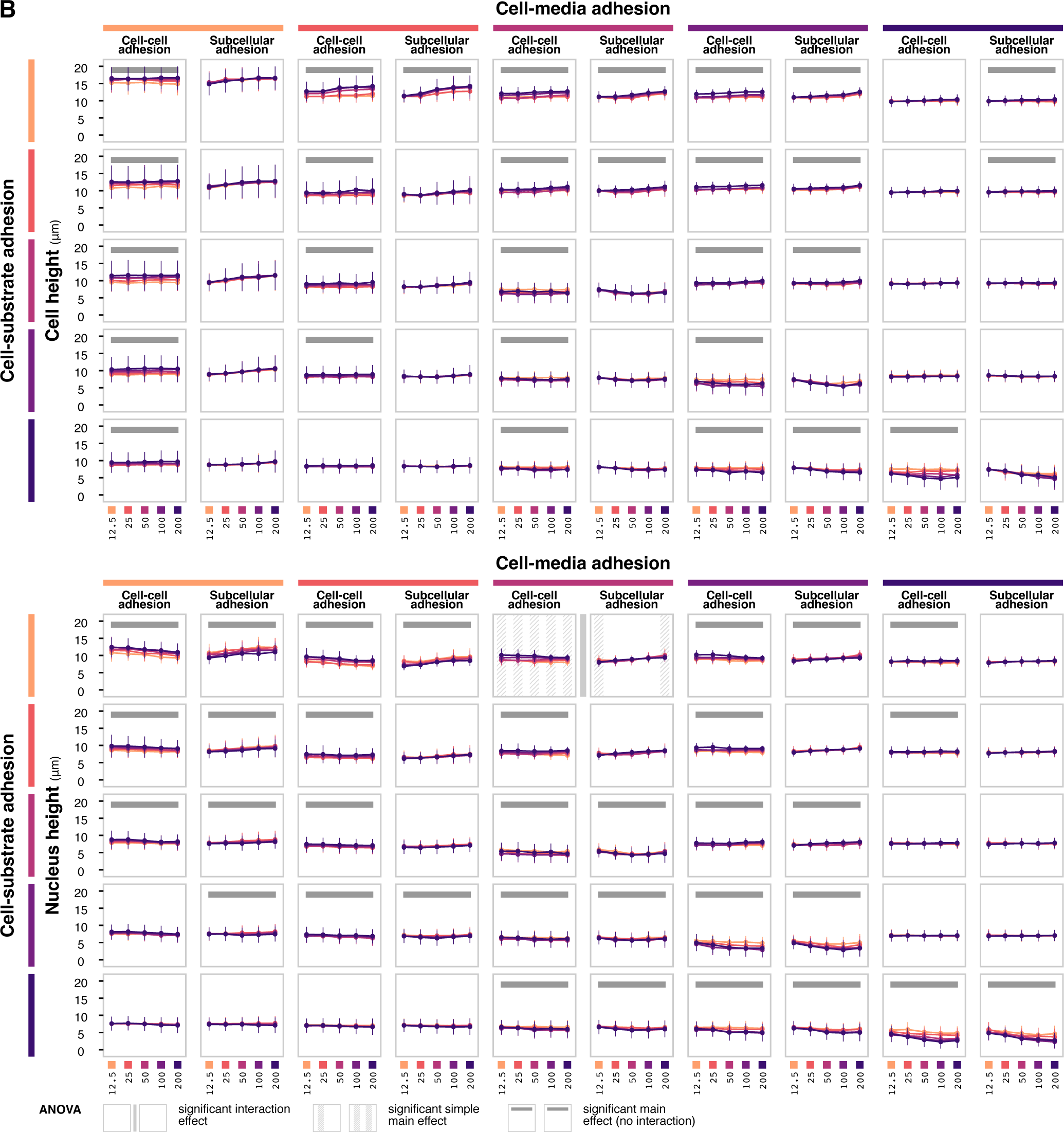
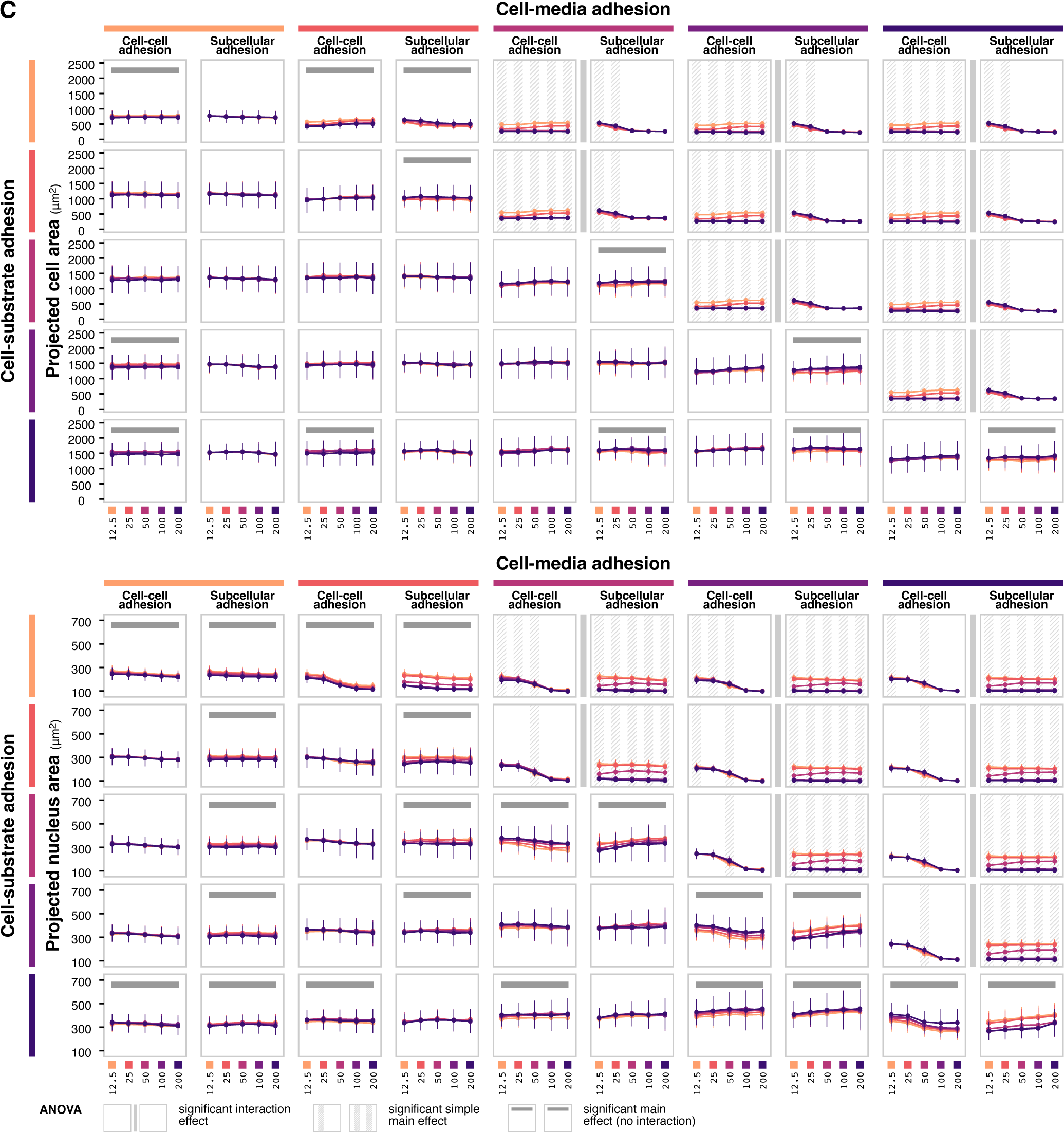
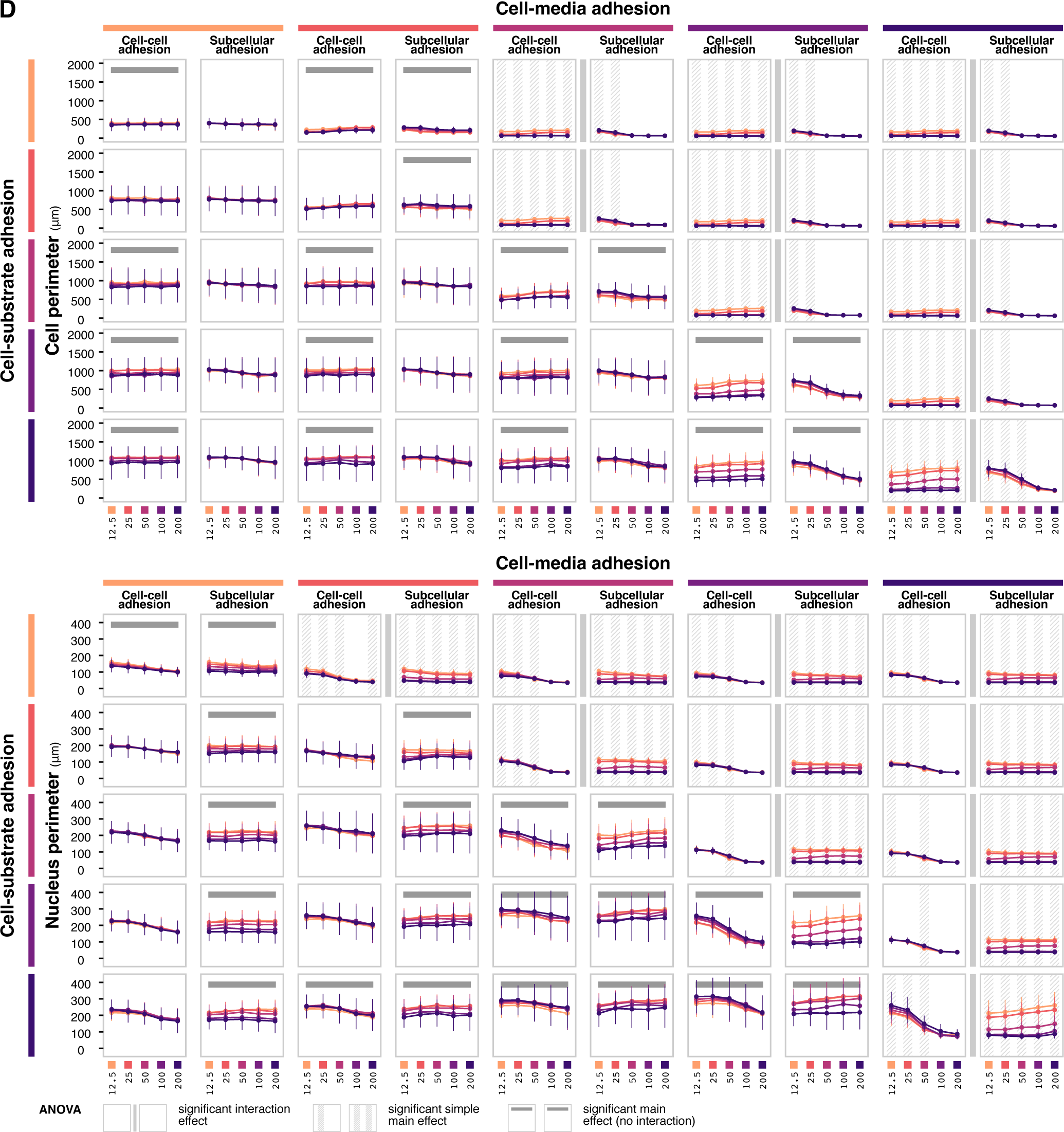
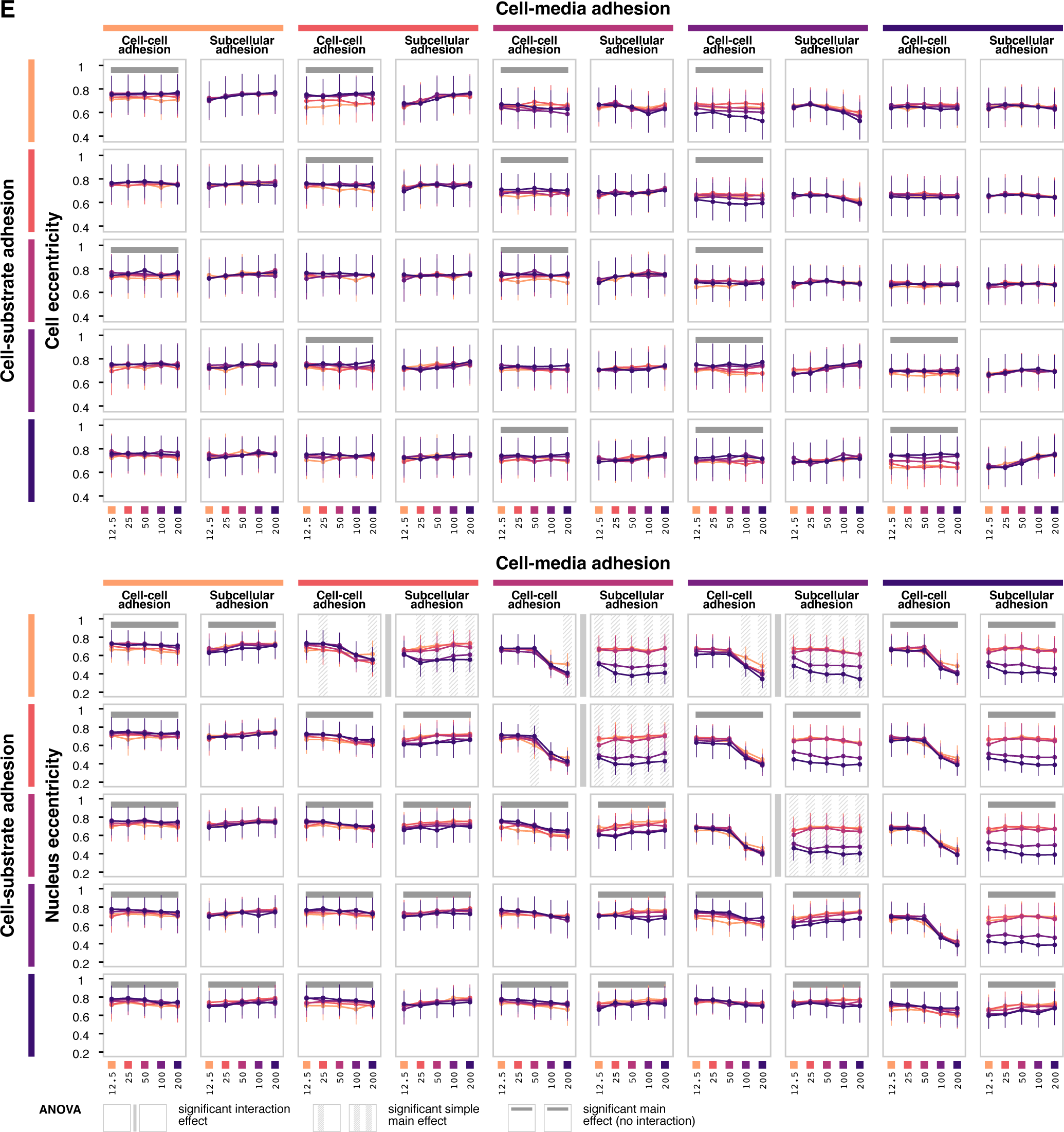
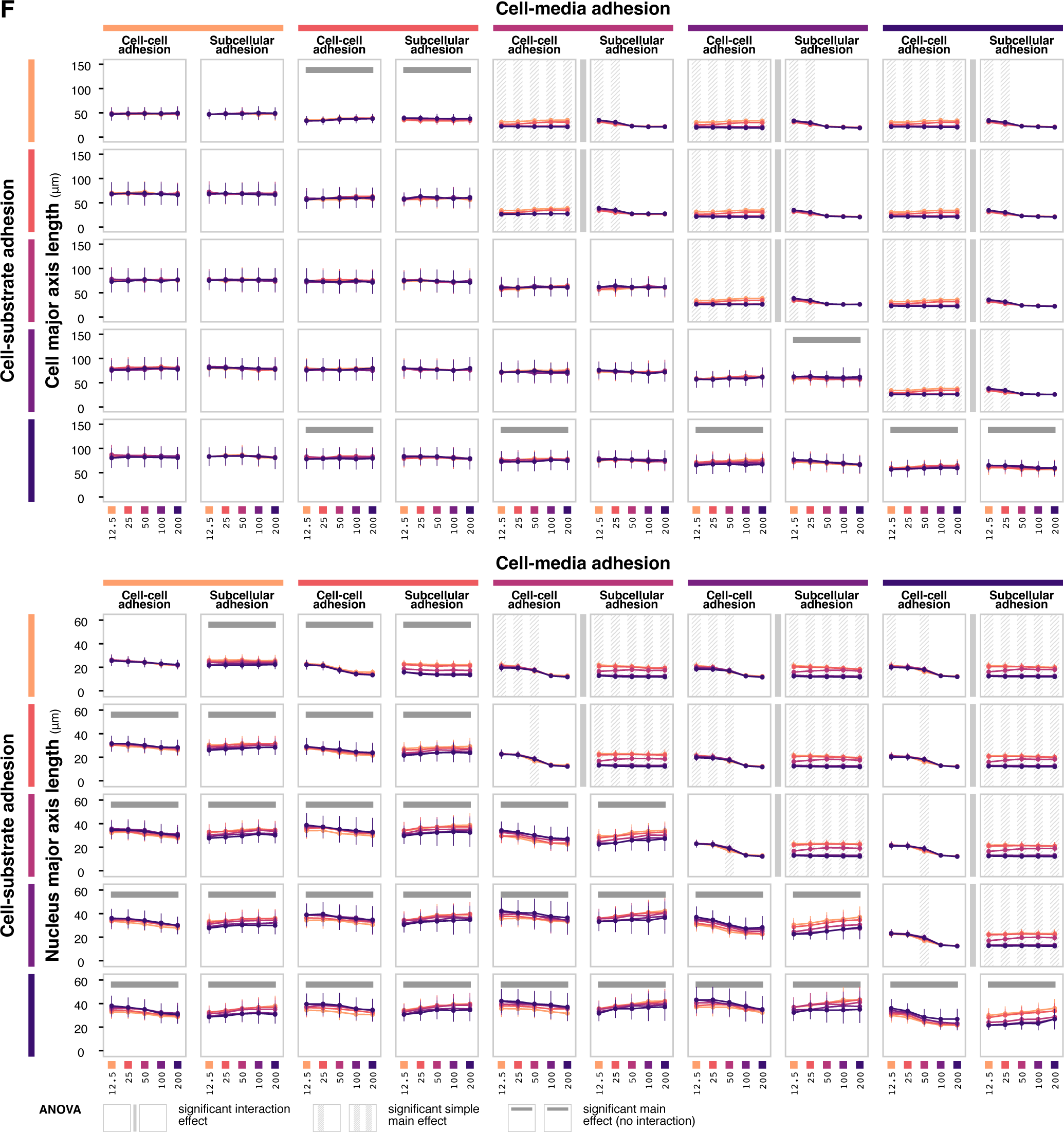
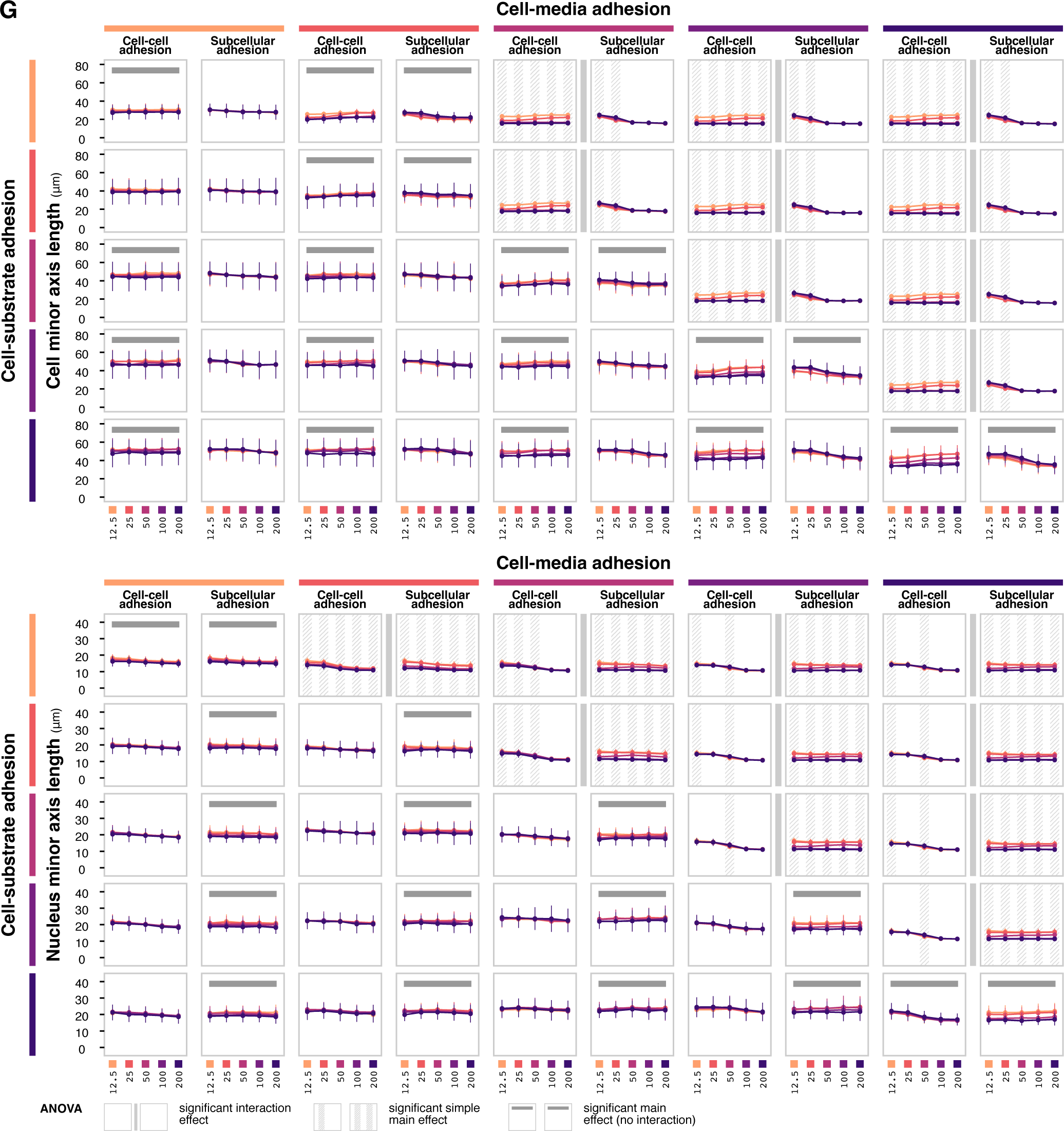
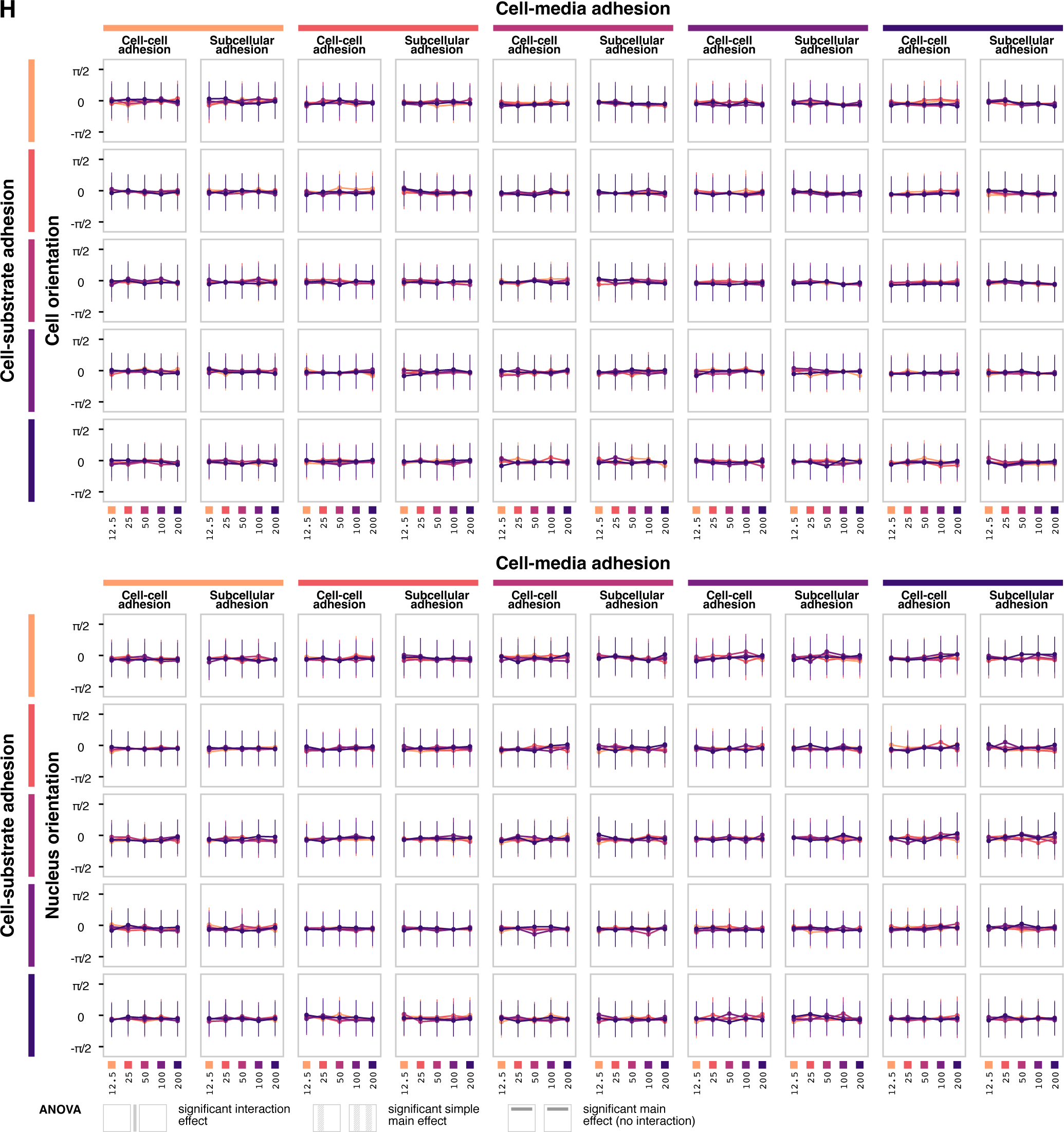
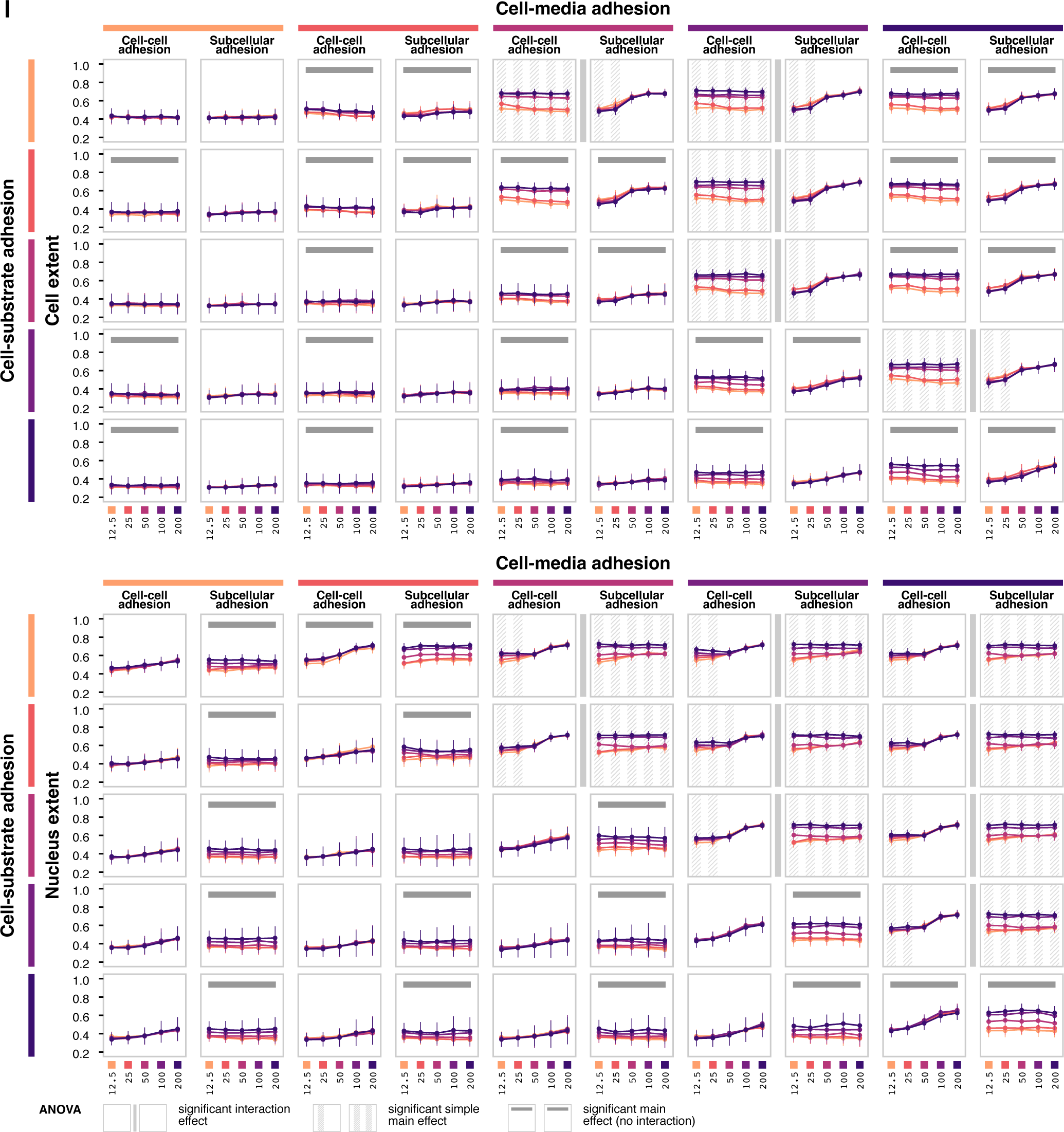
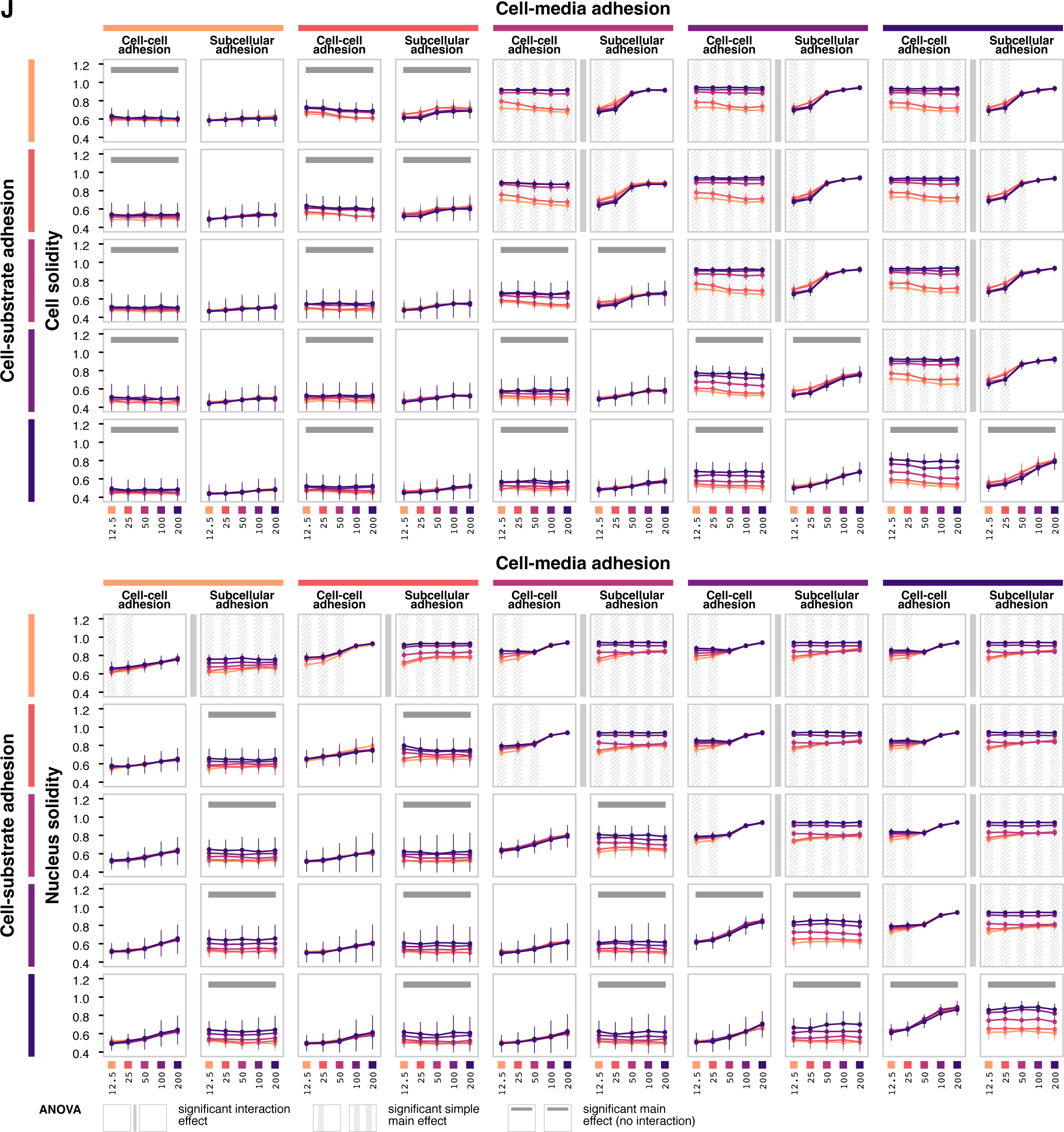
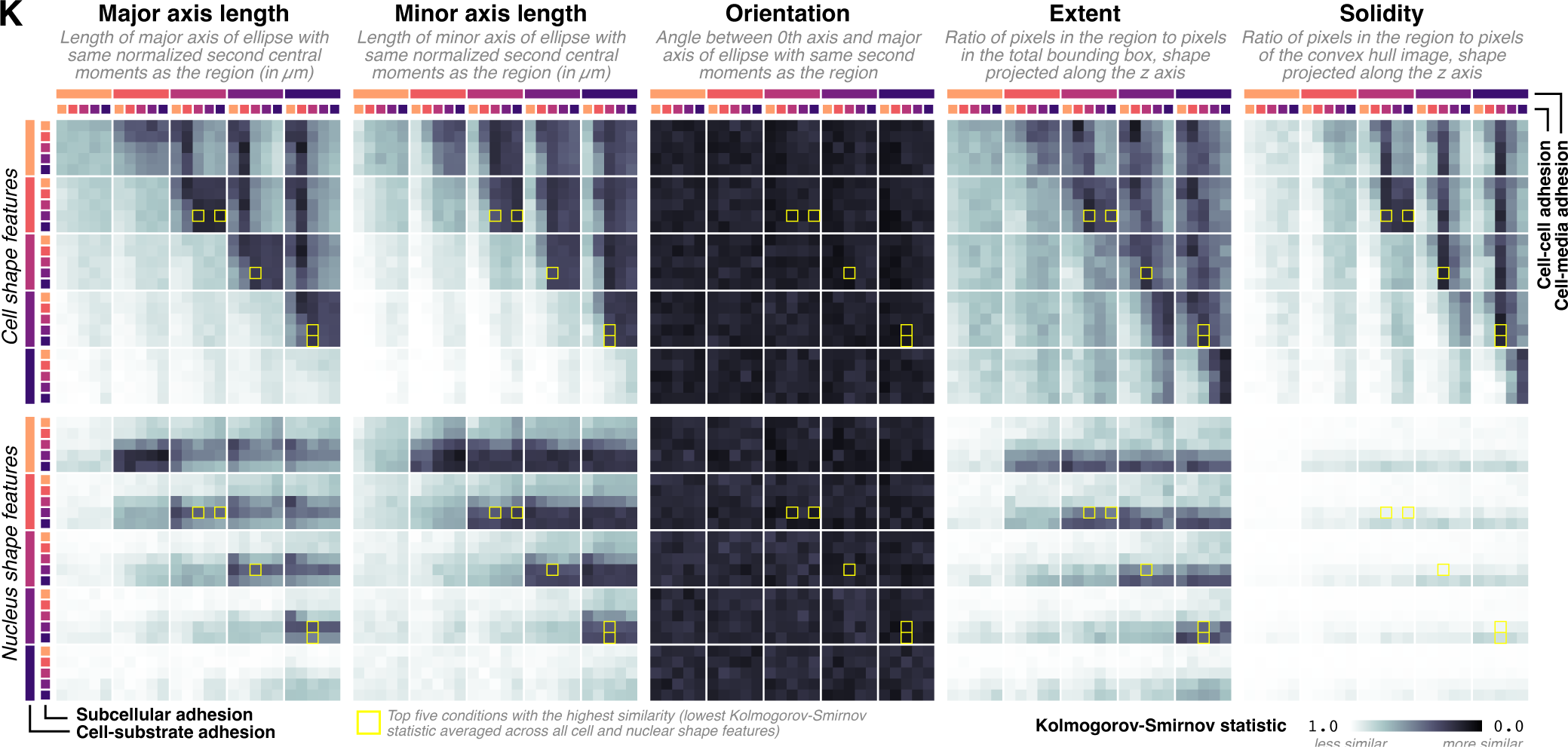
Relative adhesion parameter simulations. *Related to Figure 3*. ANOVA interval plots for cell and nuclear (**A**) volume, (**B**) height, (**C**) area, (**D**) perimeter, (**E**) eccentricity, (**F**) major axis length, (**G**) minor axis length, (**H**) orientation, (**I**) extent, and (**J**) solidity. ANOVA is performed separately for each combination of cell-media and cell-substrate adhesion. Each data point shows the mean value of the feature at *t* = 1 hour for the given level of cell-cell adhesion or subcellular adhesion, holding the other condition constant. Error bars show standard deviation across *n* = 10 replicates. Annotations indicate statistically significant interaction effects, simple main effects (if interaction is significant) and main effects (if interaction is not significant) at *α* = 0.05. (**K**) Heatmaps showing Kolmogorov-Smirnov statistic calculated between the simulated and observed distributions of different features under different simulation conditions. Darker colors indicate lower Kolmogorov-Smirnov statistic values, which indicates the simulated distribution is more similar to the observed distribution. Conditions producing the highest average similarity are marked by yellow borders (corresponding to the yellow highlights in Panel B).

**Figure S3.**
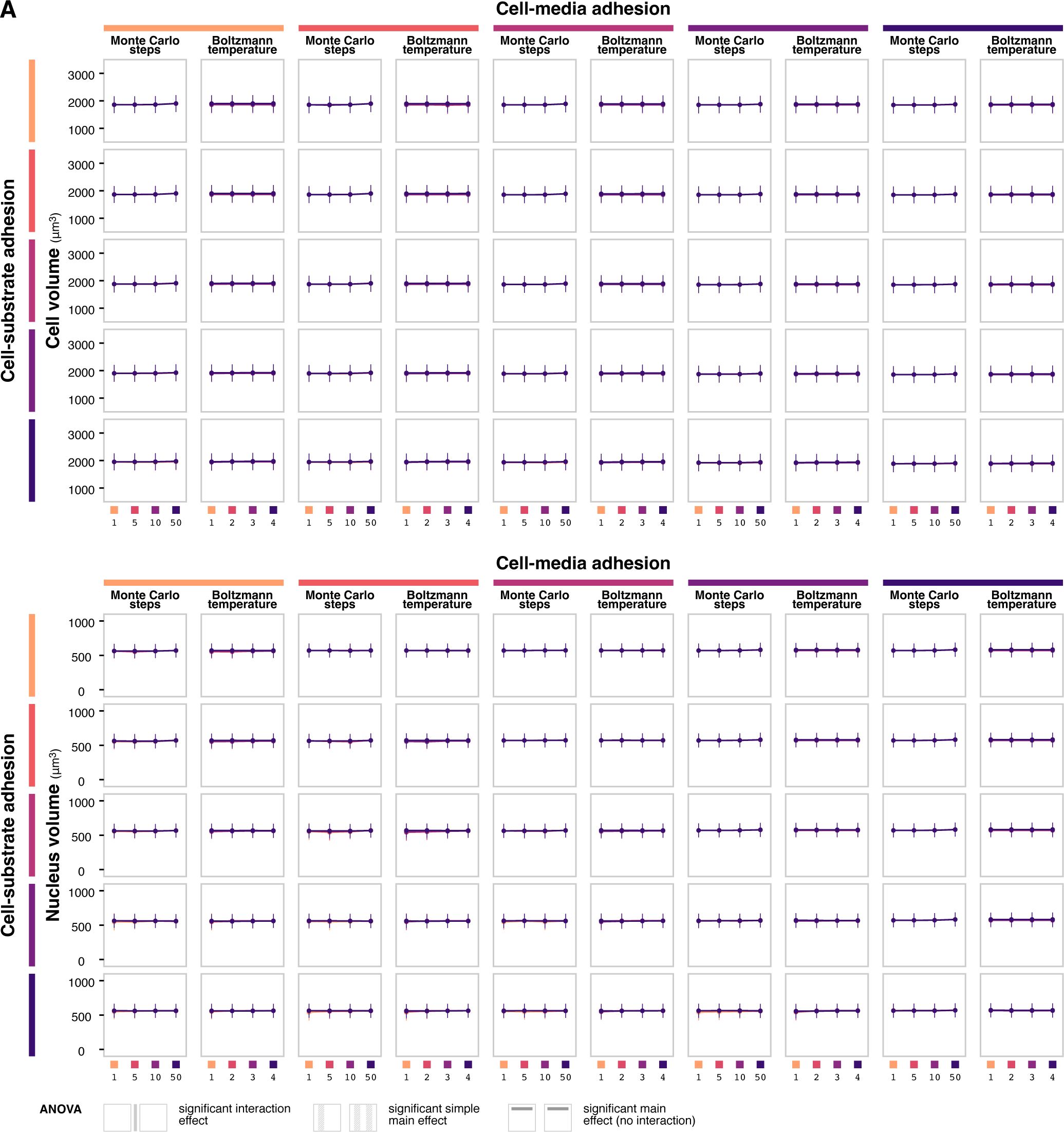
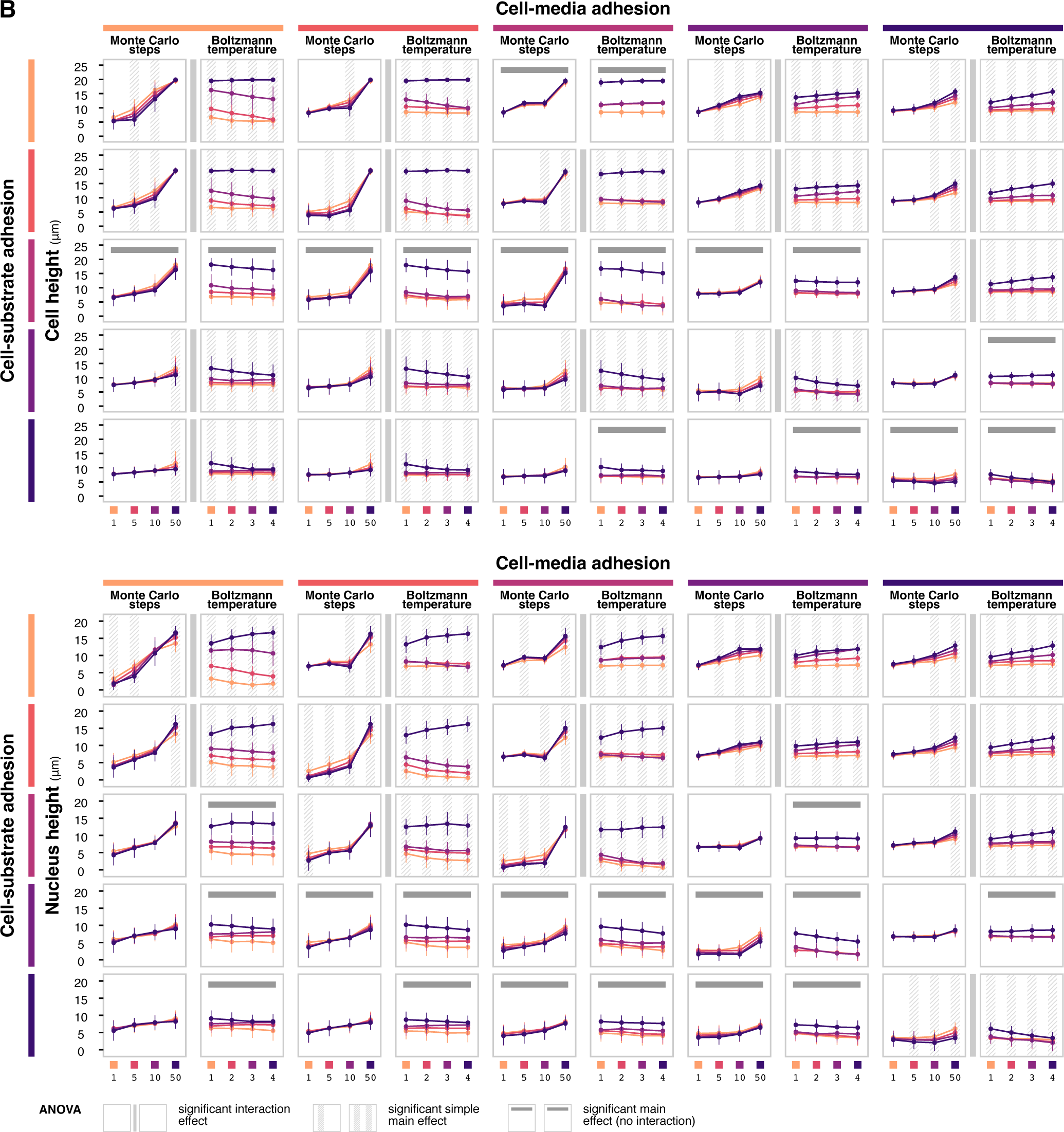
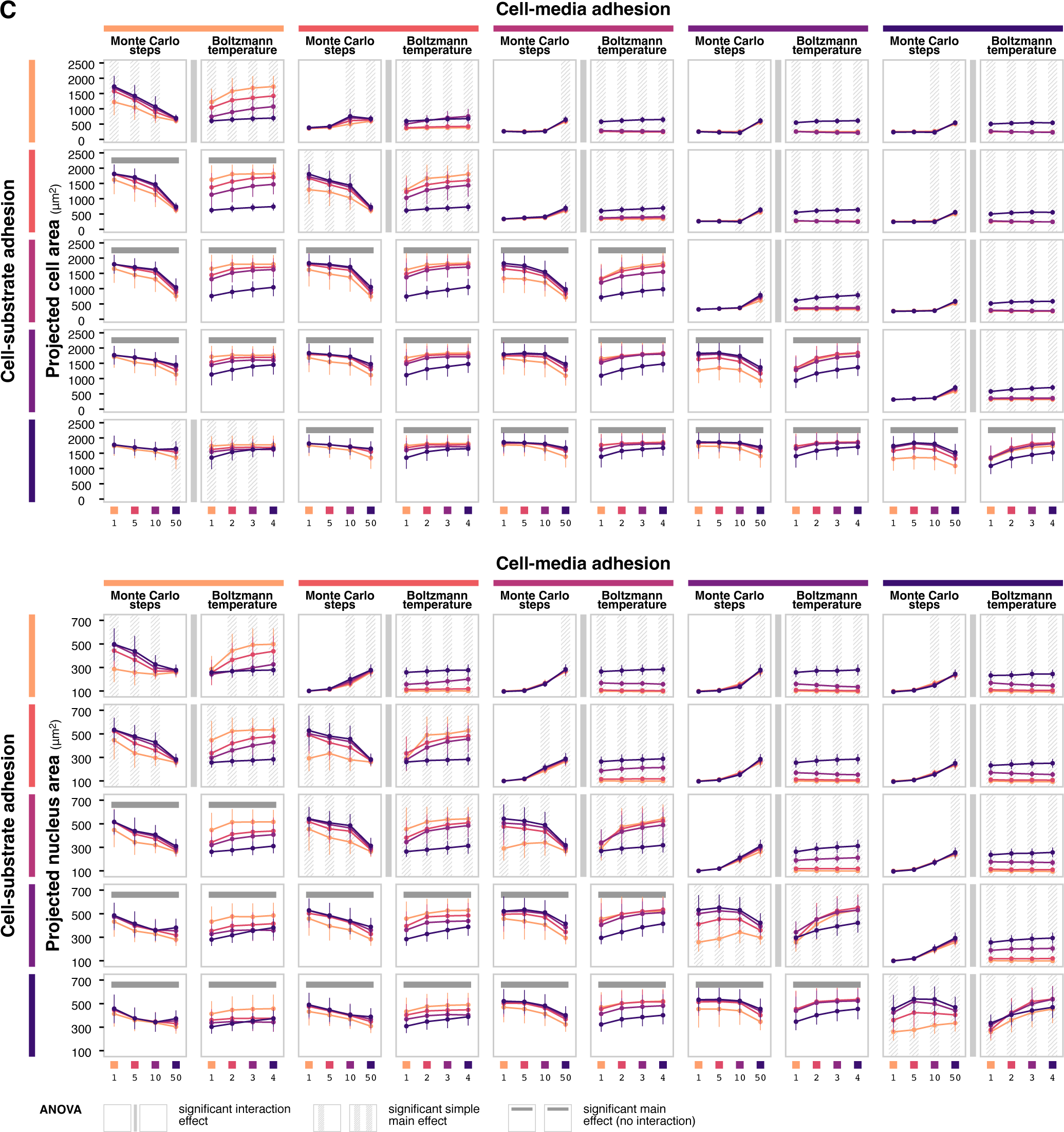
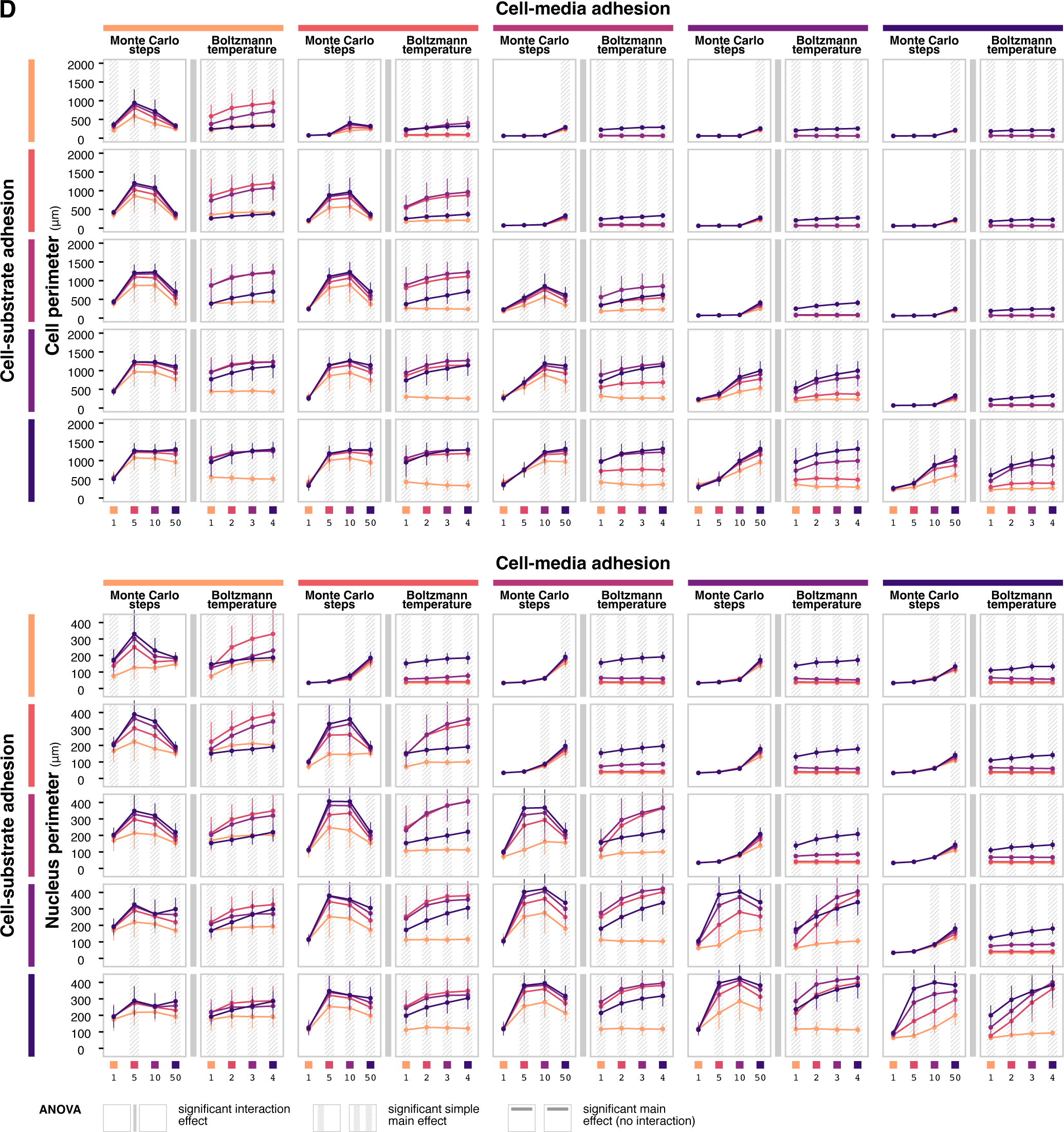
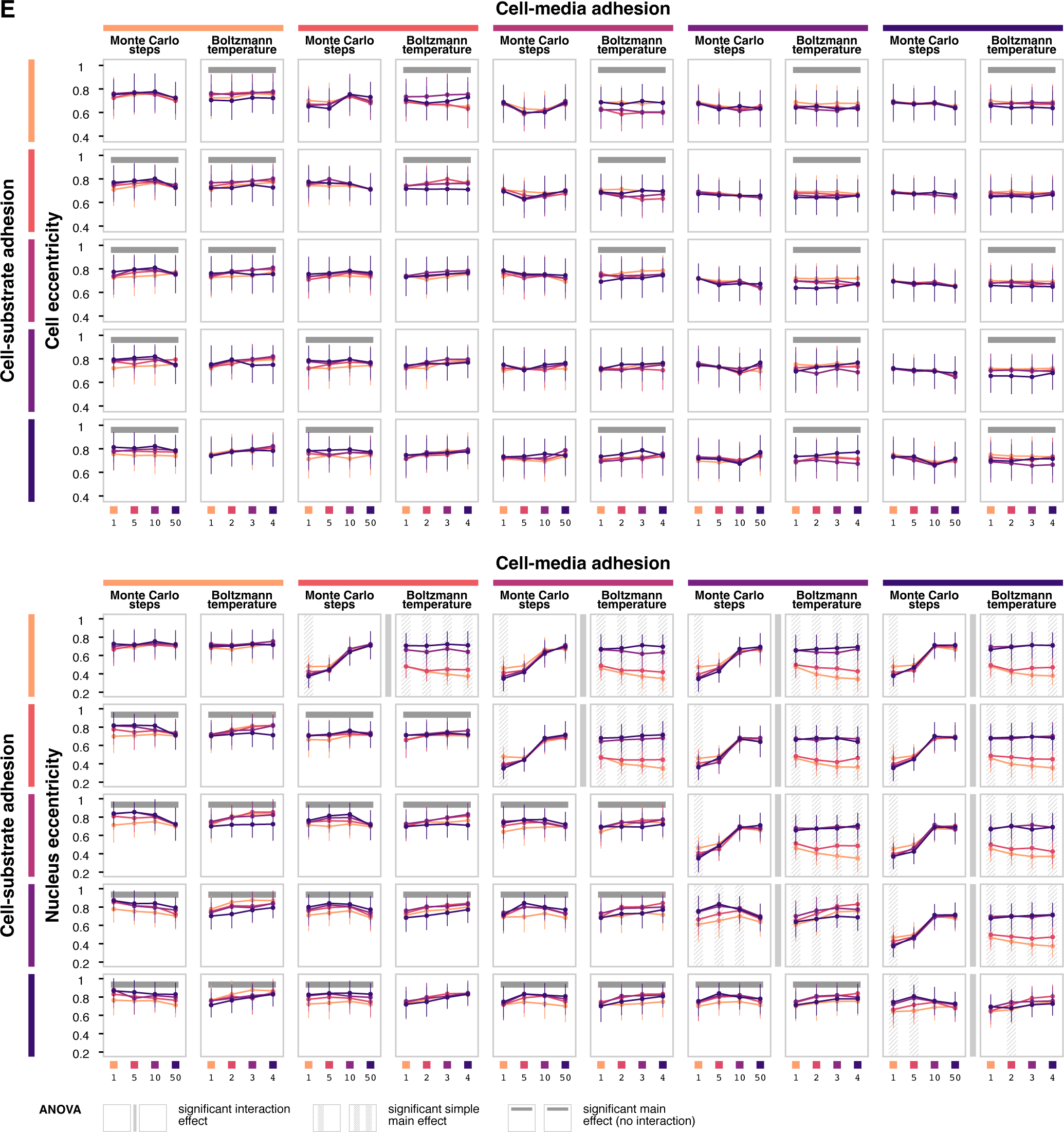
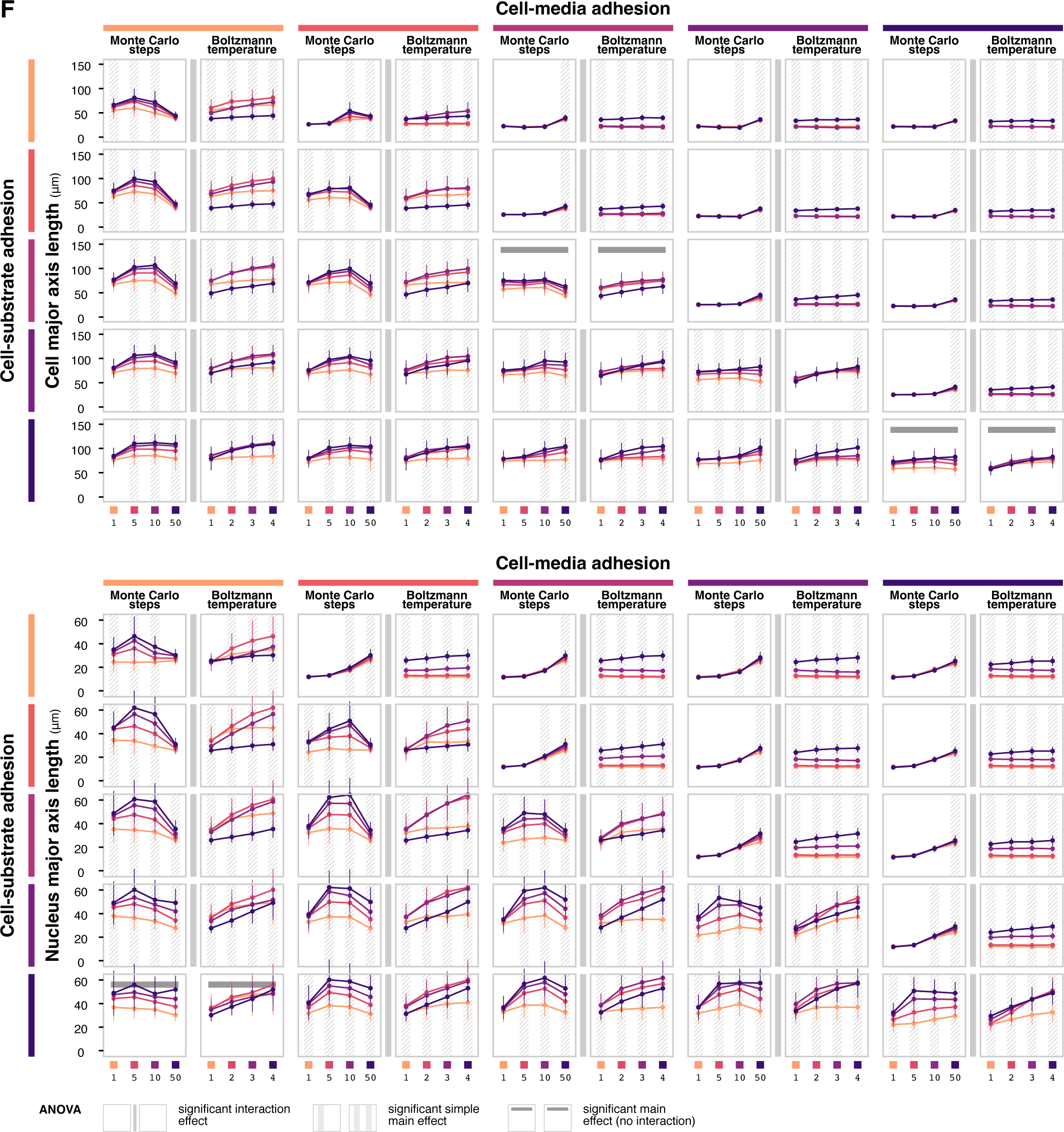
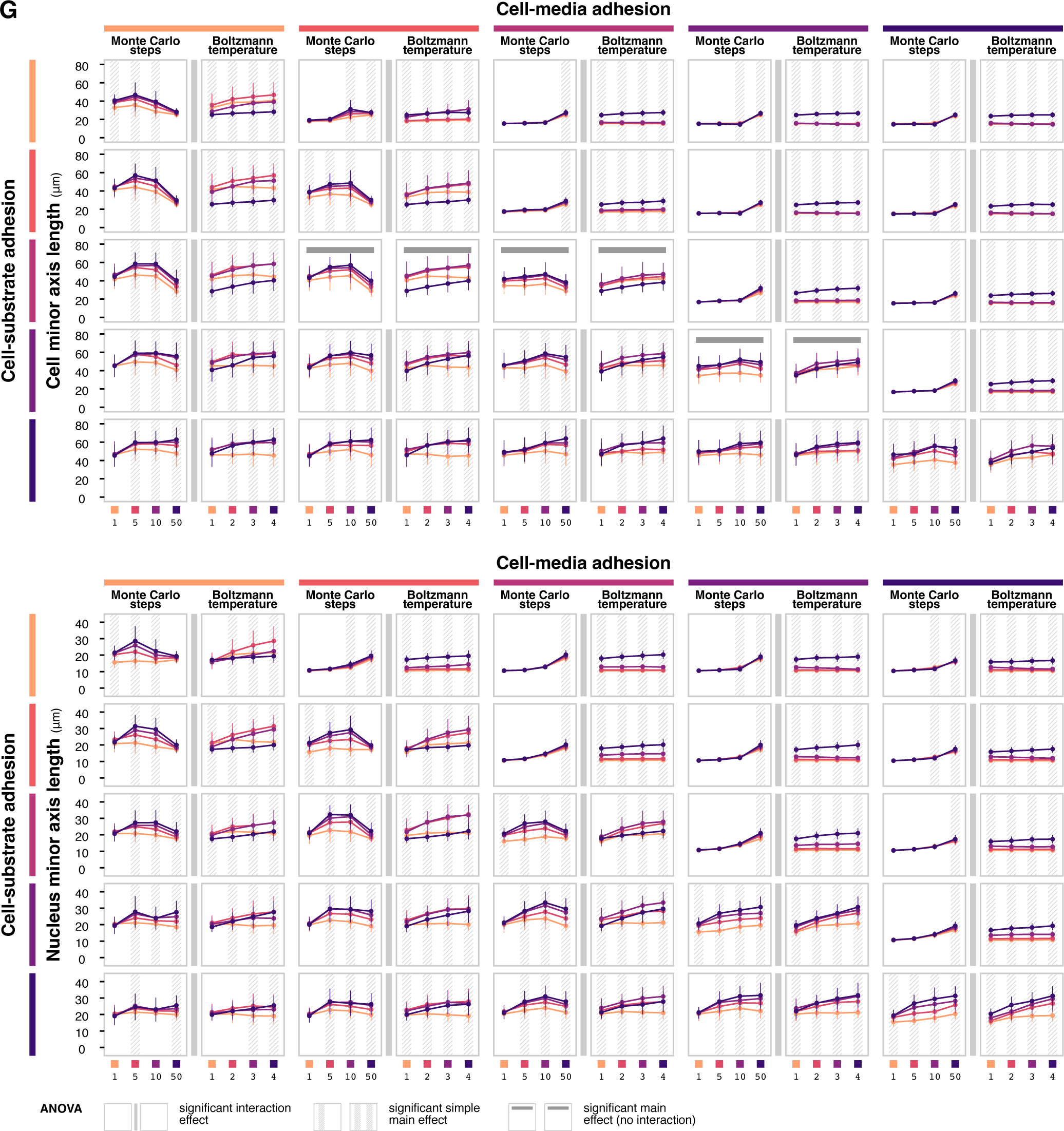
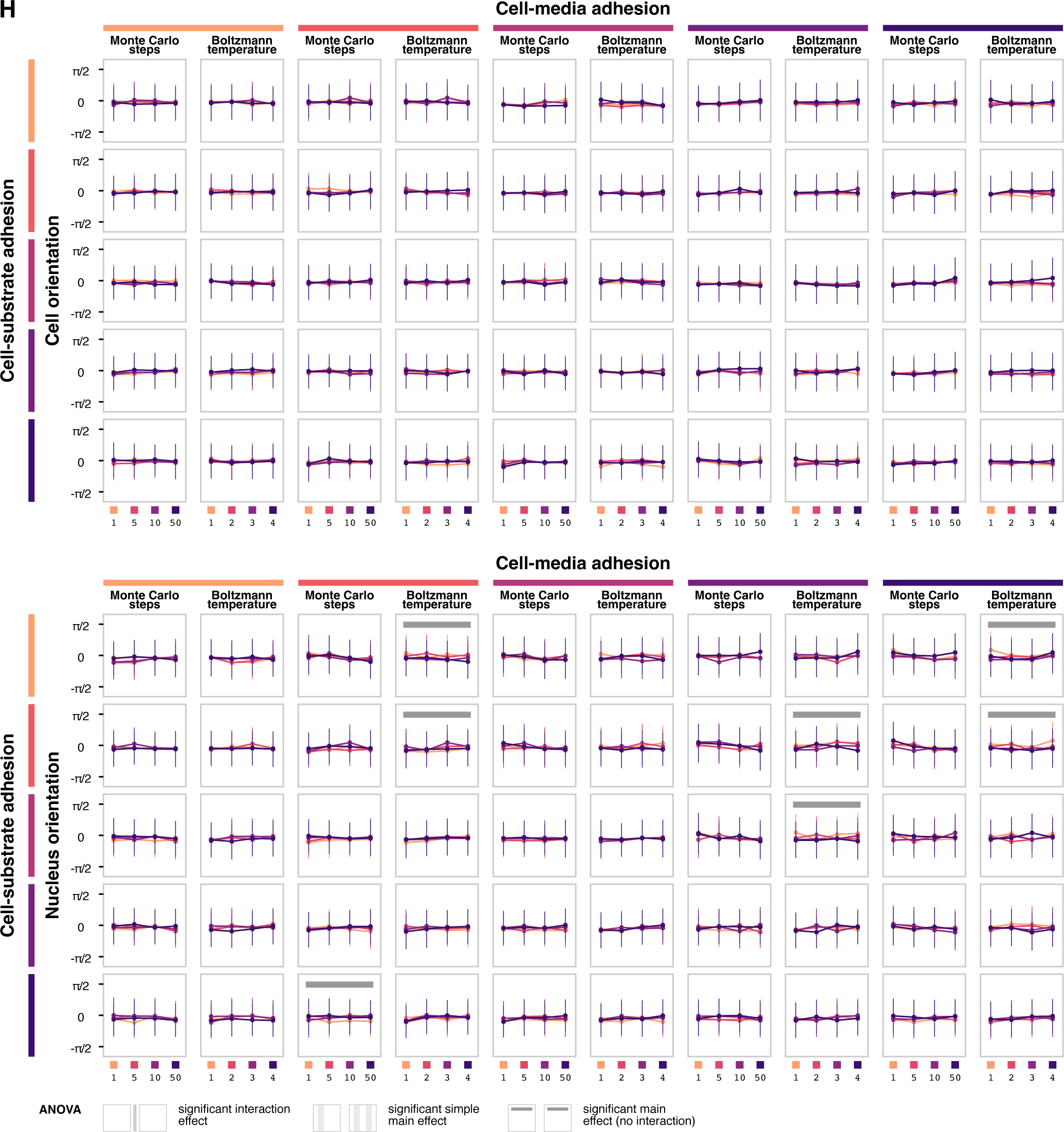
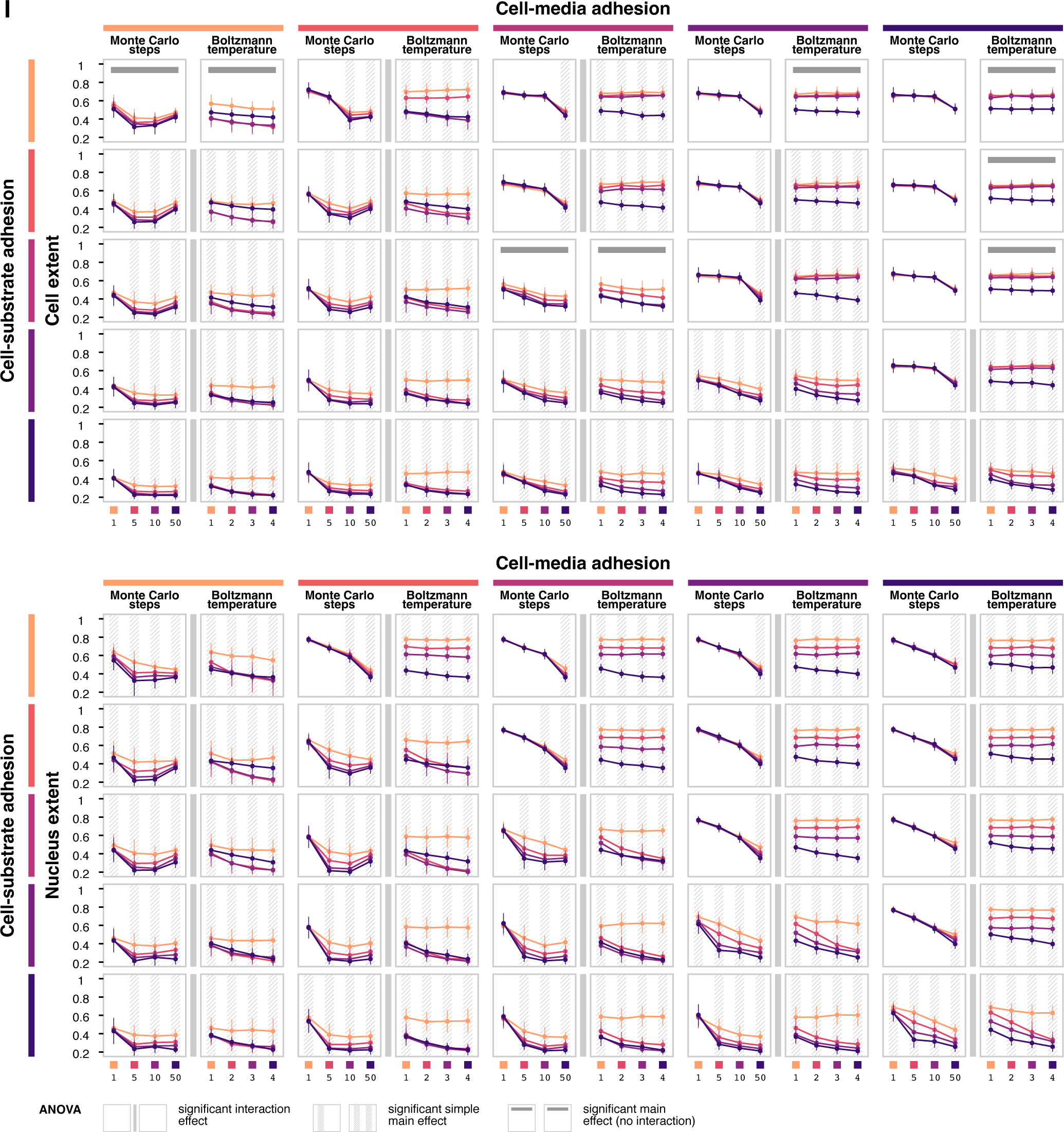
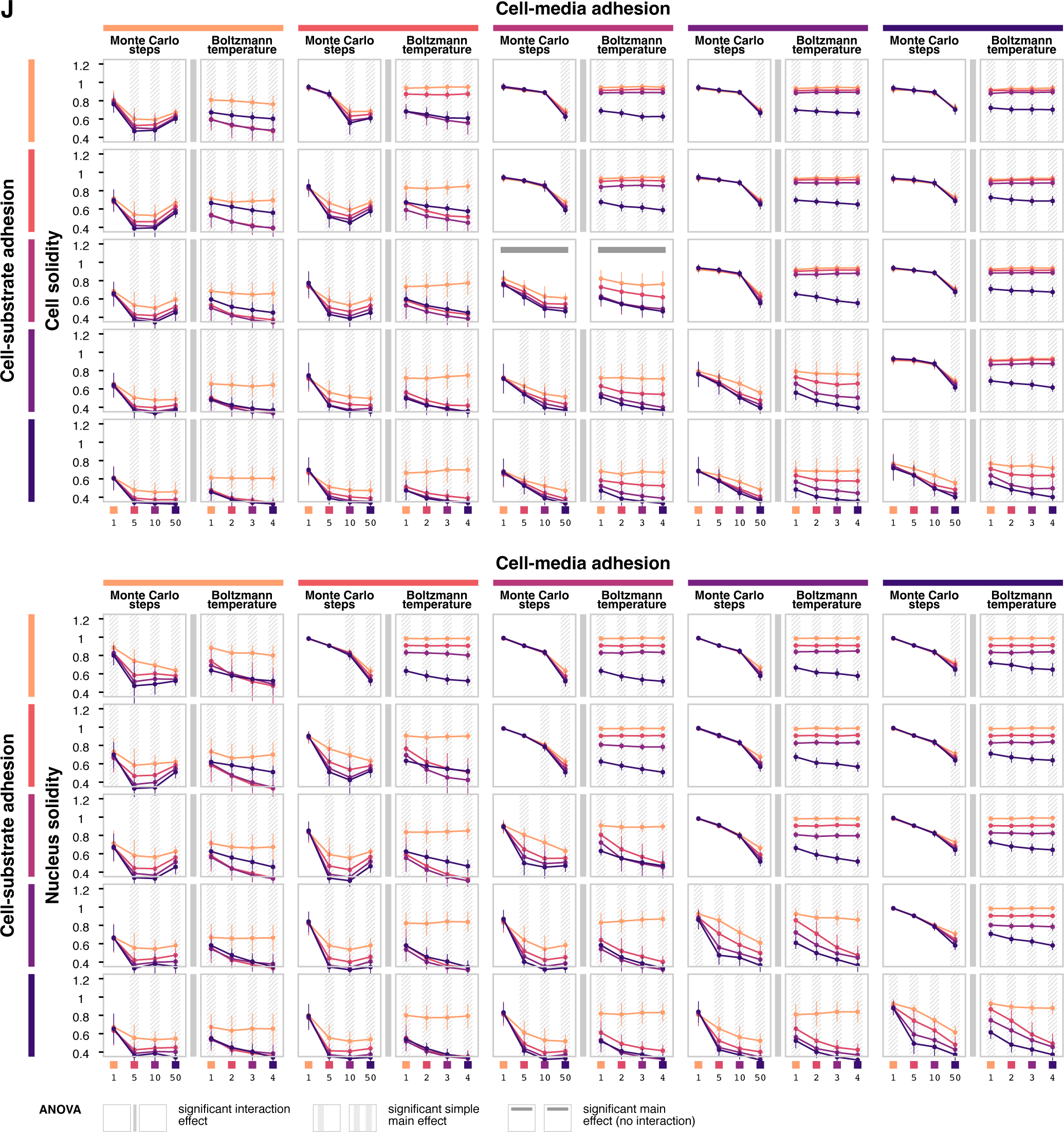
Framework-specific parameter simulations. *Related to Figure 4*. ANOVA interval plots for cell and nuclear (**A**) volume, (**B**) height, (**C**) area, (**D**) perimeter, (**E**) eccentricity, (**F**) major axis length, (**G**) minor axis length, (**H**) orientation, (**I**) extent, and (**J**) solidity. ANOVA is performed separately for each combination of cell-media and cell-substrate adhesion. Each data point shows the mean value of the feature at *t* = 1 hour for the given level of Monte Carlo steps or temperature, holding the other condition constant. Error bars show standard deviation across *n* = 10 replicates. Annotations indicate statistically significant interaction effects, simple main effects (if interaction is significant) and main effects (if interaction is not significant) at *α* = 0.05.

**Figure S4.**
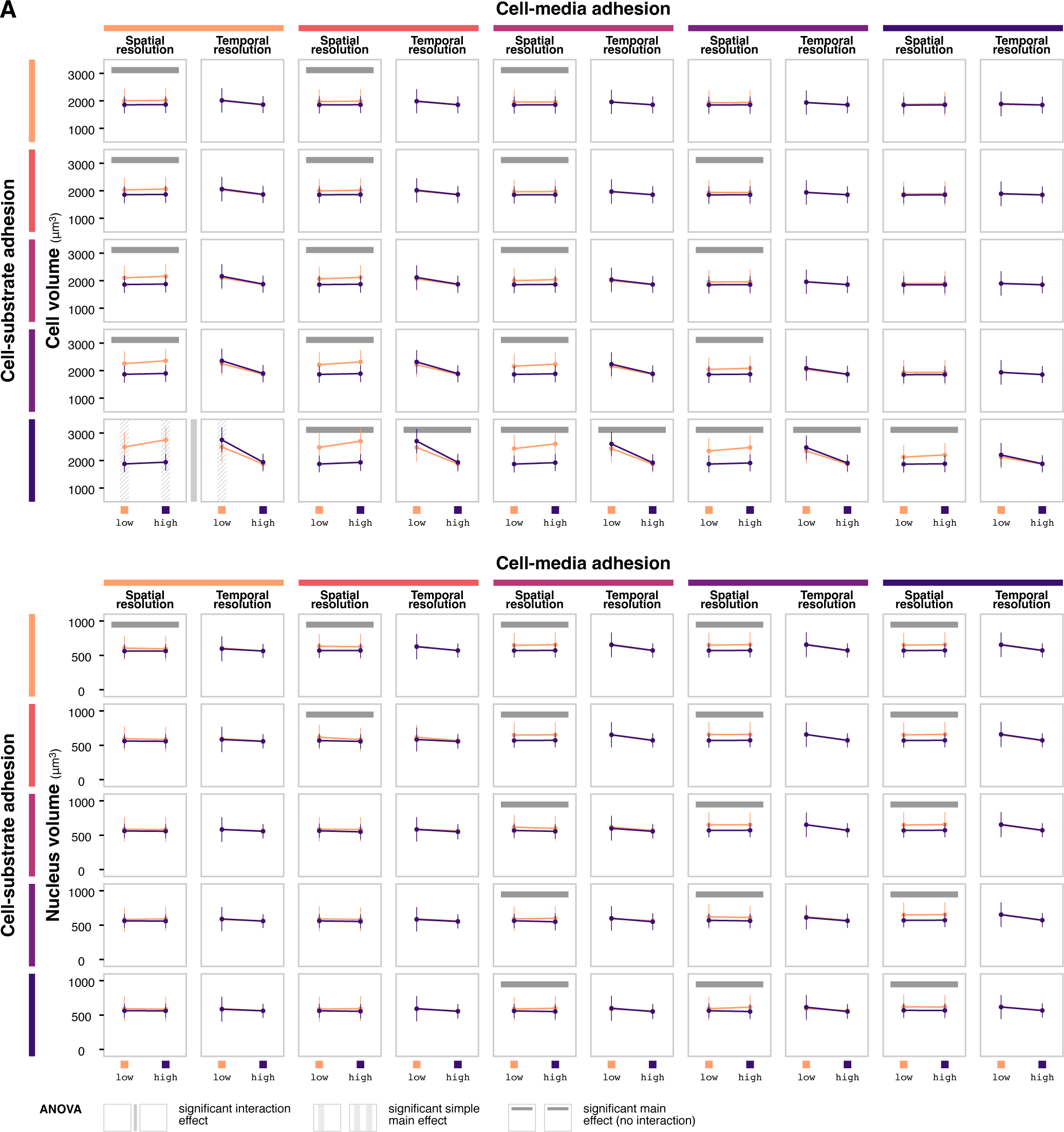
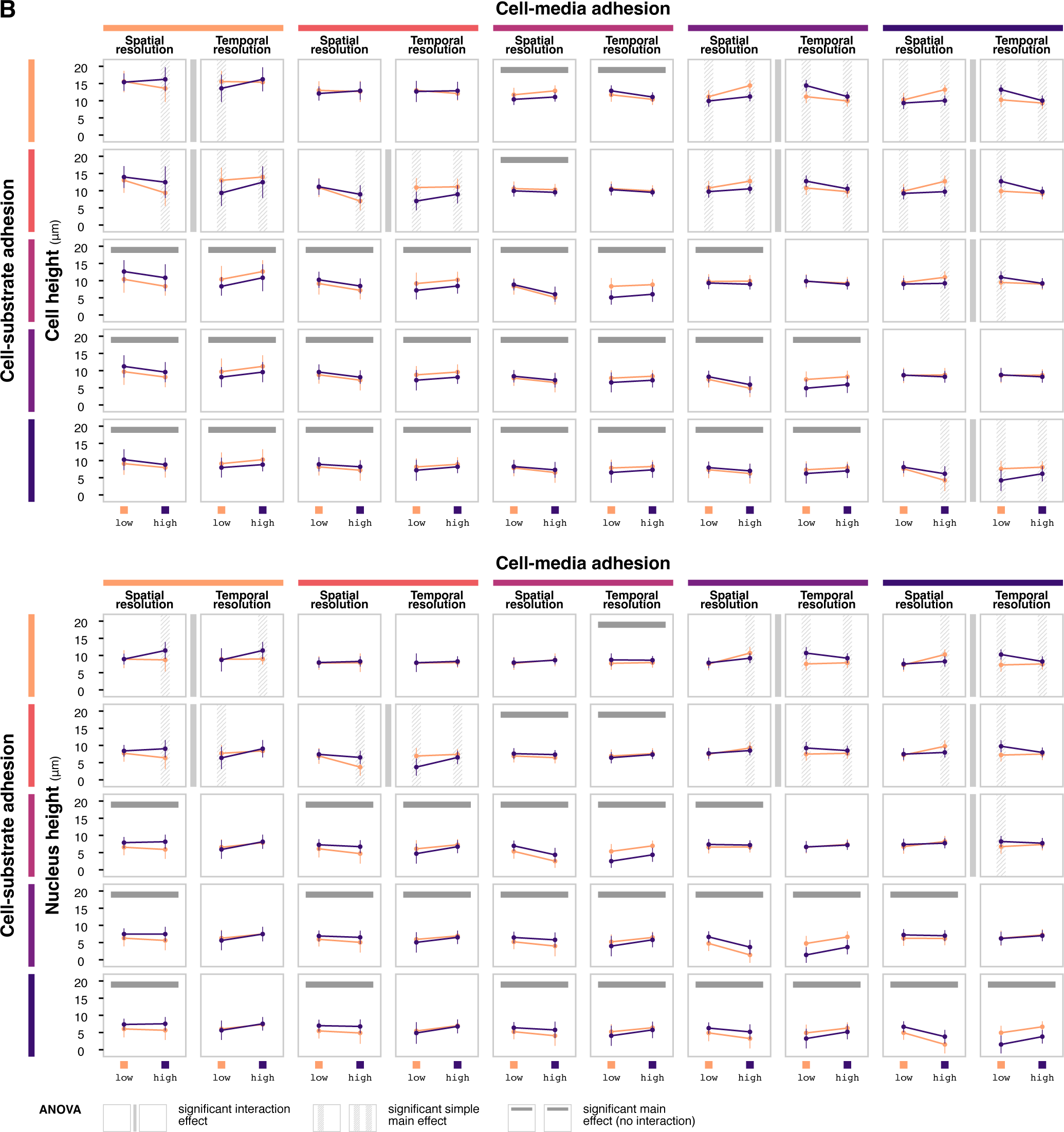
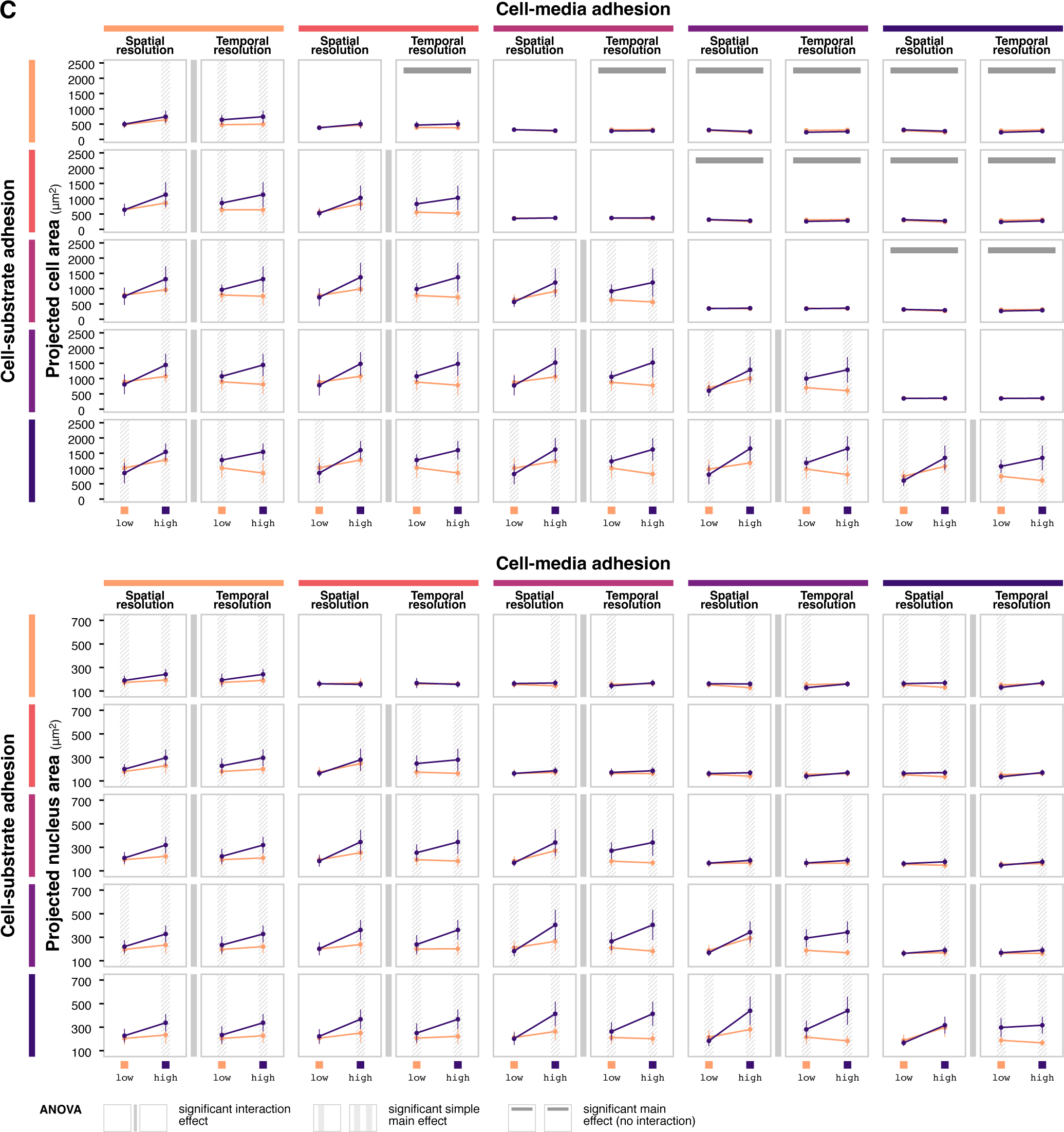
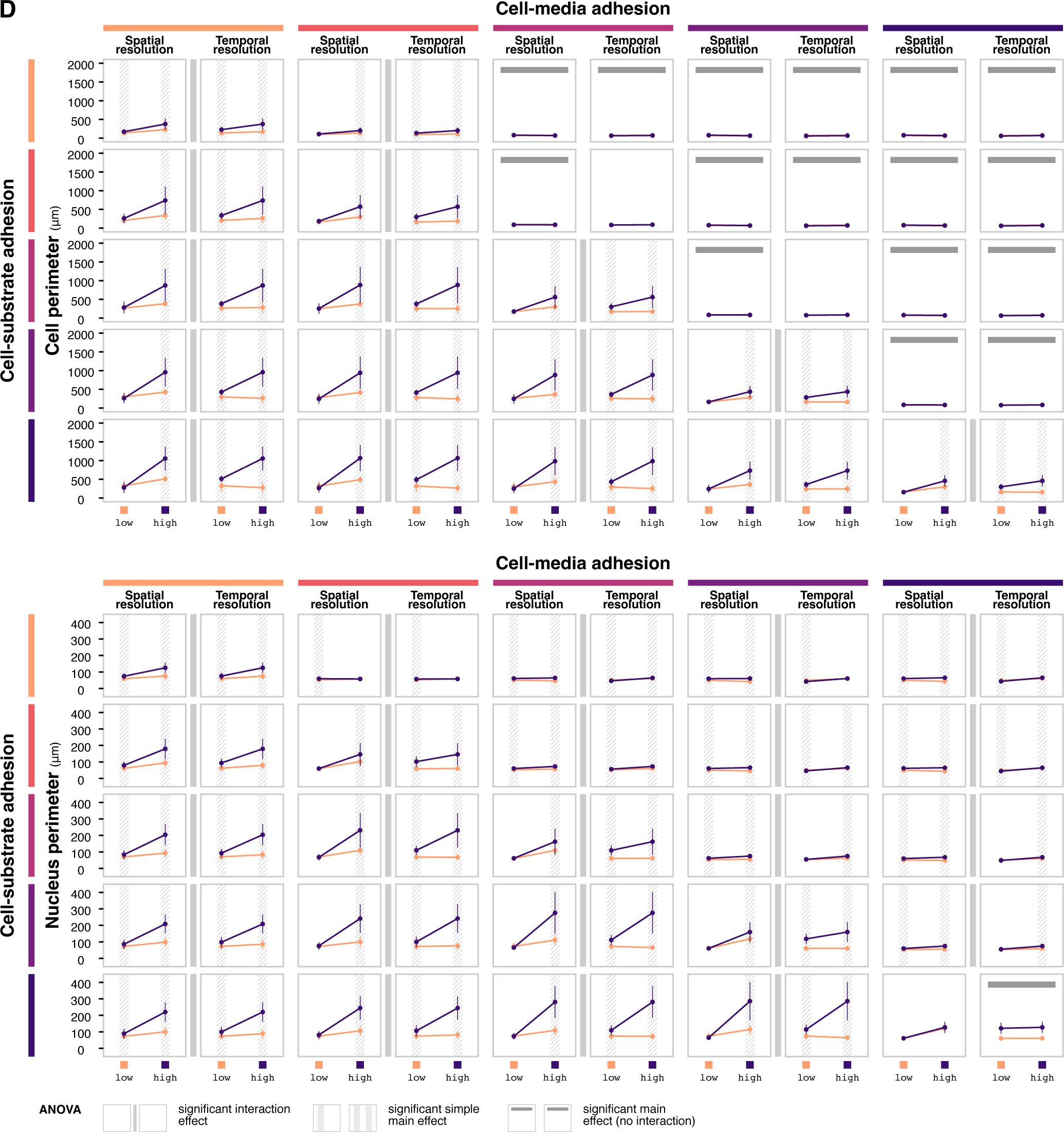
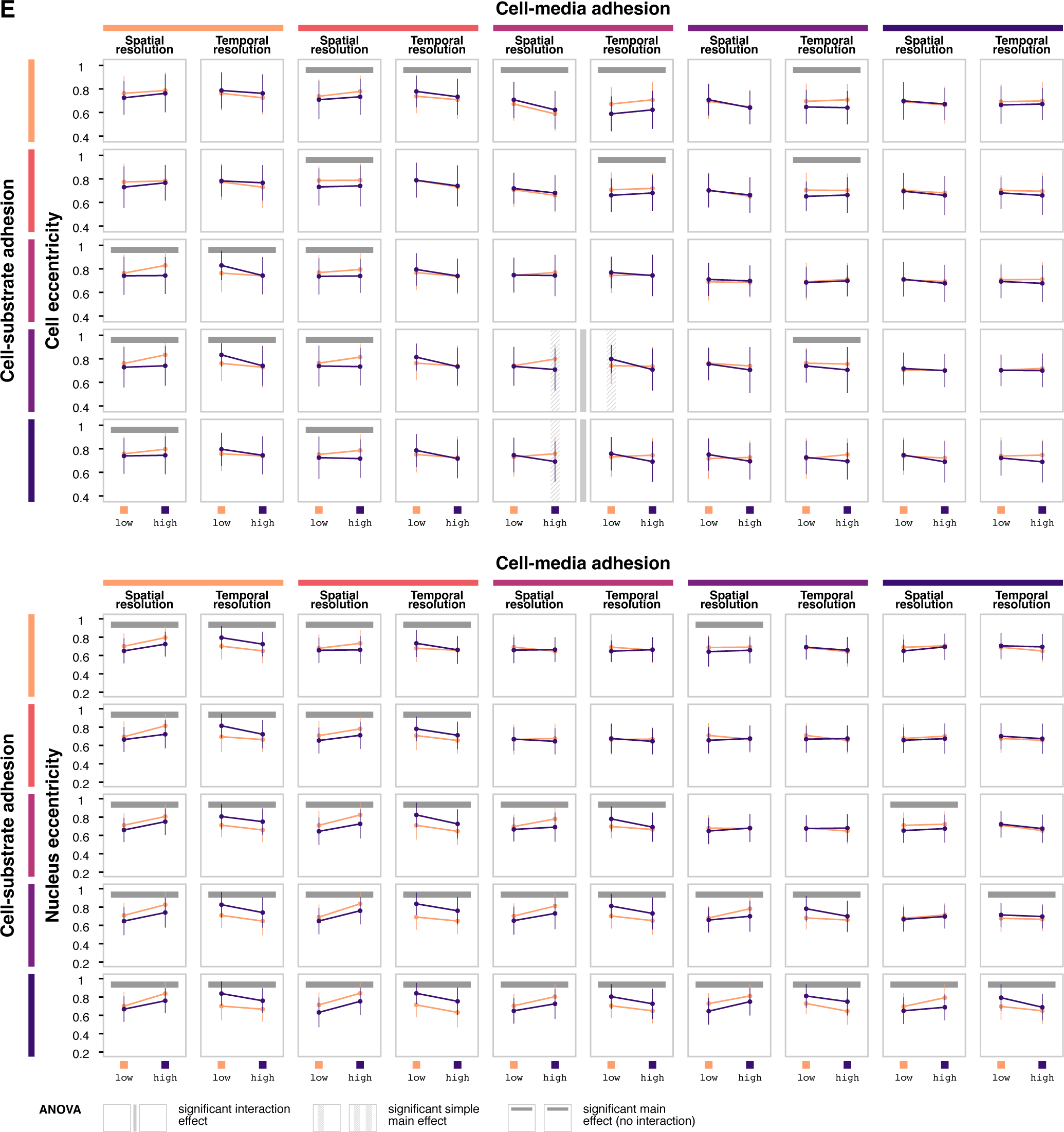
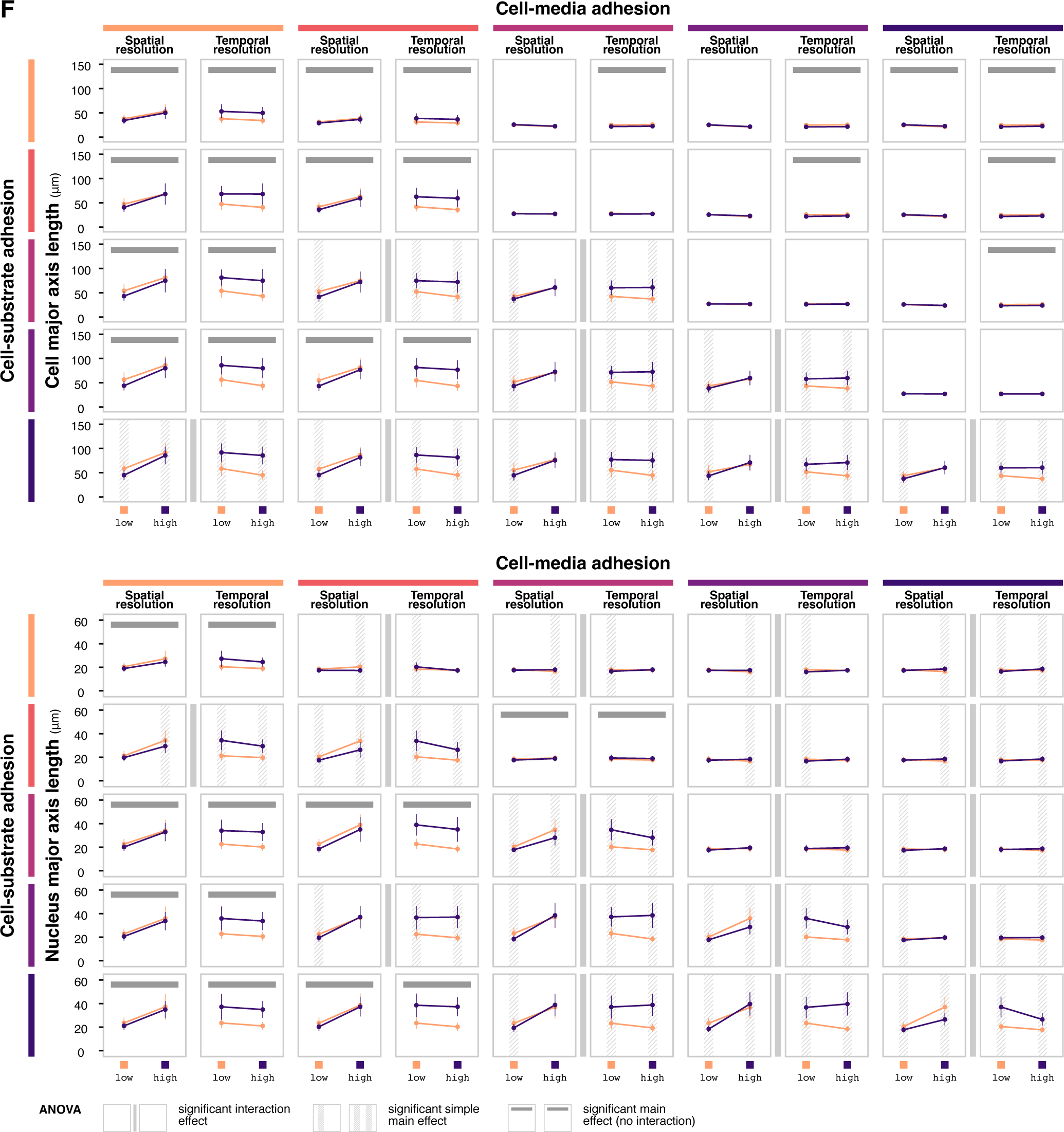
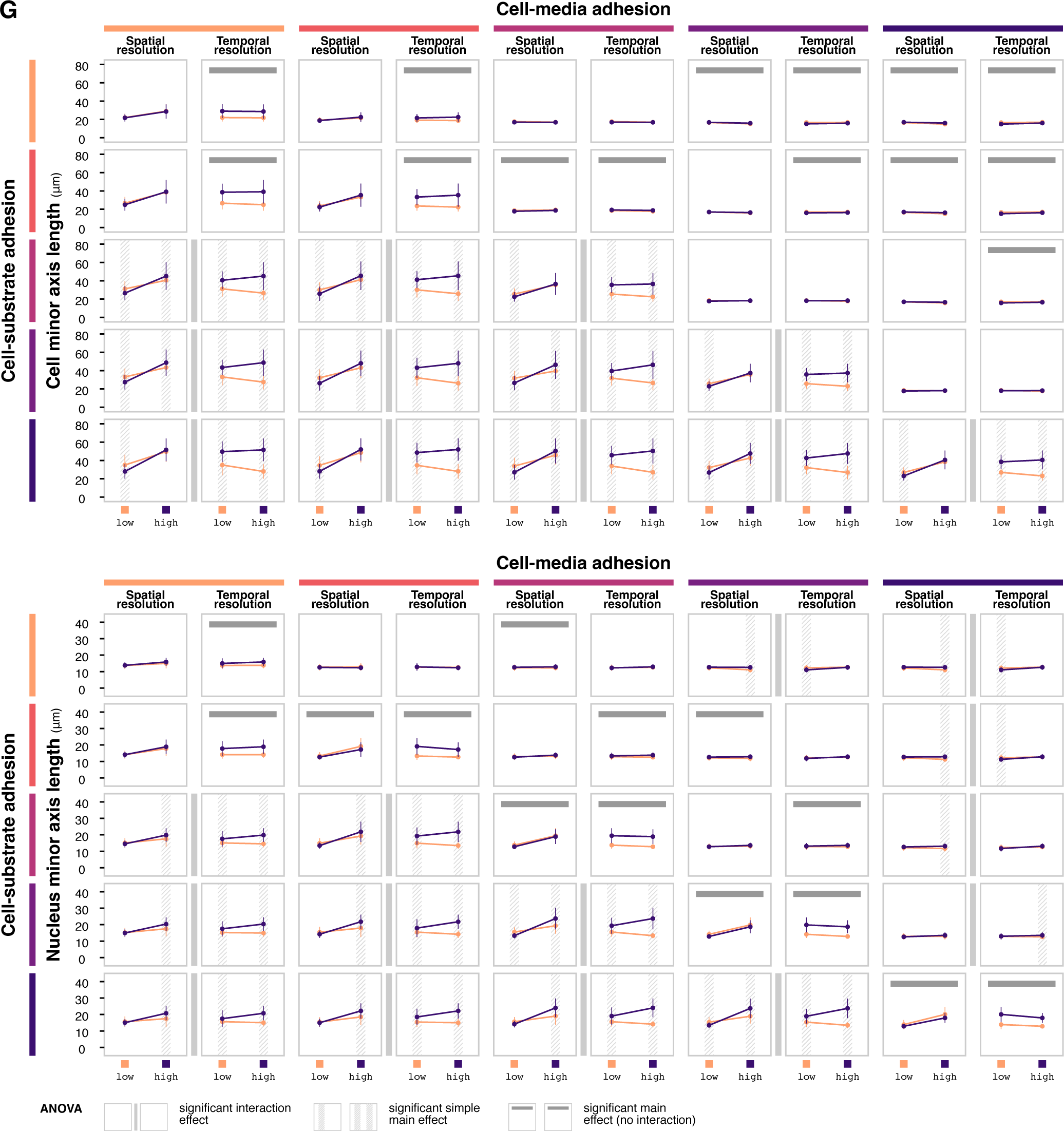
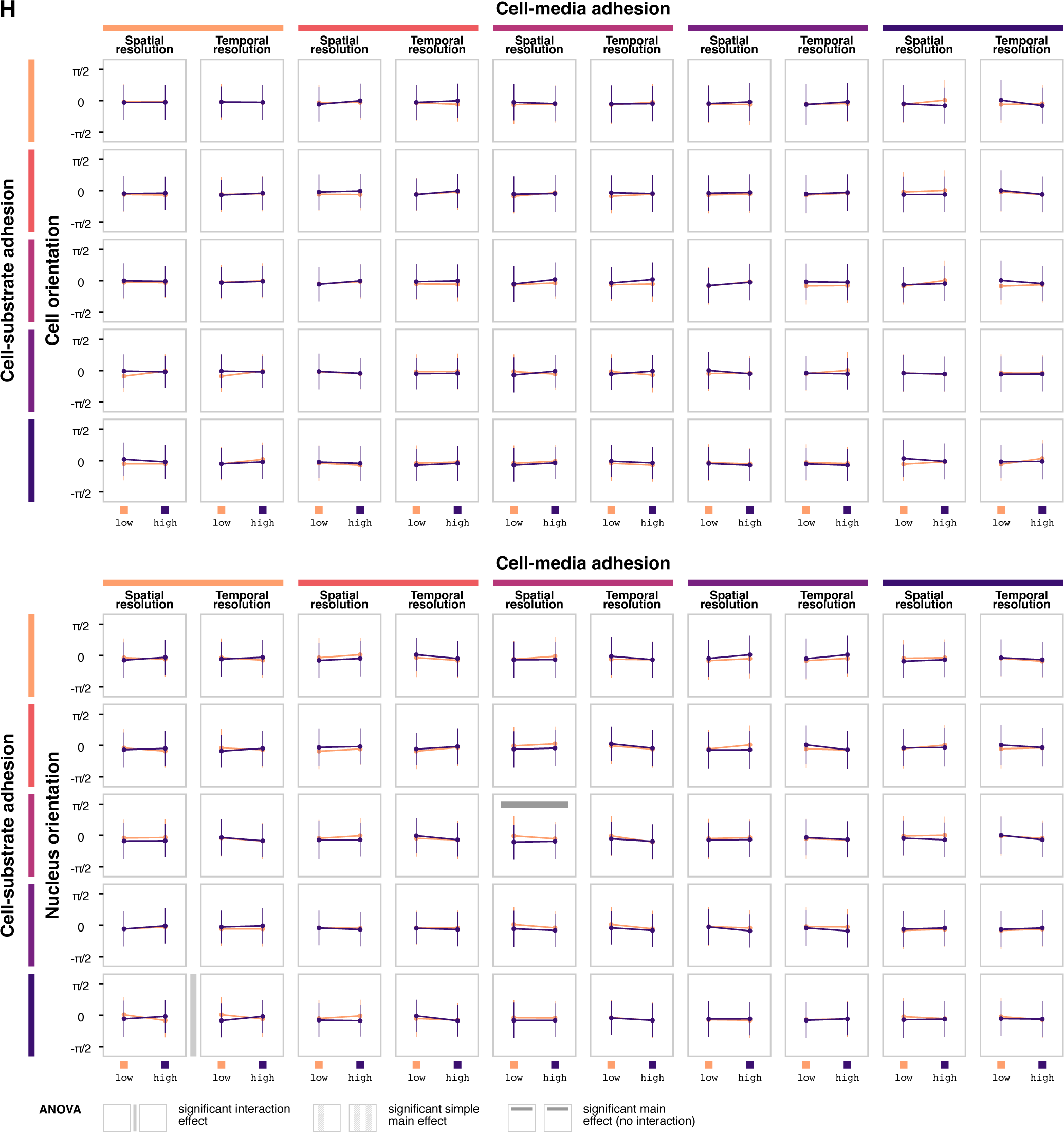
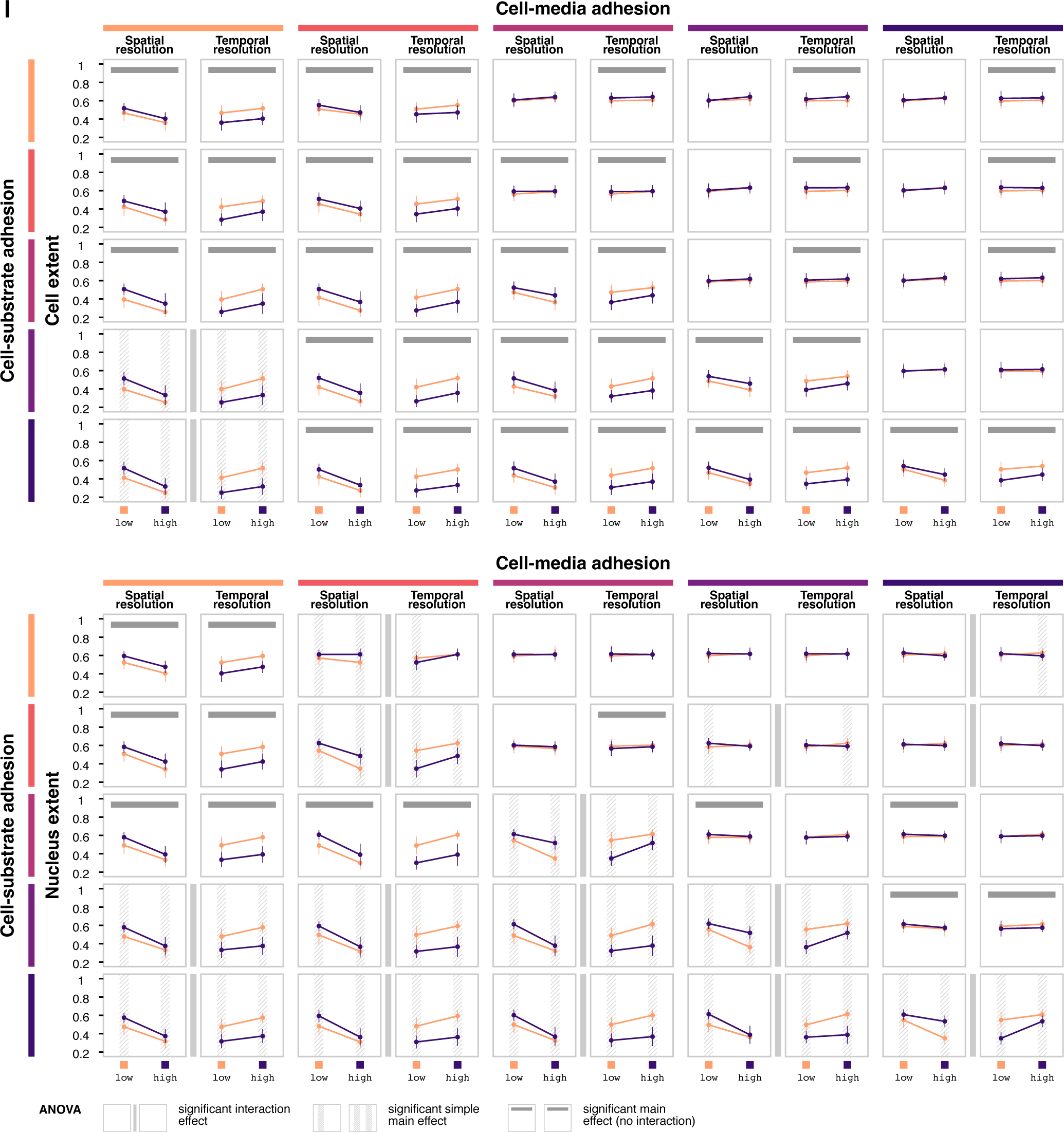
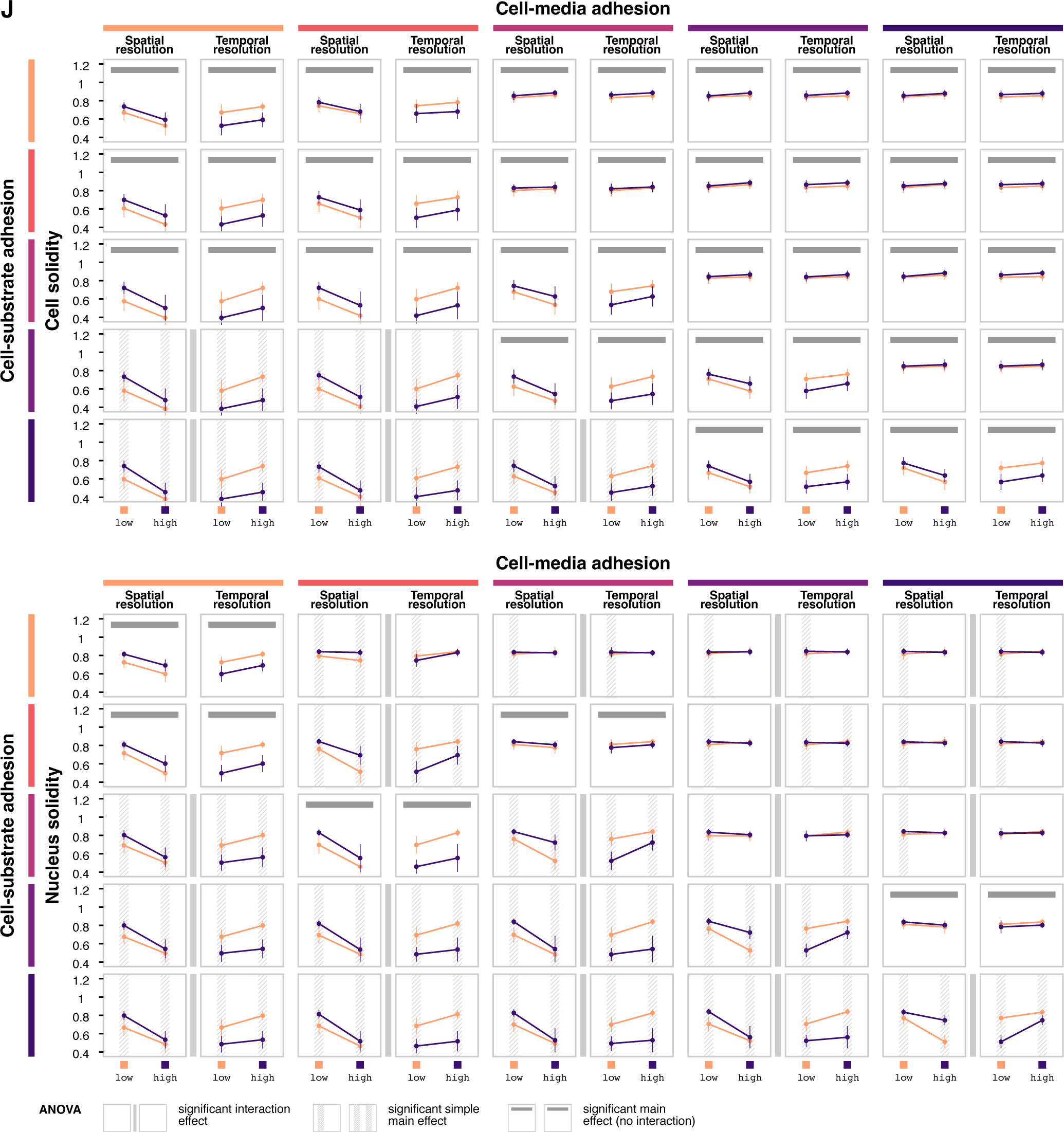
Spatial and temporal resolution parameter simulations. *Related to Figure 5*. ANOVA interval plots for cell and nuclear (**A**) volume, (**B**) height, (**C**) area, (**D**) perimeter, (**E**) eccentricity, (**F**) major axis length, (**G**) minor axis length, (**H**) orientation, (**I**) extent, and (**J**) solidity. ANOVA is performed separately for each combination of cell-media and cell-substrate adhesion. Each data point shows the mean value of the feature at *t* = 1 hour for the given level of spatial or temporal resolution, holding the other condition constant. Error bars show standard deviation across *n* = 10 replicates. Annotations indicate statistically significant interaction effects, simple main effects (if interaction is significant) and main effects (if interaction is not significant) at *α* = 0.05.

**Figure S5.**
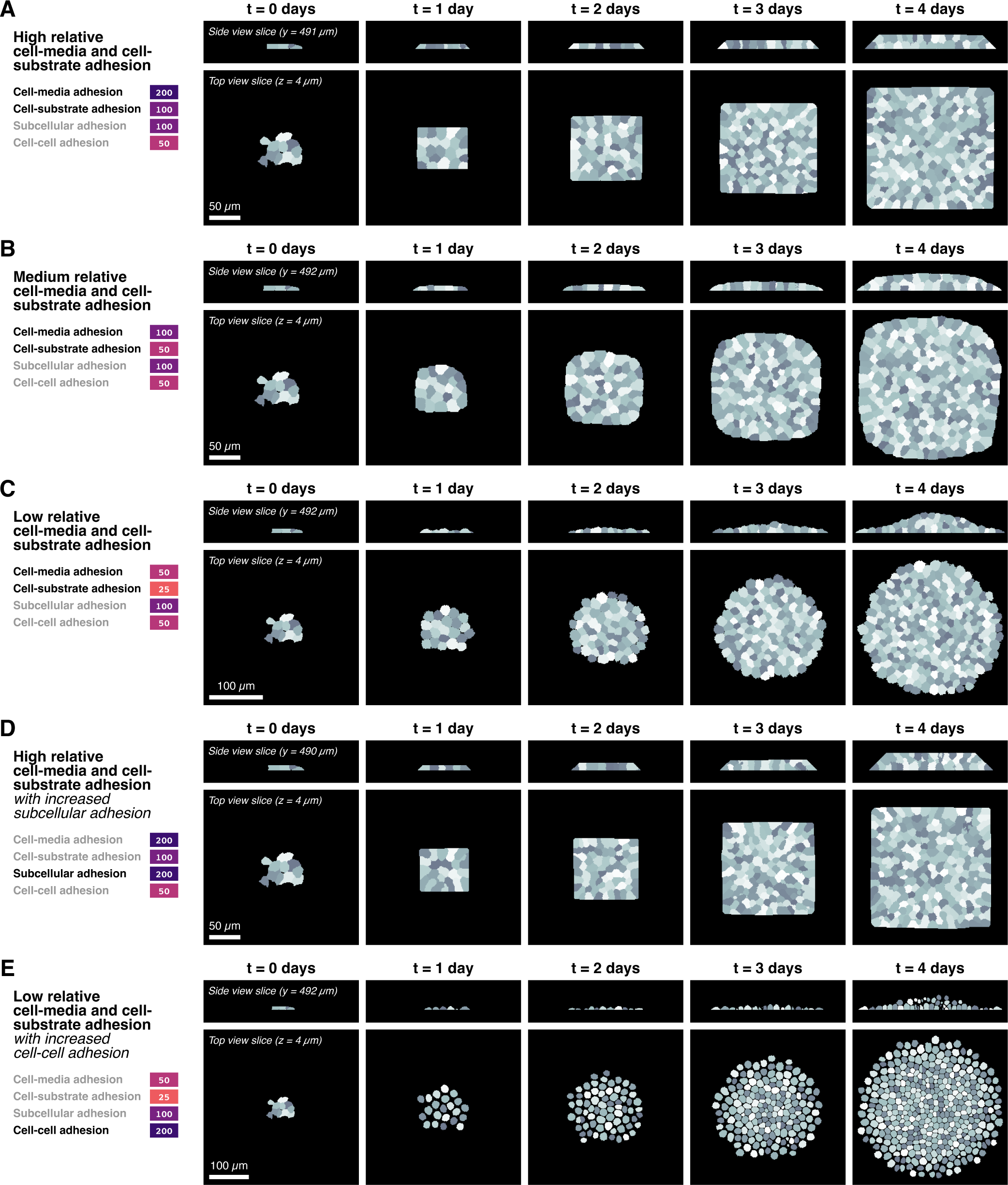
Full simulations for selected adhesion parameter conditions. *Related to Figure 6*. Cell and colony shapes for select time points across replicates for relative adhesion condition (**A**) high relative cell-media and cell-substrate adhesion, (**B**) medium relative cell-media and cell-substrate adhesion, (**E**) low relative cell-media and cell-substrate adhesion, (**D**) high relative cell-media and cell-substrate adhesion with increased subcellular adhesion, and (**C**) low relative cell-media and cell-substrate adhesion with increased cell-cell adhesion.

**Table S1.**
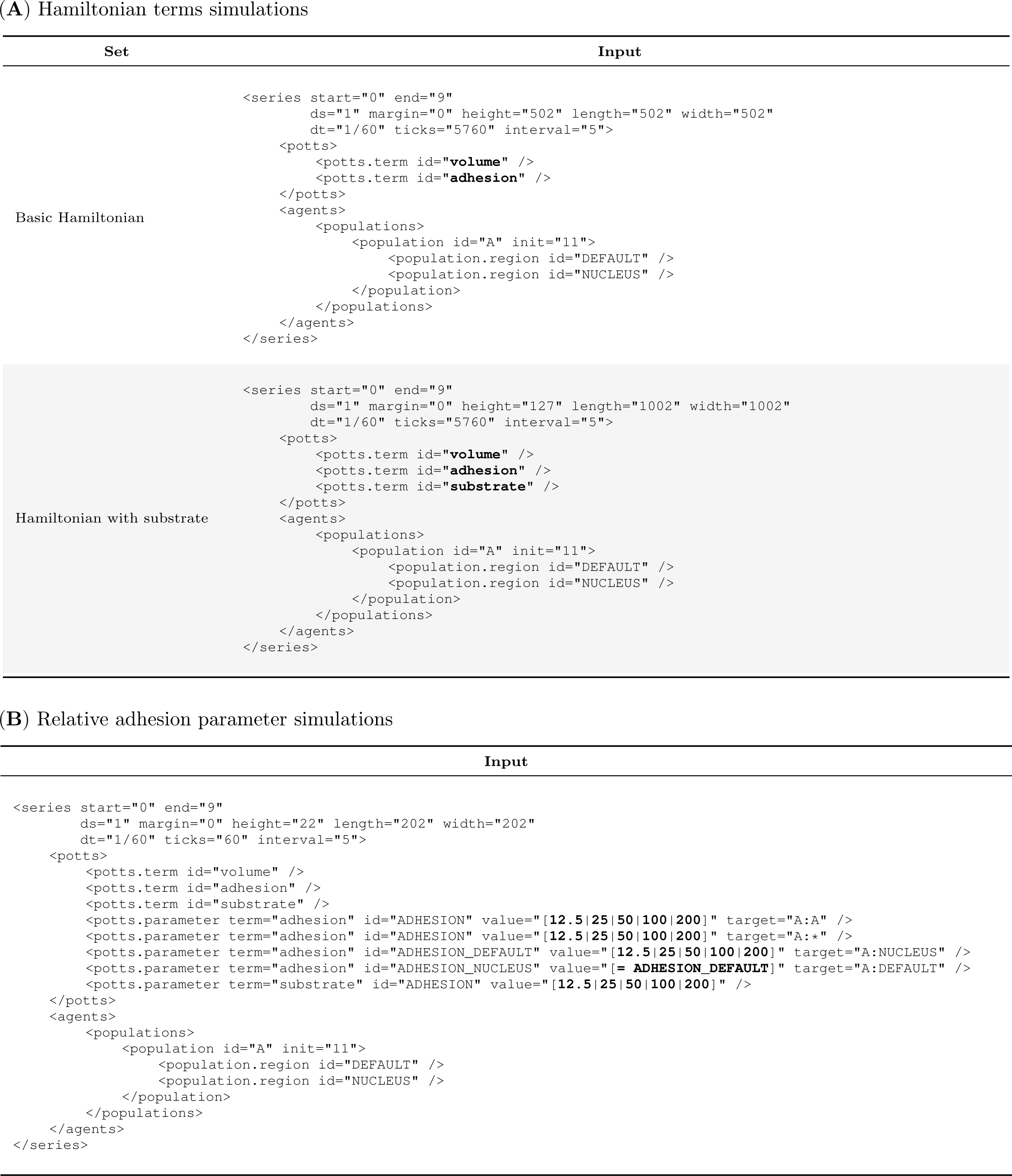
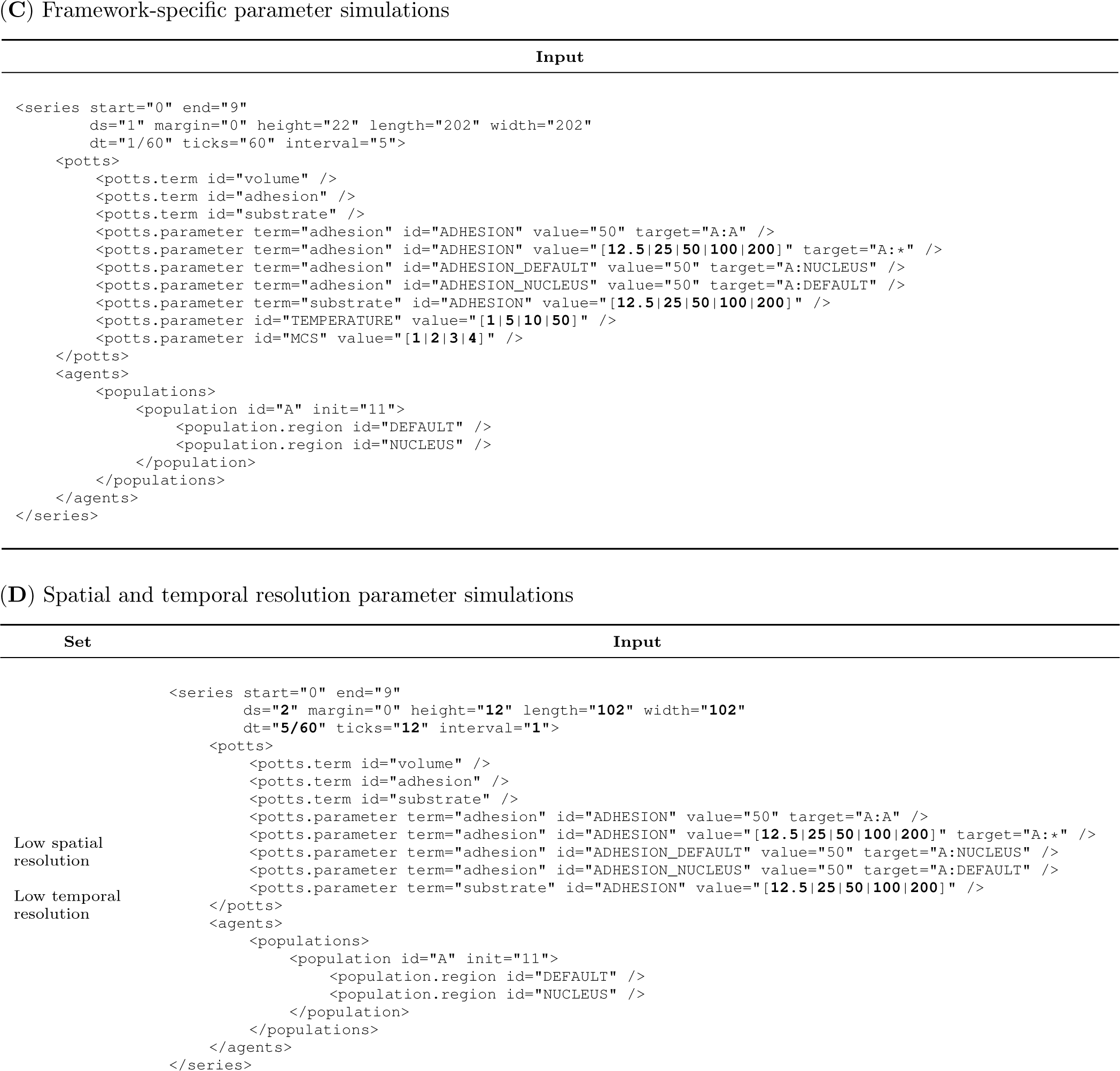
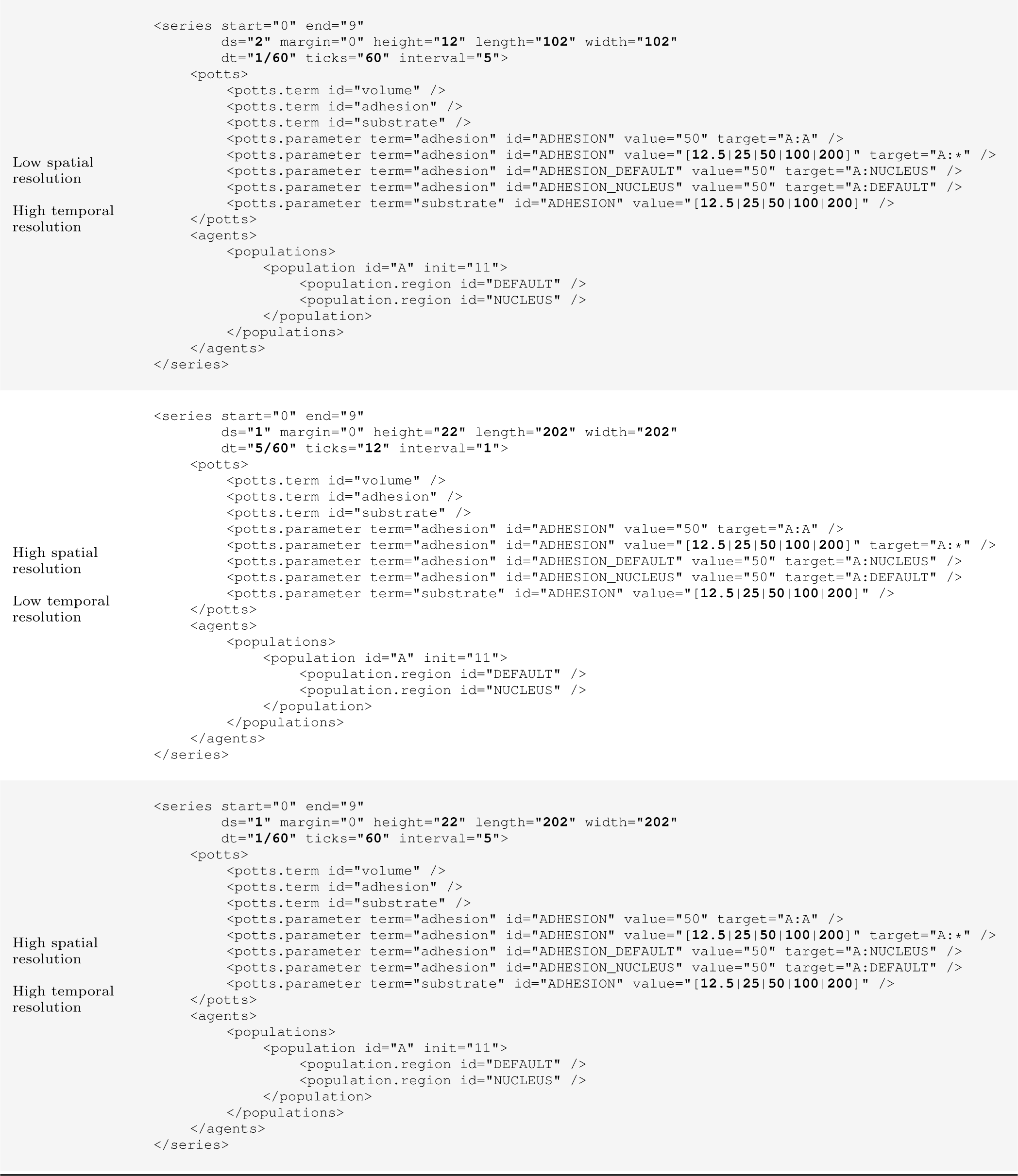
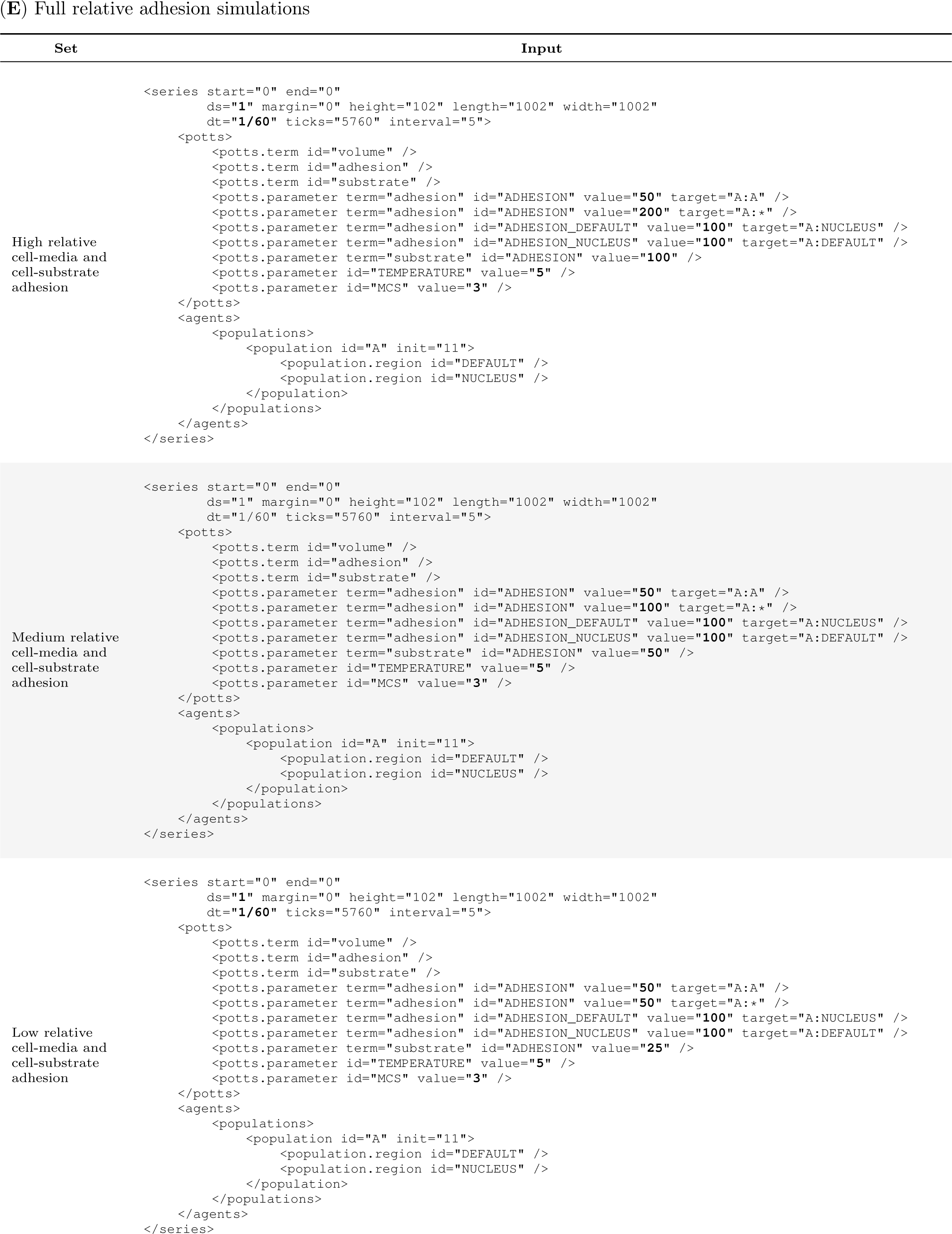
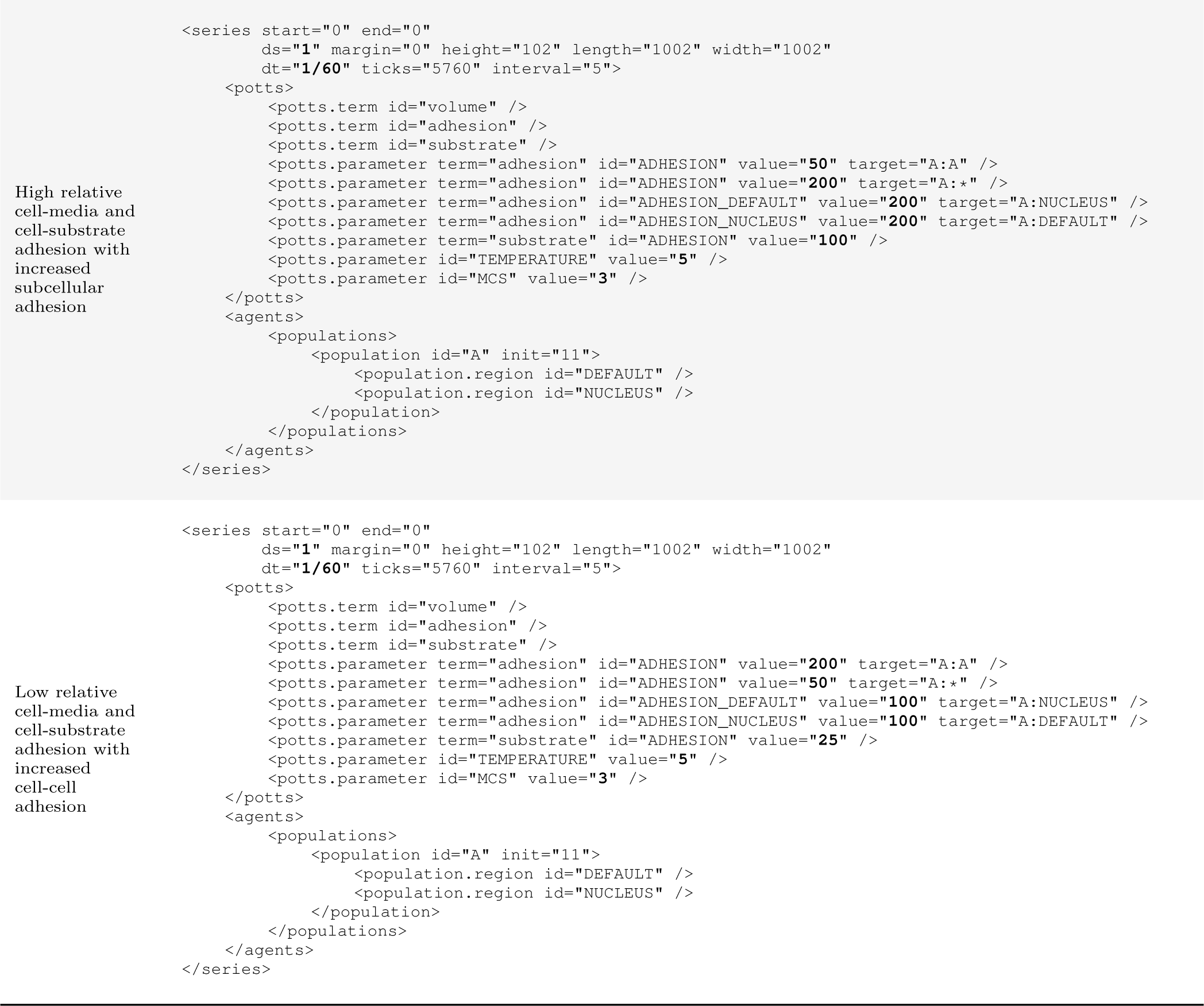
Input options used to run simulations. For clarity, wrapping `<SET>` tags and `<SERIES>` name attributes are not shown. For each set, simulations were run for every combination of bold options grouped by square brackets and separated by pipes.

**Table S2.** Wall clock time for simulations. Simulations are run on AWS EC2 using `m5.large` (2 vCPU and 16 GiB RAM) and `m5.xlarge` (4 vCPU and 32 GiB RAM) instances.

(A) Hamiltonian terms simulations
| Level | Time (hr) | N |
| --- | --- | --- |
| Basic Hamiltonian | 214.2 ± 10.6 | 10 |
| Hamiltonian with substrate | 214.5 ± 14.6 | 10 |

(B) Relative adhesion parameter simulations
| Parameter | Level | Time (min) | N |
| --- | --- | --- | --- |
| Cell-media adhesion | ● 12.5 | 2.8 ± 1.3 | 1250 |
| Cell-media adhesion | ● 25 | 2.5 ± 1.1 | 1250 |
| Cell-media adhesion | ● 50 | 2.1 ± 0.8 | 1250 |
| Cell-media adhesion | ● 100 | 1.7 ± 0.7 | 1250 |
| Cell-media adhesion | ● 200 | 1.7 ± 0.7 | 1250 |
| Cell-substrate adhesion | ● 12.5 | 2.3 ± 1.1 | 1250 |
| Cell-substrate adhesion | ● 25 | 2.3 ± 1.1 | 1250 |
| Cell-substrate adhesion | ● 50 | 2.2 ± 1.0 | 1250 |
| Cell-substrate adhesion | ● 100 | 2.0 ± 1.0 | 1250 |
| Cell-substrate adhesion | ● 200 | 1.9 ± 0.9 | 1250 |
| Cell-cell adhesion | ● 12.5 | 2.2 ± 1.0 | 1250 |
| Cell-cell adhesion | ● 25 | 2.2 ± 1.0 | 1250 |
| Cell-cell adhesion | ● 50 | 2.1 ± 1.0 | 1250 |
| Cell-cell adhesion | ● 100 | 2.2 ± 1.1 | 1250 |
| Cell-cell adhesion | ● 200 | 2.1 ± 1.1 | 1250 |
| Subcellular adhesion | ● 12.5 | 2.4 ± 1.1 | 1250 |
| Subcellular adhesion | ● 25 | 2.4 ± 1.1 | 1250 |
| Subcellular adhesion | ● 50 | 2.3 ± 1.0 | 1250 |
| Subcellular adhesion | ● 100 | 2.0 ± 1.0 | 1250 |
| Subcellular adhesion | ● 200 | 1.8 ± 0.9 | 1250 |

(C) Framework-specific parameter simulations
| Parameter | Level | Time (min) |  |  | N |
| --- | --- | --- | --- | --- | --- |
| Cell-media adhesion | ● 12.5 | 5.6 | ± | 4.2 | 800 |
| Cell-media adhesion | ● 25 | 5.2 | ± | 4.1 | 800 |
| Cell-media adhesion | ● 50 | 4.5 | ± | 3.8 | 800 |
| Cell-media adhesion | ● 100 | 3.9 | ± | 3.5 | 800 |
| Cell-media adhesion | ● 200 | 4.0 | ± | 3.6 | 800 |
| Cell-substrate adhesion | ● 12.5 | 5.0 | ± | 4.3 | 800 |
| Cell-substrate adhesion | ● 25 | 4.9 | ± | 4.1 | 800 |
| Cell-substrate adhesion | ● 50 | 4.8 | ± | 4.0 | 800 |
| Cell-substrate adhesion | ● 100 | 4.4 | ± | 3.6 | 800 |
| Cell-substrate adhesion | ● 200 | 4.2 | ± | 3.4 | 800 |
| Monte Carlo steps | ● 1 | 1.9 | ± | 1.3 | 1000 |
| Monte Carlo steps | ● 2 | 3.7 | ± | 2.5 | 1000 |
| Monte Carlo steps | ● 3 | 5.5 | ± | 3.6 | 1000 |
| Monte Carlo steps | ● 4 | 7.4 | ± | 4.8 | 1000 |
| Boltzmann temperature | ● 1 | 1.3 | ± | 0.9 | 1000 |
| Boltzmann temperature | ● 5 | 3.9 | ± | 2.7 | 1000 |
| Boltzmann temperature | ● 10 | 5.8 | ± | 3.6 | 1000 |
| Boltzmann temperature | ● 50 | 7.6 | ± | 4.3 | 1000 |

(D) Spatial and temporal resolution parameter simulations
| Parameter | Level | Time (sec) |  |  | N |
| --- | --- | --- | --- | --- | --- |
| Cell-media adhesion | ● 12.5 | 52.6 | ± | 80.0 | 200 |
| Cell-media adhesion | ● 25 | 47.0 | ± | 72.1 | 200 |
| Cell-media adhesion | ● 50 | 39.0 | ± | 54.2 | 200 |
| Cell-media adhesion | ● 100 | 34.8 | ± | 48.7 | 200 |
| Cell-media adhesion | ● 200 | 32.9 | ± | 46.2 | 200 |
| Cell-substrate adhesion | ● 12.5 | 45.9 | ± | 67.3 | 200 |
| Cell-substrate adhesion | ● 25 | 45.3 | ± | 69.7 | 200 |
| Cell-substrate adhesion | ● 50 | 42.7 | ± | 65.8 | 200 |
| Cell-substrate adhesion | ● 100 | 38.0 | ± | 55.1 | 200 |
| Cell-substrate adhesion | ● 200 | 34.5 | ± | 49.8 | 200 |
| Spatial resolution | ● high | 79.9 | ± | 68.7 | 500 |
| Spatial resolution | ● low | 2.7 | ± | 1.8 | 500 |
| Temporal resolution | ● high | 68.3 | ± | 77.4 | 500 |
| Temporal resolution | ● low | 14.2 | ± | 15.8 | 500 |

(E) Full relative adhesion simulations
| Level | Time (hr) | N |
| --- | --- | --- |
| Low relative cell-media and cell-substrate adhesion | 412.7 | 1 |
| Medium relative cell-media and cell-substrate adhesion | 406.4 | 1 |
| High relative cell-media and cell-substrate adhesion | 411.7 | 1 |
| High relative cell-media and cell-substrate adhesion with increased subcellular adhesion | 403.7 | 1 |
| Low relative cell-media and cell-substrate adhesion with increased cell-cell adhesion | 393.5 | 1 |

## Notes

### Competing Interest Statement

The authors have declared no competing interest.

